# Volatile but persistent co-existence of self-compatibility and self-incompatibility in plants

**DOI:** 10.1101/2025.11.30.691363

**Authors:** Amit Jangid, Ohad Noy Feldheim, Tamar Friedlander

## Abstract

Plants employ diverse mating strategies, including self- and cross-fertilization or their combination. Despite the ubiquity of mixed mating – the use of both self- and cross-fertilization within the same population – its evolutionary origins and dynamic stability remain elusive. Here, we study self-incompatibility, in which self-fertilization is disabled by molecular recognition and enabled when mutations disrupt this recognition. Using population-level stochastic simulations, we model the effects of two main parameters on the prevailing mating mode: promiscuity of molecular recognition and inbreeding depression, which penalizes self-compatibility. We reveal a phase diagram with three phases: complete self-incompatibility, complete self-compatibility, and a mixture combining both. The mixed phase exhibits newly-described vigorous, non-decaying fluctuations in the proportions of individuals occupying either mating mode. Pollen limitation extends the parameter range over which complete self-incompatibility is favored, contrary to the commonly held view. This study offers new insight into the evolutionary dynamics of plant reproductive systems.

## Introduction

Flowering plants are characterized by striking variation in their reproductive systems, facilitating different mating strategies including self-fertilization, cross-fertilization, and their mixture [1, 2]. Considerable theoretical work evaluated the benefits and costs of self-fertilization relative to cross-fertilization. A self-fertilizing parent enjoys a twofold transmission advantage as the contributor of two parental gametes to the offspring, compared to just one gamete under biparental reproduction [3, 4, 5, 6]. Self-fertilizing plants also enjoy the benefits of reproductive assurance, which is advantageous under mate limitation, as might happen in the population margins [5, 7], or when colonizing new territories [8, 9]. Yet, offspring produced by self-fertilization could suffer from various genetic defects, and a large proportion of them might be inviable, a phenomenon known as ‘inbreeding depression’ [10, 11].

Evolutionary transitions between mating modes are thought to occur frequently, as suggested by the prevalence of differences in mating systems within families and among closely related species [12]. Moreover, for some species, the mating mode could vary widely between and even within populations (reviewed in [13, 14, 15]). In particular, ‘mixed mating’ – the use of both self- and cross-fertilization within the same plant population – is common. Yet, it remains controversial whether mixed mating in plants is evolutionarily stable or is just a transitory population state. Specifically, Lande and Schemske proposed that only ‘pure’ mating strategies – either primarily outcrossing or primarily selfing – are stable, while their mixture is only transient [16]. Several models for the evolution of mixed mating were later proposed, assuming, for example, temporal variation in factors selecting for the mating mode [17, 18]. Yet, these models rely on specific assumptions or are suitable only in special cases, and there is yet no consensus model explaining the prevalence of mixed mating in plants – see [13] for a review.

Here we focus on self-incompatibility (SI) – widespread genetic mechanisms in plants that ensure cross-fertilization [19, 20, 21, 22, 23, 24, 25, 26]. Species harboring self-incompatibility mechanisms are subdivided into multiple ‘mating types’. The pollen recipient plant then uses molecular recognition to reject its own pollen type and accept only pollen of other types. The type is encoded by a single highly diverse genomic locus (‘S-haplotype’) that encodes genes expressed in the male and female reproductive organs.

Self-incompatibility (SI) can break down via various mutations [27, 28], leading to self-compatible (SC) plants capable of self-fertilization. The transition from self-incompatibility to self-compatibility is considered one of the most frequent evolutionary transitions in flowering plants [29, 30, 31]. Analyses show that closely related species could differ in their mating mode, suggesting the occurrence of evolutionary transitions between SI and SC following speciation [32, 33, 12, 34]. Some natural populations exhibit a perplexing picture, with different populations of the same species exhibiting either SI, SC, or a mixture of the two [35, 36, 37, 38, 39]. Transitions between SI and SC mating modes were also demonstrated to be developmental [40] or depend on environmental conditions [28]. Transitions to self-compatibility (SC) were considered irreversible because they typically stimulate the purging of deleterious mutations and, consequently, reduce inbreeding depression, thereby decreasing the cost of self-fertilization. The transition is also often accompanied by the “selfing syndrome” – the reduction or loss of floral traits associated with insect pollination [24], diminishing the possibility of outcrossing and again hampering the reverse transition. The complexity of SI systems, which depend on the coordinated functionality of multiple genes, was thought to preclude their re-establishment once disrupted [41, 42]. Nevertheless, recent evidence indicates that, in certain cases, SC arises from transcriptional silencing of SI-related genes [28] rather than from mutations in their coding sequences, raising the possibility that transitions to SC may be reversible.

In addition to the prevalence of self-incompatibility, its tight linkage to plant fitness, on the one hand, and its fragility, on the other hand, call for a theoretical study of the dynamics and reversibility of transitions between SI and SC reproductive strategies. In particular, it remains unclear whether SC and SI co-existence in the same population (‘mixed mating’) is long-lived or whether the empirically observed mixed populations are essentially in transience. Recent climatic changes are thought to bias the frequencies of different plant mating systems in different ways [43, 44, 45], further underscoring the need to study the relationship between environmental factors and plant mating mode.

In a seminal paper, Charlesworth and Charlesworth [46] analyzed the balance between self-compatible and self-incompatible sub-populations at steady state and their stability to invasion, depending on the probability that offspring produced via self-fertilization are inviable (the ‘inbreeding depression’ value *δ*), the proportion of self-pollen received, and the number of distinct self-incompatible mating types. They found that a minimal inbreeding depression value (*δ >* 1*/*2) is needed for the invasion of a self-incompatible mutant into an otherwise self-compatible population. Complementarily, they found that a self-compatible mutant cannot invade a self-incompatible population whose number of types exceeds a critical value. In between, they found an intermediate parameter regime exhibiting an SI-SC co-existence with dynamically stable proportions [46]. Stable SI-SC co-existence was also obtained in additional models [47, 48, 49]. Yet, none found long-lived non-decaying variation in the SI-SC proportions.

Here, we study the most prevalent SI mechanisms - the collaborative non-self recognition self-incompatibility, demonstrated in several large plant families relevant to agriculture: the Solanaceae family (tomato, potato, tobacco, Petunia), Rosaceae (apple, pear, loquat), Rutaceae (citrus), Rubi-aceae (coffee), and Plantaginaceae (snapdragon) [50]. This mechanism is also known as the ‘RNase-based SI’ because the female component in this mechanism is a cytotoxic ribonuclease (RNase), and different mating types have different RNase alleles [51]. The male component in this mechanism (‘SLF’) is an F-box protein involved in targeted protein degradation [52, 53]. If the pollen is compatible, the maternal plant’s RNase molecules are recognized by the pollen’s SLF proteins and consequently degraded, allowing for fertilization. Otherwise, if the pollen is incompatible, the maternal plant’s RNase molecules arrest the pollen tube growth, and fertilization is inhibited. Multiple SLF genes are encoded in every S-haplotype to collaboratively recognize and detoxify various non-self RNases and allow for a sufficient number of mating partners [54]. Self-compatibility in this system could be achieved either via the gain of an SLF gene compatible with the RNase of the same S-haplotype [55, 56] or via inactivation or silencing of the RNase [37, 57, 28].

We have recently proposed a novel theoretical framework to study the evolution of this self-incompatibility mechanism [58], inspired by thermodynamic models of gene regulation [59, 60]. This model uniquely incorporates a biophysical representation of molecular recognition between male- and female-type-specifying proteins into the evolutionary model. An emergent property of this model is the spontaneous self-organization of the population into distinct mating types, as in natural populations, where the number of types emerges naturally, rather than being a preset parameter, as in previous models. Using this framework, we studied the evolutionary trajectories of mating type emergence and decay. This modeling framework crucially relies on the promiscuity of molecular recognition between the female and male type-specifying proteins, in agreement with empirical evidence [54, 55, 34]. The magnitude of this newly introduced parameter – the extent of promiscuity – turns out to be fundamental to our model’s behavior [61]. The model explicitly accounts for mutations that drive self-incompatible genotypes toward self-compatible ones and vice versa. The use of population-level stochastic simulations enables the study of its dynamical behavior, and in particular, the stability of the different mating modes to mutations. All these desired properties render the model highly suitable for studying the questions regarding dynamical transitions between mating modes stated above.

Here, we use this model to investigate the roles of two primary parameters — inbreeding depression and molecular recognition promiscuity — in shaping population mating modes and the number of SI mating types, through stochastic simulations. Our choice of stochastic simulations is dictated by the extremely high dimensionality of a genotype in our model, which incorporates multiple proteins, each represented by a sequence of amino acids, and by the fact that a finite population samples only a tiny fraction of this enormous genotype space, rendering the model unsusceptible to deterministic analysis. We find three population mating modes in different regions of the promiscuity-inbreeding depression parameter space. Apart from the full SI and full SC mating modes demonstrated in previous theoretical models [46, 16, 48, 49, 62, 63], we discover a third new “volatile mixed mode”. In the latter, both SC and SI sub-populations are found in dynamically unstable proportions, yet the population does not collapse into either a purely SI or a purely SC state, even after a very long time. This is the first time such an unstable yet long-lasting mixed mode has been theoretically predicted, in the absence of fluctuations in the model parameters. The two transitions between the mating modes show characteristics reminiscent of phase transitions in thermodynamic systems, with a smooth transition between the full SC and mixed phases and a sharp transition between the mixed and full SI phases.

We provide a full phase diagram for the model and analytically calculate the phase boundaries, finding excellent agreement with our simulation results. In the full SI regime, we find that the number of mating-specificities increases with interaction promiscuity. In the volatile mixed phase, we observe vigorous non-decaying fluctuations, not only in the SI population proportion but also in the number of mating-specificities. Transitions between SI to SC mating modes turn out to be reversible in a great part of the parameter space, as long as the inbreeding depression does not decrease below a threshold value.

## Results

### The model

We use here a slightly modified version of our previous model [58]. For completeness, we fully describe the model below. We consider a population of *N* diploid individuals, each composed of two S-haplotypes. Every S-haplotype includes a single RNase (encoding the female specificity) and multiple SLF paralogs (encoding the male specificity) – Fig. 1a, where the RNase could be either active or inactive. Every diploid individual plays the role of a diploid maternal plant, as well as the donor of two types of haploid pollen carrying either of its two S-haplotypes. A haploid pollen can successfully fertilize a diploid maternal plant only if it is equipped with matching SLF(s) that can detoxify the maternal plant’s two RNases – Fig. 1b, or alternatively if the maternal RNases are inactive. For simplicity, we construct the model directly in the protein domain. Each of these genes is then represented by a sequence of *L* amino acids, which could belong to either of four biochemical categories: hydrophobic H, neutral polar P, positively charged +, or negatively charged –. This AA sequence represents the protein interaction interface. An RNase-SLF pair is considered matching if the interaction energy between their interfaces is below a threshold value *E*_th_. This interaction energy between an RNase *R*_*i*_ and SLF *F*_*j*_ is defined as the sum of the pairwise interaction energies between the corresponding amino acids (see Table 1) of these proteins,

**Table 1.**
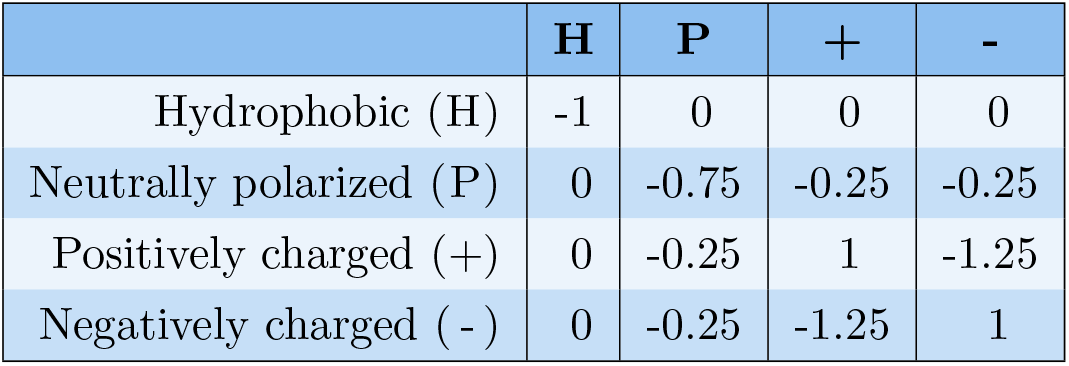
Interaction energies between pairs of amino acids.

**Table 2.**
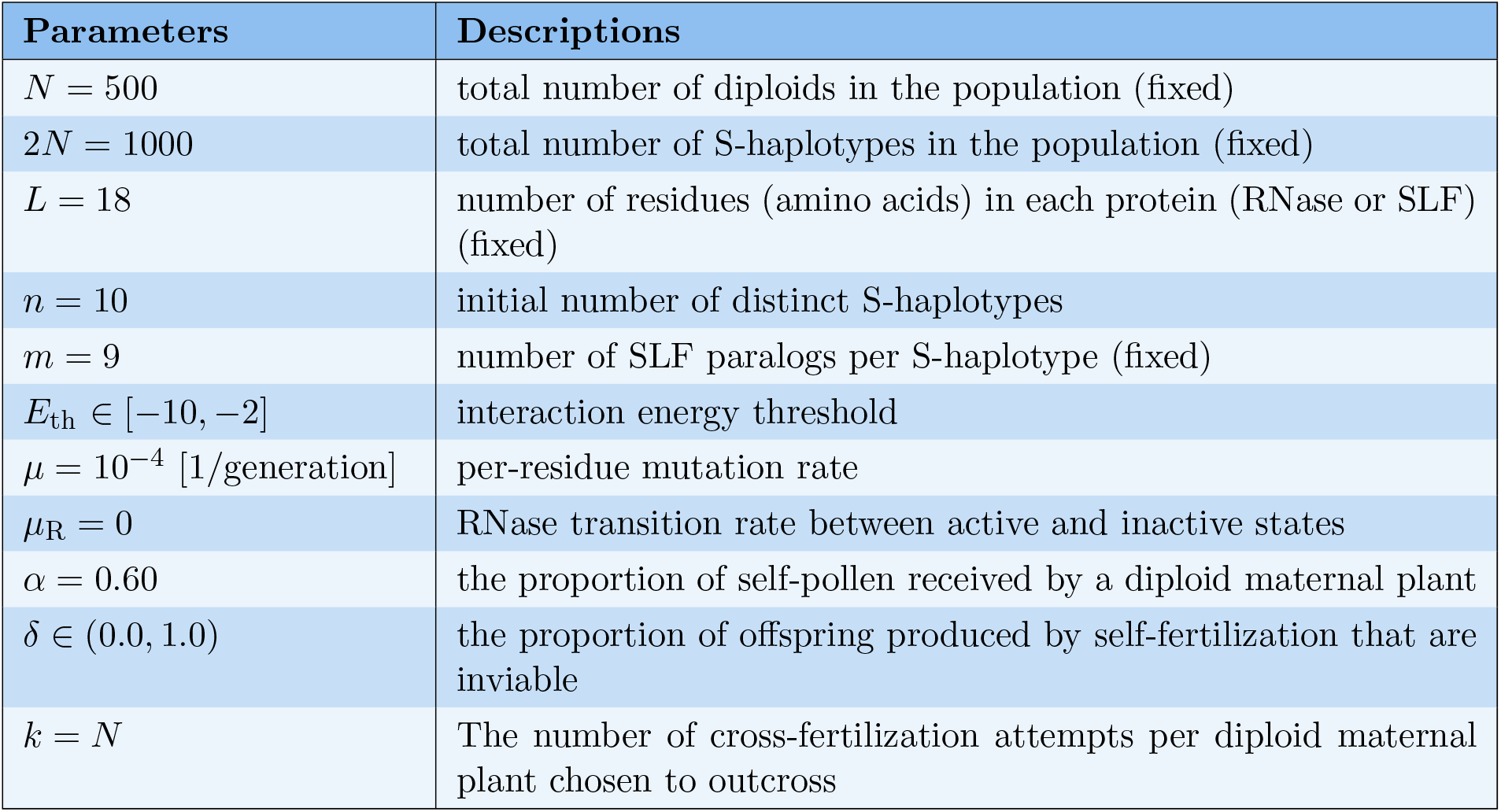
Description of parameters and default values used in the main model simulation.

**Table 3.** Female and male fitness of a single haplotype for different genotypes.

| Genotype | Female | Male |
| --- | --- | --- |
| FSC | $0.5 \frac{\alpha(1-\delta)}{1-\gamma_{\text{die}}} + 0.5 \frac{(1-\alpha)(1-(1-\frac{C_j}{2N})^k)}{1-\gamma_{\text{die}}}$ | $0.5 \frac{\alpha(1-\delta)}{1-\gamma_{\text{die}}} + \sum_{l=1; l \neq j}^N \left[ \frac{A_l}{1-\gamma_{\text{die}}} \frac{n_l^j}{C_l} \left( 1 - \left( 1 - \frac{C_l}{2N} \right)^k \right) \right]$ |
| HSC<sup>-</sup> | $0.5 \frac{\alpha'(1-\delta)}{1-\gamma_{\text{die}}} + 0.5 \frac{(1-\alpha')(1-(1-\frac{C_j}{2N})^k)}{1-\gamma_{\text{die}}}$ | $0 + \sum_{l=1; l \neq j}^N \left[ \frac{A_l}{1-\gamma_{\text{die}}} \frac{n_l^j}{C_l} \left( 1 - \left( 1 - \frac{C_l}{2N} \right)^k \right) \right]$ |
| HSC<sup>+</sup> | $0.5 \frac{\alpha'(1-\delta)}{1-\gamma_{\text{die}}} + 0.5 \frac{(1-\alpha')(1-(1-\frac{C_j}{2N})^k)}{1-\gamma_{\text{die}}}$ | $\frac{\alpha'(1-\delta)}{1-\gamma_{\text{die}}} + \sum_{l=1; l \neq j}^N \left[ \frac{A_l}{1-\gamma_{\text{die}}} \frac{n_l^j}{C_l} \left( 1 - \left( 1 - \frac{C_l}{2N} \right)^k \right) \right]$ |
| FSI | $0 + 0.5 \frac{(1-(1-\frac{C_j}{2N})^k)}{1-\gamma_{\text{die}}}$ | $0 + \sum_{l=1; l \neq j}^N \left[ \frac{A_l}{1-\gamma_{\text{die}}} \frac{n_l^j}{C_l} \left( 1 - \left( 1 - \frac{C_l}{2N} \right)^k \right) \right]$ |

whereas,
- $x$ : fraction of FSC female $y$ : fraction of HSC female $1 - x - y$ : fraction of FSI female $n_l^j$ : 1 if pollen $j$ is compatible to female $l$ else 0 $C_l$ : non-self compatible pollen to female $l$ $A_l$ : 1 - $\alpha$ if female $l$ is FSC $A_l$ : 1 - $\alpha'$ if female $l$ is HSC $A_l$ : 1 if female $l$ is FSI $F$ : set of indices of FSC females $H$ : set of indices of HSC females $I$ : set of indices of FSI females

**Figure 1.**
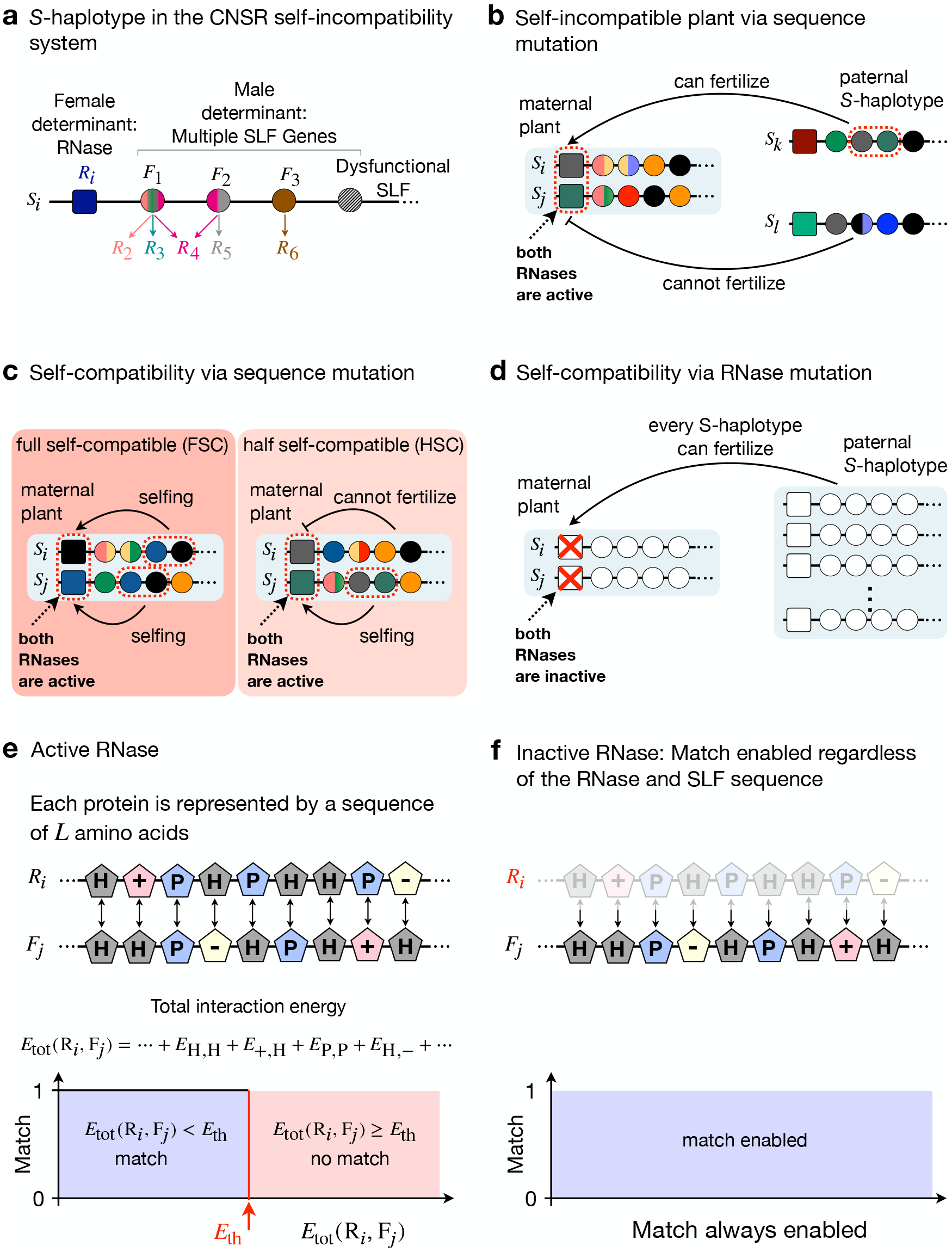
Model description. **(a)** Each S-haplotype consists of a single female determinant gene (RNase *R*_*i*_, square) and multiple male-determinant genes (SLFs *F*_1_, *F*_2_, …, circles). Each SLF could match one or more RNases, or none. **(b)** For a paternal S-haplotype to successfully fertilize a diploid maternal plant, it must be equipped with SLF(s) matching the two maternal RNases. An S-haplotype is considered ‘self-incompatible’ if it does not contain an SLF matching its own RNase, hence its fertilization by self-pollen is impossible. **(c-d)** An S-haplotype is considered ‘self-compatible’ either if it does contain an SLF matching its RNase and facilitating fertilization by self-pollen **(c)** or if its RNase is inactive **(d)**, in which case any S-haplotype can fertilize it, including the self one. **(e)** Each protein encoded in the S-haplotype is represented by a sequence of *L* amino acids of the four biochemical categories. The total interaction energy *E*_tot_ between an active RNase *R*_*i*_ and an SLF *F*_*j*_ is defined as the sum of the pairwise interaction energies between their corresponding amino acids. Two such proteins are considered matching if *E*_tot_ is smaller than an energy threshold *E*_th_. Otherwise, they are non-matching. **(f)** If the RNase is inactive, the maternal S-haplotype carrying it can be fertilized by any pollen, regardless of its SLF sequence.

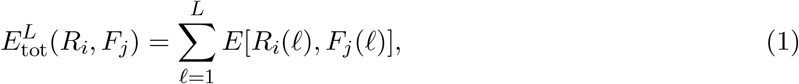

where *R*_*i*_(*ℓ*), *F*_*j*_(*ℓ*) ∈ {H, P, +, −}. If 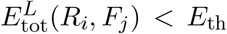, these proteins are considered matching, namely fertilization of a maternal S-haplotype carrying the RNase *R*_*i*_ by a paternal S-haplotype carrying the SLF *F*_*j*_ is enabled (Fig. 1e). We have also generalized our model to account for a soft energy threshold (see Methods and Fig. S15) and found that it retains our main results.

An S-haplotype is termed ‘self-incompatible’ if its RNase is active and it does not contain any SLF that matches this RNase (Fig. 1b) and ‘self-compatible’ otherwise (Fig. 1c-d). A diploid individual is ‘fully self-compatible’ if both of its S-haplotypes are SC and match each other, and ‘half self-compatible’ if exactly one of its S-haplotypes is SC and matches the other. Self-compatible S-haplotypes can be fertilized by either self or cross-pollen, whereas self-incompatible ones can only be fertilized by cross-pollen.

The population life-cycle is illustrated in Fig. 2. We initialize a population of *N* diploid individ-uals, constructed as shown in Fig. 1a-b. We assume non-overlapping generations. Every generation, each of the amino acids in each of the proteins can be mutated with probability *µ*. The new values of amino acids chosen to be mutated are randomly drawn from the prior amino acid distribution. Additionally, the RNase could alternate between active and inactive states with probability *µ*_R_. In each generation, the offspring population is produced one by one from the parental population by repeating the following mating procedure. A diploid individual is selected uniformly at random from the parental population to serve as a maternal parent. If the maternal S-haplotype is fully self-compatible, then with probability *α* it is self-fertilized, and with probability 1 − *α* it is outcrossed. If the maternal plant is half-self-compatible, self-fertilization and outcrossing occur with probabilities *α*′ and 1 − *α*′, respectively, where *α* and *α*′ correspond to the proportions of compatible self-pollen out of the total compatible pollen received. This model is known in the literature as ‘competitive selfing’ [13].

**Figure 2.**
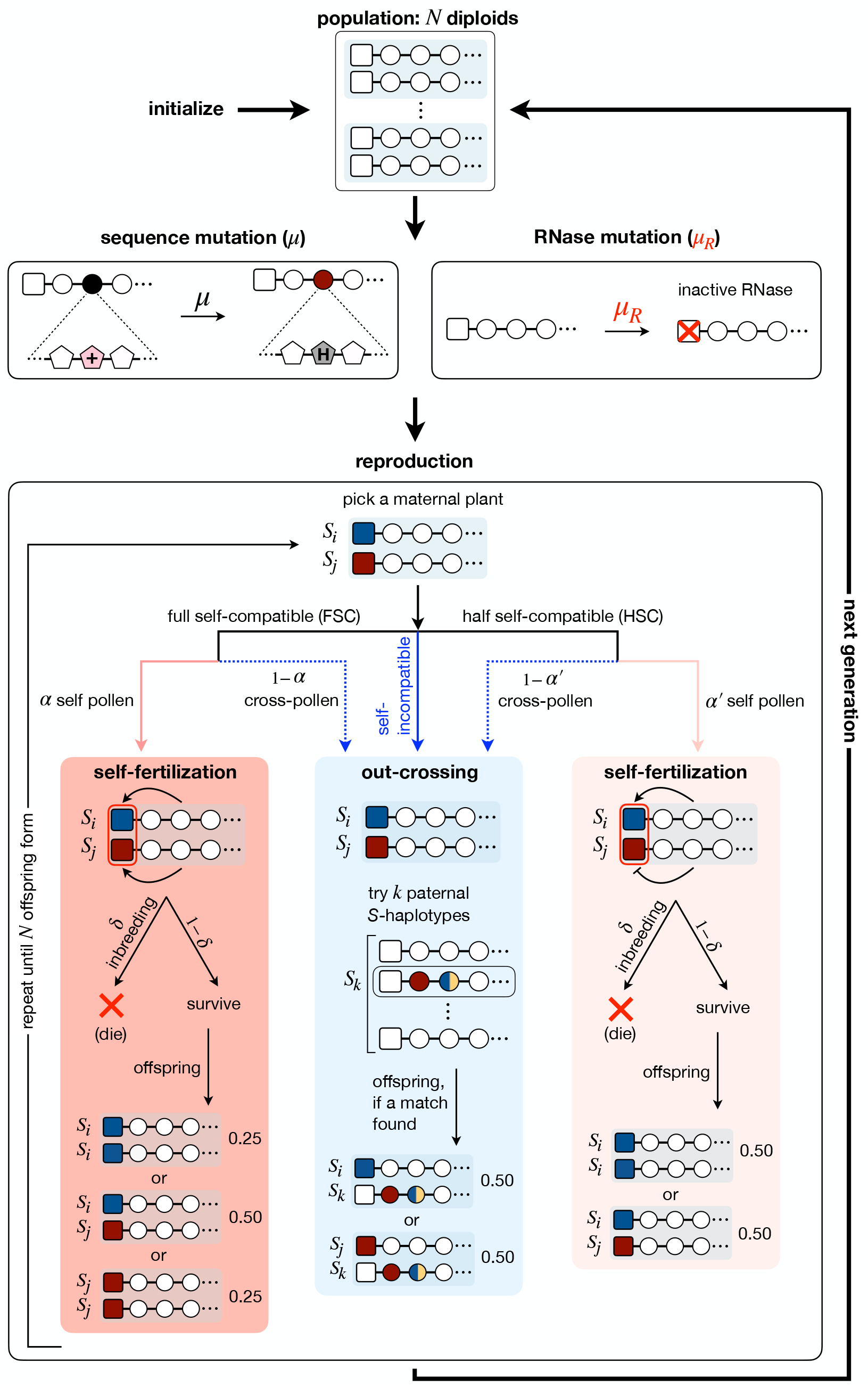
The population life cycle implemented in a stochastic simulation. The population consists of a fixed number *N* of diploid individuals, each composed of two S-haplotypes, as shown in Fig. 1. Every generation, each of the amino acids in each of the proteins can mutate with probability *µ* and each RNase could alternate between active and inactive states with probability *µ*_R_. We assume non-overlapping generations. Every generation, maternal plants are picked at random for reproduction. If the chosen female is full or half self-compatible, it can be fertilized by either cross-pollen or by self-pollen where self-fertilization offspring survive with probability 1 − *δ* relative to offspring formed by cross-fertilization. Alternatively, if the maternal S-haplotype is self-incompatible, only cross-pollination is possible. An outcrossed maternal plant is given up to *k* attempts to find a compatible paternal S-haplotype, by randomly picking S-haplotypes from the entire population and assessing their compatibility. If the maternal S-haplotype is self-compatible due to an inactive RNase, any paternal S-haplotype could fertilize. Following a successful fertilization, an offspring is produced. The offspring genotype obeys Mendelian rules. Maternal S-haplotype picking and offspring formation are repeated until *N* offspring are formed. The offspring population then replaces the parental population, completing one generation.

If the maternal plant (whether self-compatible or self-incompatible) is outcrossed, it receives *k* opportunities for cross-fertilization. In each attempt, an S-haplotype is randomly chosen from the parental population, and its compatibility as a male with the maternal parent is assessed. If it is found compatible, it serves as a paternal parent, and a diploid offspring is formed in accordance with Mendelian rules. To represent the waste of ovules by inbreeding depression, an offspring produced via self-fertilization is eliminated with probability *δ*, a parameter representing the inbreeding depression intensity. Hence, every such round can either result in a single offspring or none (due to failure of all cross-fertilization attempts or elimination of an offspring produced via self-fertilization). This procedure is repeated until a population of *N* offspring is formed. The maternal plant picking was implemented in the simulation using a computationally-efficient scheme to avoid unnecessary picking that results in inviable offspring (see Methods for details). The offspring population then replaces the parental one, completing one generation. We also tested a haploid version of this model, as in [61], providing qualitatively similar results (Supplementary text).

### The model exhibits three mating modes: fully self-compatible, fully self-incompatible, or mixed, depending on the inbreeding depression and promiscuity parameters

Self-incompatibility can occasionally break down via different mutations, forming self-compatible S-haplotypes that seemingly enjoy a twofold transmission advantage and a larger pool of potential mating partners. What are the sufficient conditions for the dynamically stable maintenance of complete self-incompatibility? Which parameters govern the choice of mating mode? The inbreeding depression intensity is considered a key driver of self-incompatibility. It was shown theoretically for simpler models of SI that a minimal value of inbreeding depression is a necessary condition for the maintenance of self-incompatibility [46, 16]. An additional parameter that plays a central role in our model is the promiscuity of molecular recognition between proteins, which governs the compatibility phenotype and the number of compatibility classes [58, 61]. Using stochastic simulations of our model, we tested the population mating mode depending on these two parameters: *δ* – the proportion of offspring produced via self-fertilization that are inviable, and *E*_th_– which sets the promiscuity of molecular recognition, to explore the parameter range for which self-incompatibility is sustainable in our model. We used a fixed population size of *N* = 500 diploid plants (1000 S-haplotypes). We assumed *n* = 9 SLF paralogs per S-haplotype, and each protein was represented by a sequence of *L* = 18 amino acids. The amino acid mutation rate was *µ* = 10^−4^ per AA per generation. Unless stated otherwise, we assumed pollen abundance, namely, a large number of cross-fertilization attempts per maternal plant chosen for reproduction. The simulations were typically run for 10^5^ generations (unless stated otherwise). To avoid transient effects, we discarded the initial *>* 50, 000 generations of our simulations and analyzed only results from that time point onward.

We found that the population attained either of three mating modes (Fig. 3a-b). Under high inbreeding depression and high promiscuity, the population was almost fully self-incompatible and spontaneously organized into mutually exclusive compatibility classes [58], such that each S-haplotype affiliated with a class is incompatible both as male and as female with all other members of its own class and conversely compatible both as male and as female with all members of all other classes. Note that the classification refers to S-haplotypes and not to diploid individuals (that are essentially heterozygous under self-incompatibility, and hence cannot be affiliated with a single class). While not all S-haplotypes were necessarily affiliated with any of the classes, typically only a small fraction remained unclassified (Fig. S11 for the diploid model and Fig. S27 for the haploid model). In contrast, under low inbreeding depression, essentially almost all S-haplotypes were self-compatible, and no class structure emerged. These two mating modes conform with previous analyses [46]. Yet, between these two regimes, we discovered an additional mixed population state, in which a temporally variable proportion of the population is self-incompatible, and the remaining part is self-compatible. The mixed state is characterized by large and non-decaying fluctuations in the population proportions of self-compatible and self-incompatible S-haplotypes. Previous works assumed that the mating mode is at steady state and obtained fixed SI-SC proportions [46, 47, 48, 49]. Such temporally non-decaying variation in the SI-SC proportions was not reported before. This volatile mixed regime stands in contrast to the other two mating regimes in which these proportions are essentially either one or zero, respectively, with only minor fluctuations (Fig. 3c). The mixed state is long-lasting and even after a very long time does not decay to any of the pure mating modes (Figs. S12-S13 for the diploid model, and Figs. S31-S32 for the haploid model).

**Figure 3.**
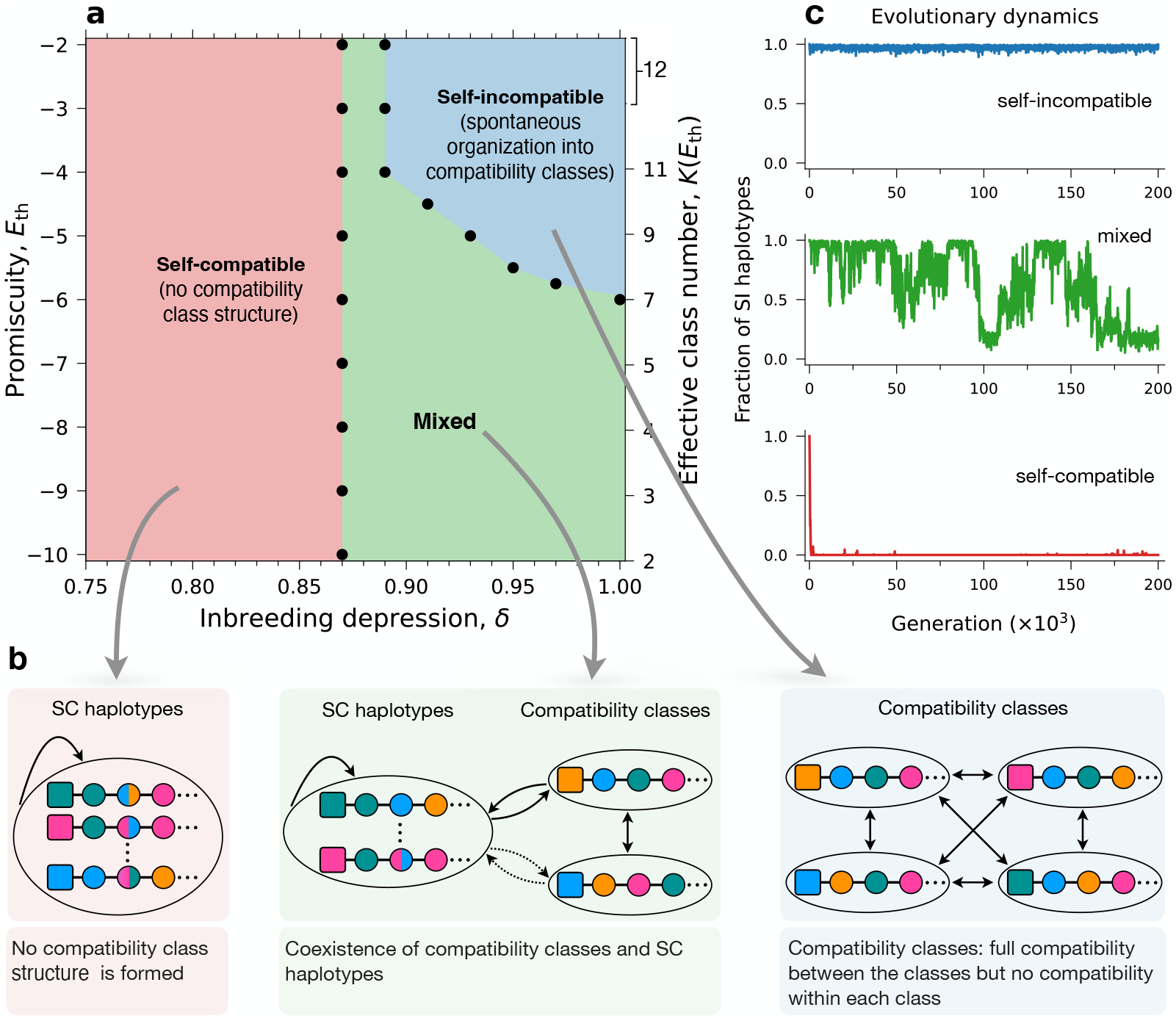
The model exhibits three different phases: fully self-compatible population, fully self-incompatible population partitioned into classes, or the dynamically unstable mixture of these two mating modes. **(a)** The phase diagram resulting from this model as a function of the inbreeding depression *δ* and the energy threshold *E*_th_ (promiscuity) parameters as obtained in simulations. The black dots mark the boundaries between the phases (see Methods). The secondary y-axis on the right shows the mean effective number of classes for each *E*_th_, and fixed *δ* = 1.0 as obtained in our simulations. **(b)** Schematic illustration of the population structure in the three phases. Under high inbreeding depression and intermediate-to-high promiscuity, the population clusters into distinct compatibility classes, each of which is fully compatible with all the others. At the other extreme of low inbreeding depression, the population is fully self-compatible, namely, each individual can self-fertilize, and no class structure forms. In between, the population exhibits a mixture of self-incompatible sub-population partitioned into compatibility classes, and a self-compatible sub-population that could potentially be compatible with the classes, too (dashed arrows). **(c)** Examples of the self-incompatible population fraction over time in the three phases. The self-compatible (top, blue) and self-incompatible (bottom, red) are dynamically stable, whereas the mixed phase (middle, green) exhibits vigorous fluctuations.

To characterize the time-scale of fluctuations in the self-incompatible population proportion, we calculated its temporal autocorrelation in the three different phases (Fig. 4f). We observe a decaying correlation in the population proportion of self-incompatible individuals in the mixed phase, in marked contrast to the more stable proportions in both the SI and SC phases, again marking it as a separate phase with unique temporal behavior.

**Figure 4.**
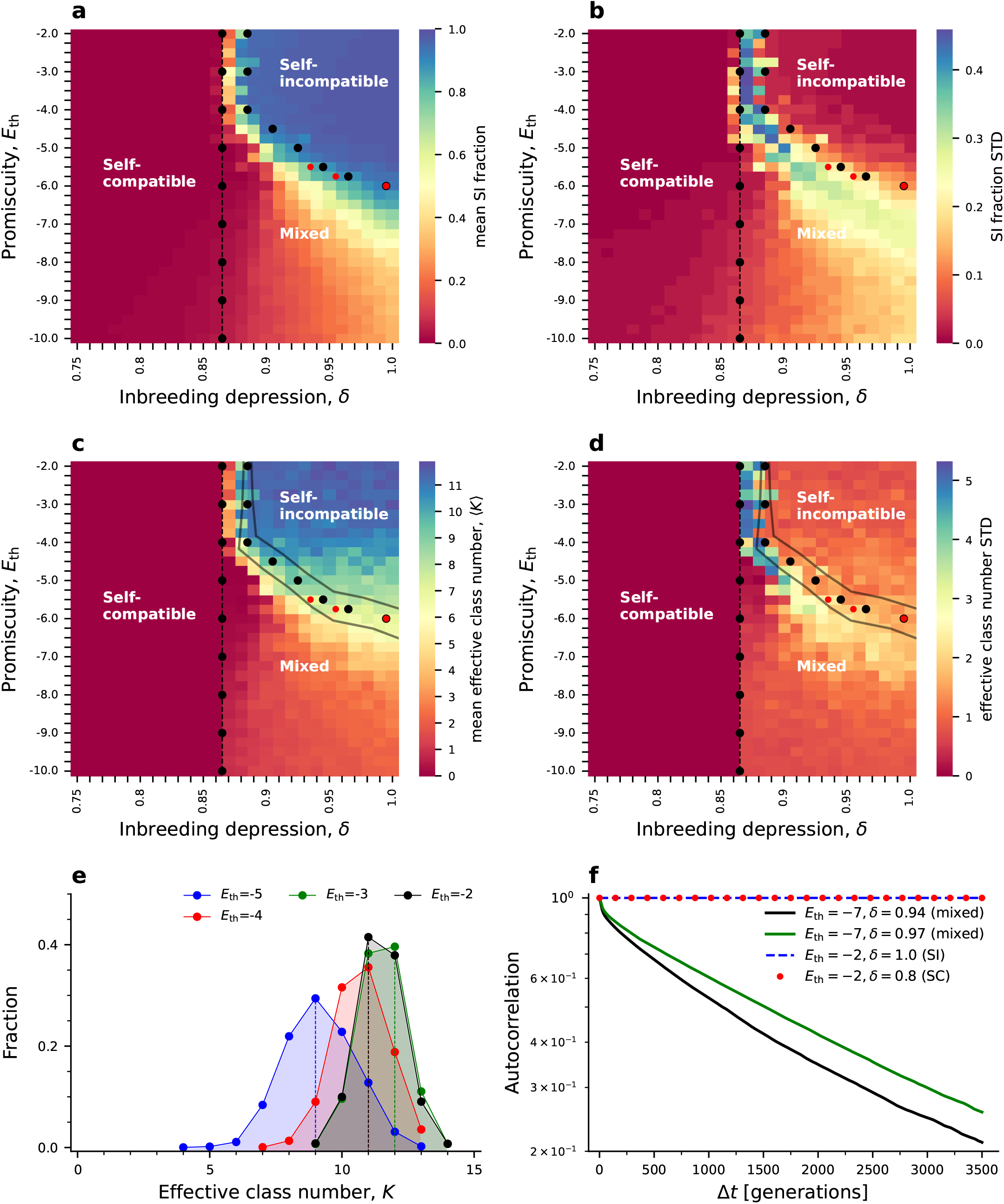
The number of classes exhibits two phase transitions in the *δ*-*E*_th_ plane. Maps of the mean **(a)** and standard deviation **(b)** of the SI population proportion, and mean and standard deviation **(d)** of the effective class number obtained in simulations for different combinations of the inbreeding depression *δ* and promiscuity *E*_th_. In the left part of the plane *δ < δ*^∗^, we find nearly zero SI proportion and zero classes, in accord with the full self-compatibility regime, as shown in Fig. 3. The right part *δ > δ*^∗^ occupies the mixed and self-incompatible phases observed in Fig. 3 and indeed shows either partial or full SI population proportion and a non-zero number of classes. The black dots in (a-d) mark the boundaries between the different phases and were calculated using the genotype proportions in the simulation data. The red dots show the analytically calculated *K* (Eq. (5)) mapped to *E*_th_, based on the simulation results. As a fixed *E*_th_ value produces a distribution of *K* values, we also illustrate ±*σ*_K_*/*⟨*K*(*E*_th_)⟨, where *σ*_*K*_ is the standard deviation, for the different *E*_th_ values (black solid lines). The boundary between the self-compatible and mixed phases is where the proportion of half SC genotypes is *x* = 0.1 (proportion of full SC genotypes is 0.90, and SI genotypes is ≈ 0), and the boundary between the mixed and self-incompatible phases shows the parameter combinations in which the proportion of full SC genotypes is 0.01. **(e)** Effective class number distributions for different *E*_th_ values at *δ* = 1.0. The dashed vertical lines mark the most probable class number for each *E*_th_. **(f)** The unbiased temporal correlation of the SI population proportion *r*(*t*), for two parameter combinations within the mixed phase (see Methods). For reference, we also show the autocorrelation in the SI (dashed blue) and SC (red dots) phases.

We analytically interpret the boundaries between these three regimes, similarly to a previous analysis by Charlesworth and Charlesworth [46]. To calculate the boundary between the full SC and mixed phase, assume that most S-haplotypes are self-compatible, denoted by *S*_C_, and only a small fraction of the population is SI. Assume that all SI S-haplotypes are of a single type *S*_1_. Thus, only two genotypes are possible: a full SC *S*_C_*S*_C_ genotype and a half-SC *S*_C_*S*_1_. *S*_C_ pollen is compatible with both *S*_C_*S*_C_ and *S*_C_*S*_1_ females. *S*_1_ pollen in contrast is only compatible with *S*_C_*S*_C_ females, but not with *S*_*C*_*S*_1_ because it is self-incompatible. Consequently, *S*_1_*S*_1_ offspring cannot be produced. As long as no additional SI types compatible with *S*_1_ emerge, no additional genotypes could form. To determine the inbreeding depression value *δ*^∗^ for which the proportion *x* of the half-self-compatible genotype is non-decreasing, we account for the proportion *x*′ of these genotypes in the next generation:

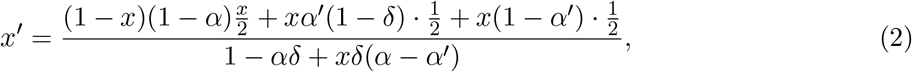

where *α* and *α*′ are the self-pollen proportions received by *S*_*C*_*S*_*C*_ and *S*_*C*_*S*_1_, respectively. In Eq. (2) we account for the formation of *S*_C_*S*_1_ offspring by *S*_C_*S*_C_ females (in proportion 1 − *x*) fertilized by non-self *S*_1_ pollen (with probability 1 − *α*), and *S*_C_*S*_1_ females (in proportion *x*) receiving either self or cross-pollen of type *S*_C_ (with probabilities *α*′ and 1 − *α*′, respectively), and normalize by the total offspring formed by all possible pollinations.

Assuming that in a half-self-compatible genotype both pollen types are produced in equal amounts, 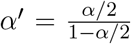. To obtain the minimal inbreeding depression value *δ*^∗^ for which the half-SC genotype proportion is non-decreasing we substitute *x* = *x*′ in Eq. (2), yielding:

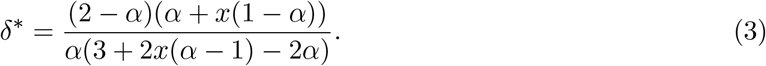

If we substitute *x* = 0, Eq. (3) will coincide with the boundary obtained analytically in [48] for the full SC region (denoted there as L). To obtain the second boundary, between the mixed and the full SI phases, assume that full SC genotypes are negligible and account only for full SI and half-SC genotypes. Assume the self-incompatible sub-population is composed of *K* equally-sized distinct compatibility classes, such that each is compatible with the remaining *K* − 1. We also assume that all SC S-haplotypes are compatible, both as males and as females, with each other and with all the SI S-haplotypes. Under these assumptions, we solve for the marginal inbreeding depression at the boundary as a function of *K*:

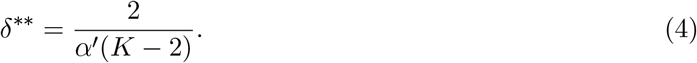

Substituting 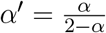 as before, yields

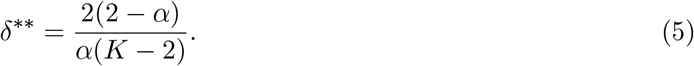

This equation is the special case of Eq. 4b in [46] under the assumption that full SC individuals are negligible. We conclude that the transition between the full SC and mixed phases occurs at a fixed inbreeding depression value *δ*^∗^. In contrast, the transition between the mixed and full SI phases occurs at an inbreeding depression value *δ*^∗∗^(*K*) that inversely depends on the number of classes, *K*. Thus, for a larger number of classes, a lower inbreeding depression is sufficient to stably maintain full SI. The intuitive explanation is that the disadvantage of SI is its inability to fertilize as male females of its own class. The larger the number of classes, the smaller the population proportion of each class, thereby diminishing this penalty.

Substituting *α* = 0.6 as used in our simulations, and *x* = 0.1, we obtain *δ*^∗^ = 0.87. In Fig. 4a-b, we present the self-incompatible population proportion obtained in our simulations and plot the calculated boundaries (black dots) on top. The boundary between the mixed and full SI regimes was calculated for a 1% population proportion of full SC genotypes. We find excellent agreement between the analytically calculated boundaries (red dots, lines) and the simulation results (black dots).

We note that the existence of three mating regimes is relatively insensitive to the exact parameter values chosen (but see more detailed discussion of special cases below). The choice of *x* = 0.1 is arbitrary, and the boundary shows little sensitivity to the exact value chosen (see boundaries obtained for different *x* values in Fig. S25 for the haploid model).

### The population self-incompatibility pattern exhibits two phase transitions

Our analysis so far (Eq. (3), Eq. (5) and Fig. 3) demonstrated three mating modes, or ‘phases’, where the boundaries separating them (‘phase transitions’) were determined by two fundamental parameters of the system: *δ* and *E*_th_.

A crucial property of the self-incompatible phase in our model is the spontaneous self-organization of almost all S-haplotypes into compatibility classes, such that all individuals in the same class are bidirectionally incompatible with each other and bidirectionally compatible with all members of all other classes [58]. Having found that the unstable mixed phase exhibits a temporally fluctuating proportion of self-incompatible individuals, it is of interest to determine whether the classification property holds in this phase as well, and, if so, whether it is affected by the volatility of the potential population that could be classified. Since the classes are not necessarily of equal size, we refer below to the ‘effective class number’ defined as 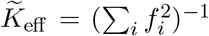, where *f*_*i*_ is the population proportion of the *i*-th class (out of the classified SI sub-population) [64]. The effective class number can also be directly calculated without first classifying the population (Methods). In Fig. 4c-d, we plot the mean and STD, respectively, of the effective class number over many simulation time points, for the entire (*δ, E*_th_) phase space. The boundaries obtained in simulation are shown as black dots, and analytically calculated boundaries are shown as dashed lines (SC-mixed) and as red dots (mixed-SI). In the volatile mixed phase, too, we observe the formation of classes that include almost the entire SI sub-population. The class number, however, is, on average, lower than in the self-incompatible phase under the same *E*_th_ value. This is partially explained by the temporal changes in the SI population proportion, as a larger SI population affords the existence of a larger number of classes and vice versa [58]. Remarkably, the volatile mixed phase stands out as having much higher temporal variation in the class number compared to the self-incompatible phase (Fig. 4d, Fig. S14). How does the class number depend on the model parameters *δ* and *E*_th_? We find that within the SI phase the class number increases with *E*_th_, but is insensitive to *δ*. In Fig. 4e we show distributions of the effective number of classes obtained in the self-incompatible regime (high inbreeding depression) for different values of *E*_th_, showing this effect.

We observe two phase transitions in this model, each with distinct characteristics. The transition from the self-incompatible to the volatile mixed occurs in a relatively narrow parameter range, and is marked not only by the transition between stable to unstable SI population proportion (Fig. 4b), but also by a decline in the class number average, and a sharp increase in its variance. In contrast, the phase transition between the SC (zero classes) and mixed (non-zero classes) occurs over a broad parameter range and is smooth.

How do the different phases depend on the promiscuity parameter? For low *E*_th_, the mixed phase is broad, and there is no full SI phase, which only appears at intermediate *E*_th_. The range of *E*_th_ values affording full SI broadens under larger population sizes (Fig. S5) and higher mutation rates (Fig. S7). The mixed phase narrows as *E*_th_ increases, and the transition to the SI phase becomes sharper. The varying width of the mixed phase is explained by the increase of the class number with *E*_th_, rendering self-incompatibility favorable under lower inbreeding depression.

### The volatile mixed phase robustly exists under different model parameters, but shrinks if cross-pollen is limited

So far, we have considered self-compatibility caused solely by mutations at the SLF-RNase interaction interfaces, which could make an SLF compatible with the same-haplotype RNase. Empirical results have shown an additional route to self-compatibility via various loss-of-function mutations in the RNase (e.g., stop-codon or inactivation) [37]. This alternative mechanism renders the S-haplotype carrying the mutated RNase not only self-compatible but also compatible with any pollen regardless of its SLF sequence. In additional simulations, we tested both mutation types together. We mimicked the RNase mutations by letting RNases alternate between active and inactive states at a rate *µ*_R_ (Fig. 1d).

We found that when both mutation types (sequence and RNase inactivation) were allowed to cause self-compatibility, the volatile mixed phase was obtained too, and the phase diagram was very similar to the one obtained under sequence mutations alone (Fig. S4). We also found that the phase diagram and, in particular, the existence of the mixed phase were insensitive to simulation parameters such as the population size (Fig. S5) and the mutation rate (Figs. S6-S7), though its boundaries shifted.

Two model parameters we found to affect the phase diagram were *α*, the proportion of self-pollen received (Fig. S16), as it affects the impact of inbreeding depression, and *k*, the number of fertilization attempts per maternal plant. The results shown in Fig. 4 were all obtained under *α* = 0.6 and a large *k* representing pollen abundance. In Fig. 5 we show the case of pollen limitation, when each female is provided only a single fertilization attempt, namely *k* = 1. We have previously shown that the self-incompatibility proteins evolve under a combination of selection pressures to simultaneously avoid self-compatibility and enhance cross-compatibility, shaping the amino acid content of their interaction domains [61]. Under pollen abundance, females are essentially guaranteed to be fertilized. Consequently, the pressure to match non-self is almost exclusively borne by males, while females are mostly subject to the pressure to avoid self-compatibility. Pollen limitation shifts the balance between these selection pressures, placing females, too, under pressure for cross-compatibility. This in turn causes an increase in the number of compatibility classes, to reduce the pollen proportion a female rejects (compare Fig. 5c to Fig. 4c). Consequently, the advantage of the full SI phase over the SC and mixed phases extends over a larger range of parameter values. Indeed, we observe that under pollen limitation the full SI phase expands to the entire *E*_th_ range (whereas under pollen abundance it was only obtained under intermediate to high *E*_th_), and the mixed phase shrinks to only a narrow stretch of *δ* values (compare Fig. 5a-b to Fig. 4a-b). Pollen limitation is not captured by our equations, which assume that female fertilization is guaranteed. Similarly, not limiting the number of fertilization attempts per female, *k* = 1000, but increasing the proportion of self-pollen to *α* = 0.9, yielded a similar phase diagram with a narrow mixed phase (Fig. S8).

**Figure 5.**
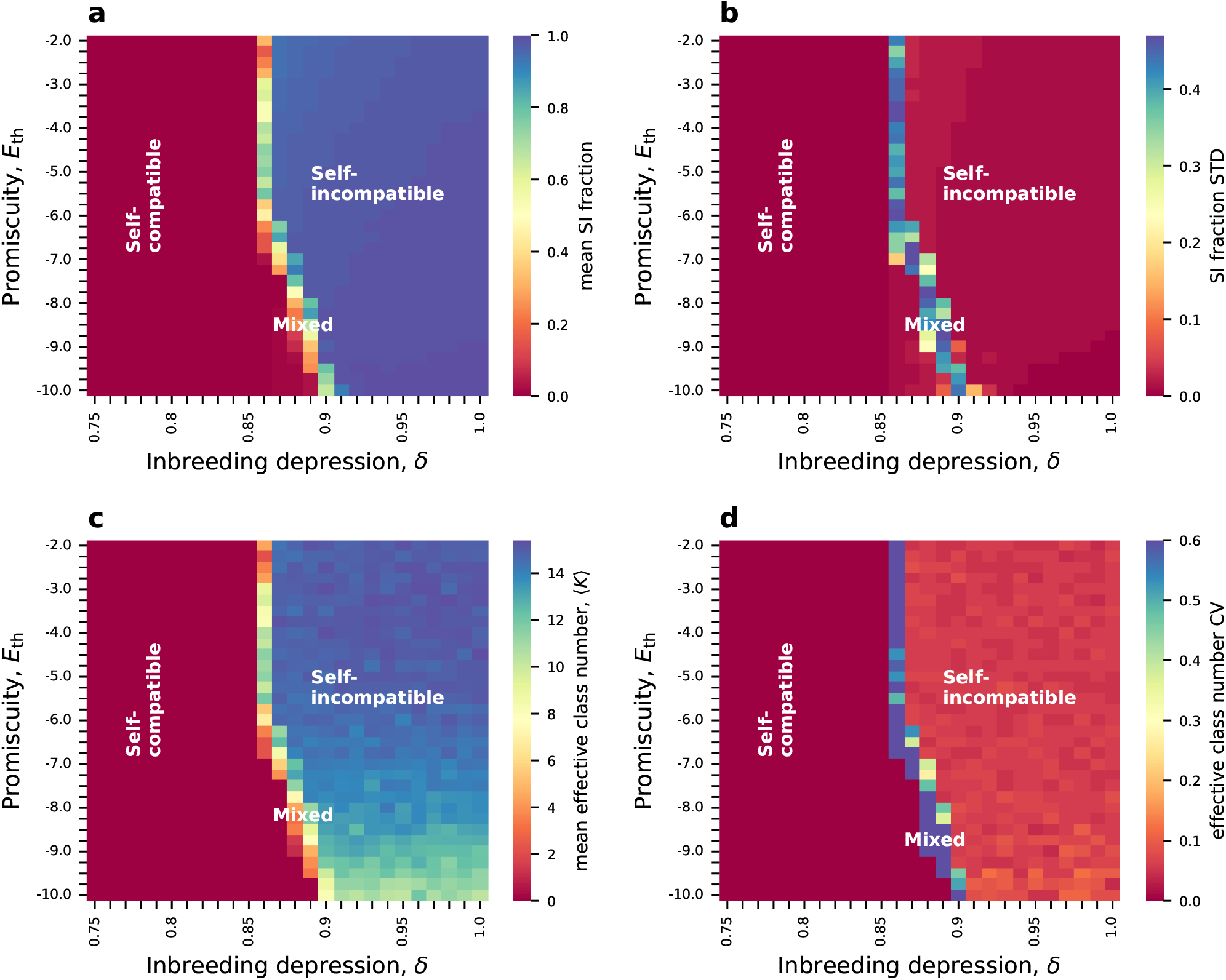
Under pollen limitation, the parameter region in which full self-incompatibility is favored significantly expands, and the region exhibiting the mixed phase shrinks. **(a-b)** Maps of the mean and standard deviation of the self-incompatible population fraction. **(c-d)** The mean and the CV (std/mean) of the effective class number with only a single cross-fertilization attempt per female, *k* = 1, mimicking pollen limitation. We observe that the mixed unstable phase still exists, but for a very narrow range of *δ* values. Another difference is that the full SI phase exists for the entire range of promiscuity values, unlike the pollen abundance case, in which it only exists for intermediate to high promiscuity. The self-pollination rate is *α* = 0.60. The figures are based on 4 independent runs for each point in *δ*-*E*_th_ plane ∈ [0.75,0.84], and [0.91,1.0], and 32 runs for *δ* ∈ [0.86, 0.91]. From each run, we extracted 2000 (a-b) and 600 (c-d) data points at 25-generation intervals, respectively. The minimal class size is 10 S-haplotypes.

### Transitions between self-incompatibility and self-compatibility are reversible in part of the parameter range

While Fig. 3 shows that a population can be in either the full self-compatible, full self-incompatible, or the mixed state depending on the inbreeding depression, the dynamical stability of these states to ongoing mutations is of great interest. The simulations of Fig. 3 were all initialized with a fully self-incompatible population. To test the dynamical stability of the population mating mode, we repeated the simulations, this time initializing them with a fully self-compatible population. The complete phase diagram obtained is shown in Fig. 6. We find that under a great part of the *E*_th_ range, the population’s final state is independent of its initial state, and the phase diagram is similar to that of Figs. 3-4. The case for high promiscuity (*E*_th_ ≥ −4.5) is markedly different, though. There, a population starting as self-compatible under inbreeding depression intensity high enough to support full self-incompatibility, as shown in Figs. 3-4, remains self-compatible even after a large number of generations. The explanation for this distinct behavior is that, under high promiscuity, the probability of a match between proteins is so high that mutations producing self-incompatible S-haplotypes become extremely rare, leading to infeasibly long waiting times for such mutations. As a back-of-the-envelope estimate, under *E*_th_ = −2 the probability that two random proteins do not match is 0.03. With 9 SLFs per S-haplotype, as in our simulations, the probability that none of these SLFs matches the same-haplotype RNase is as low as 10^−14^! To validate this explanation, we repeated the simulations, modifying them only by seeding a few self-incompatible mutants within the otherwise self-compatible population. This test indeed resulted in a transition to self-incompatibility, verifying our hypothesis (Fig. S10 for the diploid model, Fig. S28 for the haploid model). We conclude that, under high promiscuity, self-compatibility is effectively an absorbing state from which the transition back is infeasible once the entire population becomes SC, regardless of the level of inbreeding depression. However, for intermediate-to-low promiscuity, transitions between mating modes are reversible, and the population’s mating mode at equilibrium depends on the inbreeding depression and promiscuity parameters, but not on the population’s initial state.

**Figure 6.**
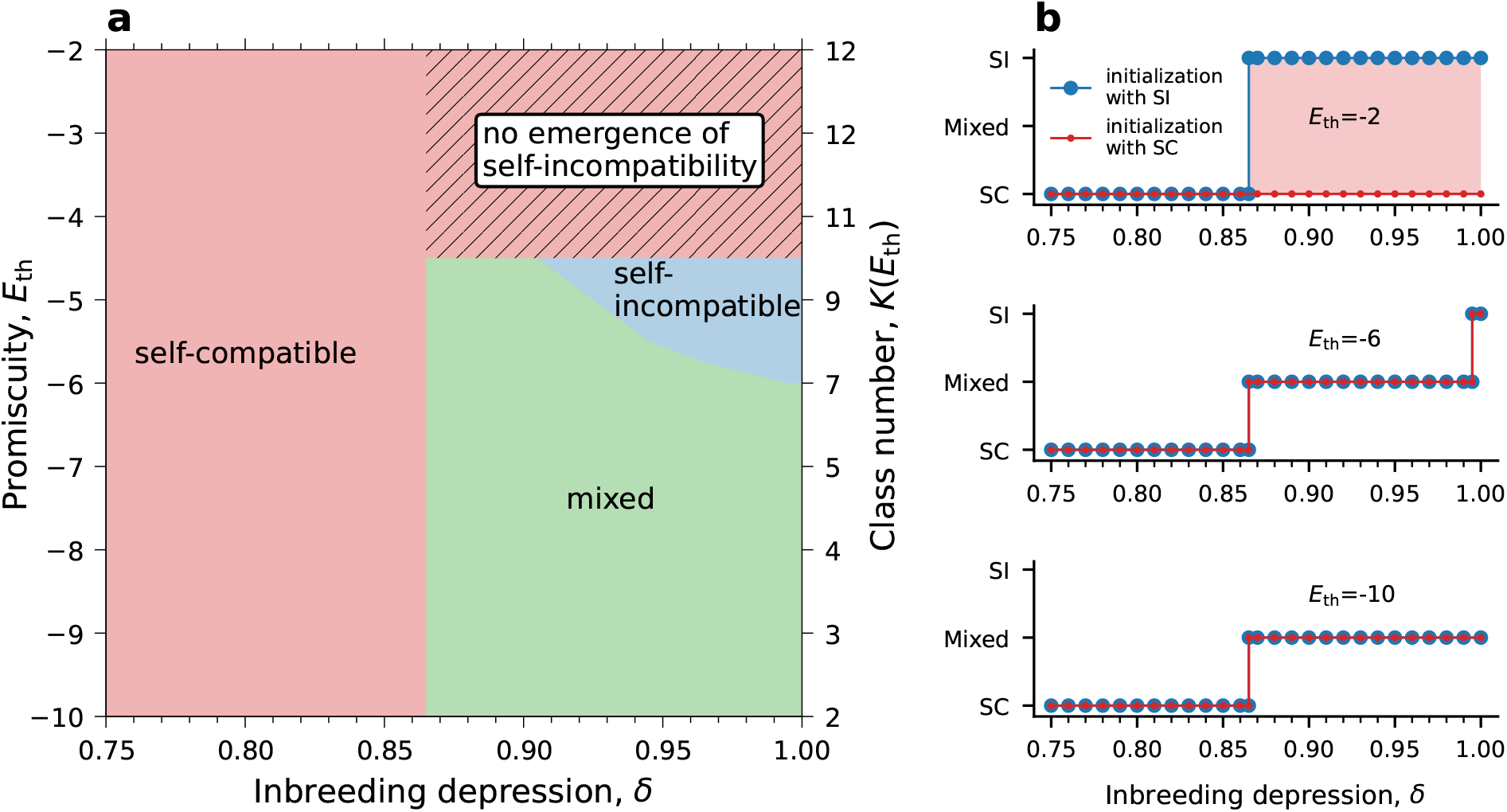
Under intermediate-to-low *E*_th_, the phase diagram is independent of the population’s initial condition: whether it starts as fully self-compatible or fully self-incompatible. **(a)** Phase diagram obtained when the population was initialized as fully self-compatible. Over most of the parameter range, we observe the same three phases with boundaries similar to those in Fig. 3, where the population was initiated as fully self-incompatible. Only for high promiscuity *E*_th_ ≥ −4.5, self-incompatibility does not emerge in realistic time (region marked by diagonal stripes). We mapped *E*_th_ into the mean effective number of classes, calculated at *δ* = 1.0 (right-side y-axis). To obtain this diagram, we ran simulations for different combinations of the parameters *E*_th_ (with a step size of 0.25) and *δ* (with a step size of 0.03) – 4 independent runs for each combination. Each simulation was run for 1 × 10^5^ generations, but we discarded the initial 0.5 × 10^5^ and analyzed only the remaining 50,000 generations. We used 2000 data points with 25-generation intervals between consecutive time points from each run. The boundaries between the phases were set as the parameter combination for which, on average, over these multiple data points, the population SI proportions equal the threshold values. **(b)** The population mating mode at steady state (SC, SI or mixed) as a function of the inbreeding depression *δ* for three different *E*_th_ values, starting from either a fully self-incompatible (red) or a fully self-compatible (blue) population. This graph is a cross-section of the phase diagrams of Fig. 3a starting from full SI, and the analogous one in (a) starting from full SC. Parameter values: *α* = 0.60.

## Discussion

Self-incompatibility is a widespread genetic mechanism that prevents self-fertilization and is found in ~40% of plant families. Because self-incompatibility relies on the coordinated action of multiple components, it is susceptible to mutations that can break it down, creating self-compatible plants capable of self-fertilization. Indeed, natural population surveys revealed many instances of self-compatibility in otherwise self-incompatible species, where some populations show a mixture of self-compatible and self-incompatible plants in the same population, and a few populations are fully self-compatible [27, 56, 55, 57]. Despite the prevalence of both SI and SC, the dynamics and reversibility of the transitions between these mating modes remain theoretically murky. Here, we focus on these questions in the context of the collaborative non-self-recognition self-incompatibility mechanism, found in the Solanaceae and Rosaceae families. We build on a previously proposed model that specifically accounts for the biophysical basis of self-incompatibility – the molecular recognition between the male and female self-incompatibility proteins. In particular, this model explicitly describes the mutations that could cause transitions between mating modes [58, 61]. Most importantly, the number of mating specificities, a crucial determinant of the balance between self-compatibility and self-incompatibility, is an emergent property of the model rather than a fixed parameter.

Our main finding is that the population pursues either of three mating modes: two pure ones - full self-compatibility (SC), and full self-incompatibility (SI) - and a volatile mixture of both (Figs. 3-4). Under full self-compatibility, every S-haplotype is compatible with almost every member of the population, including itself. Under full self-incompatibility, the population spontaneously self-organizes into mutually exclusive classes, such that every S-haplotype is compatible with all members of all other classes except its own. In the mixed state, a self-compatible sub-population co-exists with a certain number of compatibility classes, exhibiting high but incomplete compatibility between SI and SC sub-populations (Fig. S1). We find that both the self-compatible and self-incompatible phases are dynamically stable, but the mixed phase exhibits vigorous perpetual fluctuations in the SI and SC population proportions and in the number of SI classes. In particular, it does not decay to either of the pure long-lived mating modes, nor to their stable co-existence at fixed proportions, even after a very long time (Figs. S12-S13 for the diploid model, and Figs. S31-S32 for the haploid model). Around the boundaries between the different mating modes, we find that the population could alternate between mixed (both SI and SC) and pure compositions, yet unlike the pure phases, the residence time at each composition is short, and the population never settles at one of them (Fig. S18). The mixed phase is consistently obtained under different population sizes (Fig. S5), different mutation rates (Figs. S6-S7), and different types of mutations causing self-compatibility (Fig. S4), but shrinks if the supply of cross-pollen is limited (Fig. 5, Fig. S8).

The existence of multiple mating modes represents reproductive trade-offs: self-compatible females incur offspring loss due to inbreeding depression, while self-incompatible males suffer from reduced mating opportunities because they cannot fertilize females of their own class. The magnitudes of both penalties depend on the model parameters: the proportion of self-pollen *α* and the inbreeding depression value *δ* determining the harmful effects of self-fertilization, and the promiscuity of molecular recognition captured by *E*_th_ (Figs. 3-4). *E*_th_ dictates the number of classes: When a new RNase variant establishes a new class, it might be incompatible with an existing class’s SLFs, leading to the extinction of that class. The likelihood of such incompatibility diminishes as the promiscuity of interactions, represented by *E*_th_, increases. The number and size of compatibility classes determine the fitness disadvantage males face due to incompatibility with respect to females of their own class. In particular, small classes enjoy a reproductive advantage over large classes because pollen from small classes is rejected by a smaller proportion of the population’s females than pollen from large classes. Based on these reproductive trade-offs, we compute the boundaries between the different mating modes depending on the model parameters *α, δ*, and the emergent class number *K*(*E*_th_). The boundary between the full SC and mixed regimes is defined as the point at which the population is mostly SC and only a single SI type, occupying a small, arbitrary proportion of the total population, exists. For a fixed *α*, this boundary is obtained for a fixed inbreeding depression value *δ* = *δ*^∗^, strictly larger than 1/2 and essentially independent of *E*_th_. The boundary between the mixed and fully self-incompatible regimes is defined as the parameter combination for which the SC population proportion does not exceed a small, arbitrary value. This boundary depends not only on the inbreeding depression, but also on the promiscuity via the number of classes *K*. We compute the boundaries *δ*^∗^ and *δ*^∗∗^(*K*(*E*_th_)) (Eqs. (3)-(4)), finding excellent agreement with our simulation results. While these equations predict a parameter region between the two boundaries where a mixture of SI and SC sub-populations exists, they fail to predict the volatility of this mixture, as observed in our simulations.

Our analysis recovers previous results regarding the necessary conditions for the sustainability of self-incompatibility. Specifically, our result showing that a minimal inbreeding depression *δ*^∗^(*α*) (boundary between full SC and mixed phases) is a necessary condition for self-incompatibility is in line with previous studies [16, 46, 48, 62]. Our result that *δ*^∗∗^ = *δ*^∗∗^(*K*) such that for a larger number of classes, a lower level of inbreeding depression suffices to ensure full self-incompatibility, are in line with [46, 16]. In particular, our equations coincide with the analysis of Charlesworth and Charlesworth [46], yet in their equations the number of classes was a free parameter, whereas in ours it emerges as a model outcome. Previous theoretical works [46, 48, 26, 47, 49] also predicted a parameter regime supporting an SI-SC co-existence and calculated their proportions under the assumption of steady state. Interestingly, in our model, this SI-SC co-existence regime reveals a highly fluctuating SI/SC ratio. To the best of our knowledge, such a volatile mixed regime was not predicted by previous models. Notably, some of these studies modeled a different SI mechanism based on self-recognition, for which class emergence and decay are driven by processes distinct from those in non-self recognition.

What are the forces triggering the non-decaying instability in the mixed phase? Why doesn’t the population relax to one of the two stable mating modes? We previously studied the dynamics of class emergence and decay for this mechanism in the full SI phase. There, we found that under the full SI regime, classes are usually born and die one at a time [58]. Yet, in the mixed phase, it is common for multiple classes to appear or disappear within a short time window, demonstrating avalanche-like dynamics. What exactly stimulates this dynamics remains obscure. We leave it for future work to thoroughly map these class mass extinction and emergence events and to study the sources of differences in dynamics relative to the SI phase.

The RNase-based SI mechanism relies on non-self recognition, such that each S-haplotype should match as a male essentially all non-self female types in the population [25]. Under pollen abundance, the establishment of a new female type matched by some pollen types, but not by others, could cause the extinction of the non-matching pollen types [65, 58]. This stands in contrast to self-recognition SI mechanisms, in which pollen should match only a single female type — the self one; hence, type extinctions are not driven by new mutations but likely occur only through drift. Thus, we propose that mass extinctions of mating specificities are unique to the non-self-recognition SI. Different plant families, such as Brassicaceae and Papaveraceae, have distinct SI mechanisms that rely on self-rather than non-self recognition. To support our prediction, it remains to verify, both theoretically and empirically, whether the volatile mixed phase does not exist under self-recognition SI mechanisms.

Extreme pollen limitation, as may occur during the colonization of new territories or when pollinators are scarce, is often considered a trigger for the transition to self-compatibility as a means of reproductive assurance. In such cases, while cross-pollen is rare, self-compatibility could be favorable even under higher values of inbreeding depression [47]. Intriguingly, we find that coupling the possibility for SC invasion with SI diversification, as in our model, not only does pollen limitation not trigger the transition to self-compatibility, but conversely, self-incompatibility extends to a broader range of promiscuity values that otherwise yielded mixed mating (Fig. 5). The intuitive explanation for this result is that the AA sequences of the male and female proteins evolve under a combination of two types of selection pressures: to avoid self-compatibility and to enhance cross-compatibility. Under pollen abundance, as assumed in most of our simulations (except Fig. 5), the females are essentially guaranteed to find a male partner and, within a good approximation, are subject only to the pressure to avoid self-compatibility. Pollen limitation shifts the balance between these selection pressures, such that females, too, are under pressure to match. This, in turn, makes females more matchable and, consequently, leads to a higher number of compatibility classes. This increases the full SI-mating parameter range at the expense of the mixed mating regime, as shown in Fig. 5. The balance between the selection pressures operating on these proteins was analyzed in our previous study [61].

There are conflicting opinions regarding the reversibility of transitions from SI to SC, where SC is often considered an evolutionary ‘dead-end’ [29, 12]. Here, we find that across a large part of the parameter range in our model, the population’s mating mode at steady state is independent of its initial mating mode – whether self-compatible or self-incompatible. Thus, we argue that transitions from self-incompatibility to self-compatibility could be reversible. The exception is the high promiscuity range (*E*_th_ ≥ −4.5), for which mutations transforming a self-compatible S-haplotype into a self-incompatible one are exceedingly rare, rendering the waiting time for the reverse transition infeasibly long (Fig. 6, Fig. S10). Related to that, the emergence of full self-incompatibility from a fully self-compatible population was shown to be possible by Sakai [62] in a model allowing for continuous rejection probabilities between S-haplotypes.

Inbreeding depression is the outcome of many recessive deleterious mutations [11], and hence, its magnitude can vary between individuals and evolve. In particular, inbreeding depression could be affected by the prevailing mating mode – prolonged selfing can purge deleterious mutations and decrease the inbreeding depression load [66]. Still, the purging efficacy depends on the fitness effect of these mutations [16, 26]. The time window during which the reverse transition from SC to SI is possible is deemed short because the decrease in inbreeding depression following prolonged selfing renders full SI less favorable and requires a larger number of mating specificities for its maintenance, while in parallel the number of mating specificities rapidly decreases following the SI system breakdown [42]. The possible loss of mating specificities is well captured by our model; however, we did not account for the evolution of inbreeding depression levels, assuming they were fixed and uniform across the population. Hence, our model only describes the short term when this assumption still holds. This simplifying assumption was also employed in most previous models [46, 62, 63, 65]. An exception is Lande and Schemske’s model [16], in which the inbreeding depression evolves in response to the mating mode, yielding only two long-lived mating modes: full selfing or full outcrossing. Mixed mating, as detected in natural populations, was considered by the authors to be transient. A possible extension of our model could incorporate an evolving level of inbreeding depression, with feedback from the mating mode. It would then be interesting to study whether, under this extended model, the transitions to self-compatibility remain reversible and whether the volatile mixed phase persists or diminishes. We hypothesize that the outcome should strongly depend on the model details: the fitness effects of deleterious mutations and how rapidly they are purged under selfing, if at all [16, 26, 11].

Mixed populations containing both SI and SC genotypes were detected in the wild [56, 57, 67, 37]. Still, because long-term studies tracking the proportions of SI and SC genotypes in these populations are lacking, we do not know whether they exhibit the theoretically predicted volatility of the mixed phase. If such mixed populations exhibit frequent class extinctions, as we propose, they are expected to decrease the effective population size and consequently reduce the population genetic diversity relative to purely outcrossing populations [68]. Repeated selfing is also expected to elevate genome-wide homozygosity. That could be detected in the genomes of plants in mixed populations, even if they are currently SI, suggesting frequent transitions between mating modes. We also predict that compatibility classes in mixed populations are fewer and shorter-lived compared to full SI populations. Long-term studies in populations exhibiting mixed mating and computational tests comparing the magnitudes of genetic diversity, homozygosity, and age of compatibility classes across populations employing different mating modes could validate our theoretical prediction.

Climatic changes, in particular global warming, are expected to affect plant mating modes in different ways. Plants that should adapt to varying conditions are expected to favor cross-fertilization over self-fertilization. Yet, potential depletion of pollinating insects and plant migration to new territories following climatic changes could render self-fertilization more advantageous [43, 44, 45]. A better understanding of the relationship between these factors and the plant mating mode should help minimize their harmful effects on plant populations and conserve natural habitats.

In summary, while simplified theoretical models were sufficient to predict the essential thresholds that sustain a self-incompatible population with a given number of classes, the study of a detailed biophysical compatibility model – where classes spontaneously emerge and decay – allowed us to pinpoint the role of molecular recognition promiscuity in controlling the number of compatibility classes. It provided a complete phase diagram for the different mating modes and, where applicable, the number of classes, given in terms of inbreeding depression and molecular promiscuity, and revealed a previously undiscovered mating mode. We established the reversibility of transition between mating modes across part of the parameter space. It remains to validate the existence of the volatile mixed mating mode through the collection and analysis of field data, and to further theoretically study the mechanism driving the rapid fluctuations in the population proportion of the self-compatible S-haplotypes in the mixed state, which pertains to the irregular emergence and decay of classes.

## Acknowledgements

We thank Nils Becker, Yonatan Friedman, and Avi Mayo for their comments on the manuscript and Oded Agam for useful discussions. This research was supported by the Israel Science Foundation, grant numbers 1889/19 and 2991/24 (T.F.). A.J. acknowledges funding by the Lady Davis Fellowship Trust for post-doctoral researchers. T.F. warmly thanks the Erwin Schrödinger Institute in the University of Vienna, for hospitality in June 2025 during and after the thematic program on Extreme Value Statistics in Biology, during which part of the work on this manuscript was carried out.

## Methods

### Classification of the amino acids into our biochemical categories

We have classified the 20 AAs into four biochemical categories: hydrophobic (H), neutrally polarized (P), positively charged (+), and negatively charged (–); see Table S1. The prior frequencies of these AA biochemical categories, from which a value is chosen when an amino acid is mutated, are given by *ν*_*i,i*∈{H,P,+,_−_}_= {0.5, 0.265, 0.113, 0.122}respectively [58]. The interaction energies used in our model between all possible pairs of amino acids are detailed in Table 1.

### Parameters and default values

The model parameters and their default values (unless specified otherwise) are summarized in Table The parameter *E*_th_ was varied in the range −10 to −2 with a step size of Δ*E*_th_ = 0.25. The parameter *δ* was varied in the range 0.75 to 1.0 with a step size of Δ*δ* = 0.01 in the diploid model, and from 0.0 to 1.0 with a step size of Δ*δ* = 0.03 in the haploid model.

### The stochastic simulation

#### Initialization of the S-haplotype population

We initialize the S-haplotype population with *n* distinct S-haplotypes, where each type is self-incompatible and bidirectionally compatible with the remaining *n* − 1 S-haplotypes. We employ the following three steps to draw these initial types:

1. **Draw RNases sequences**: First, generate *n* distinct RNases R = {R_1_ … R_*n*_}by randomly drawing sequences from the distributions of AA frequencies obtained in evolved RNases at steady state. Different distributions were used for different *E*_th_ values – see Table S2.
2. **Draw SLFs sequences**: Construct *n* pools of SLF sequences such that each pool P_*i*_ = {S_*m*_}, *i* ∈ (1 … *n*), contains *m* SLF sequences, that can all detoxify the *i*-th RNase, and potentially others as well. To construct the SLF pools, we generate SLFs, one at a time, check which RNase(s) they detoxify, and add them to the corresponding pool(s). Any SLF that does not detoxify any RNase, or that detoxifies all RNases, is discarded. Repeat this step until each pool has *m* SLF genes. The SLF sequences are drawn from the prior distribution *ν*.
3. **Form** 2*N* **self-incompatible and complete S-haplotypes**: For each RNase R_*i*_ ∈ R, choose in total *n* − 1 SLF genes, one from each pool P_*j*_, *j* ∈ (1 … *n*), *j* ≠ *i* and make sure none of these SLFs detoxifies R_*i*_. If it does, replace it with another SLF from the same pool until an SLF that does not detoxify R_*i*_ is found. After generating *n* distinct S-haplotypes, replicate each type 2*N/n* times, to obtain a total of 2*N* S-haplotypes, and shuffle their order.
4. **Form** *N* **self-incompatible diploid plants**: Produce *N* diploid individuals by randomly joining together pairs of distinct S-haplotypes. Index the S-haplotypes such that every two homologous S-haplotypes receive consecutive indices, e.g., H_1_H_2_, H_3_H_4_, … H_2*N*−1_H_2*N*_.

The above procedure produces SLFs that can potentially match multiple maternal plants.

### Main model life-cycle: mutation and reproduction

1. **Mutation**: Each position in each protein can mutate with probability *µ* per generation per AA. If a particular position is chosen to mutate, the amino acid currently existing there is replaced by a randomly chosen amino acid drawn from the prior distribution *ν*.
2. **Female plant picking**: Female plants chosen to reproduce can be either FSC, HSC (allowing both self and cross-fertilization), or SI (allowing only cross-fertilization). In the case of self-fertilization, only a proportion 1−*δ* of the self-offspring survives. We compute the frequencies of offspring produced by either type of female, where we distinguish between self (s) and non-self (ns) fertilization:

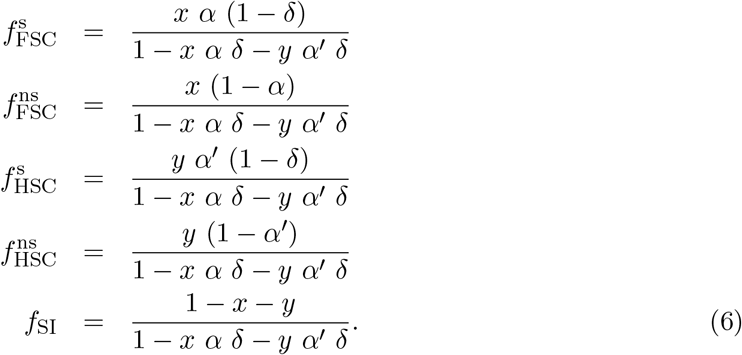

*x, y*, and 1 − *x* − *y* are the population fraction of FSC, HSC, and SI female plants in the current generation, *α*, and *α*′ are the proportions of self-pollen received by FSC and HSC maternal plants, respectively, and *δ* is the proportion of self-fertilization offspring that do not survive. Here, *α*′ = (*α/*2)*/*(1 − *α/*2). Full derivation of 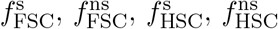,and *f*_SI_ is provided in the supplementary information file. Note: The calculation of the above frequencies is based on the assumption that female fertilization is guaranteed.
3. **Offspring formation**: Divide the diploid maternal plants into three sets: FSC, HSC, or SI. With the probabilities 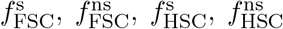,and *f*_SI_ calculated in the previous step, pick a diploid maternal plant to reproduce as a female, from the three sets and pick the female’s mode of reproduction (self or non-self). For example, with probability 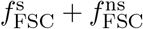 the female will be chosen from the FSC set and with probability *α* it will self-fertilize (otherwise it will be outcrossed), etc.

If the chosen female plant H_2*i*−1_H_2*i*_ is FSC and chosen to self-fertilize (with probability 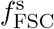), the offspring will have genotype H_2*i*−1_H_2*i*−1_ or H_2*i*−1_H_2*i*_, or H_2*i*_H_2*i*_ with probabilities 0.25, 0.5, and 0.25 respectively. If the chosen female plant H_2*i*−1_H_2*i*_ is HSC and chosen to self-fertilize (with probability *f* ^s^), the offspring will have genotype H_2*i*−1_H_2*i*−1_ or H_2*i*−1_H_2*i*_ in equal proportions, considering that only H_2*i*−1_ S-haplotype can fertilize the female plant, but not H_2*i*_. If the female plant H_2*i*−1_H_2*i*_ is chosen to outcross (whether it is FSC, HSC or SI) it is fertilized by cross-pollen. In this case, it is given *k* attempts to match a randomly chosen non-self S-haplotype H_*j*_, as a male. Once a match is found, the offspring will have genotype H_2*i*−1_H_*j*_ or H_2*i*_H_*j*_ in equal proportions.

Repeat this step until *N* offspring are formed.

Steps 1-3 are a single generation. Repeat these steps until the desired number of generations is reached.

### Model-I: sequence mutation and RNase inactivation mutation

1. **Mutation**: In addition to the sequence mutations, as in the main model, each RNase can alternate between active and inactive states with probability *µ*_R_ per generation. An S-haplotype with an inactive RNase is compatible as a female with any S-haplotype as a male, irrespective of its SLF sequences. That also includes compatibility with self-pollen. In this model, an S-haplotype is considered self-compatible if its RNase is inactive or if it is equipped with an SLF that detoxifies its self-RNase.
2. **Female plant picking**: Same as in the section “Female plant picking”.
3. **Offspring formation**: The fertilization scheme is the same as ‘Offspring formation’ in the main model. Note, however, that an S-haplotype with inactive RNase is compatible as a female with any other S-haplotype as a male, regardless of the female’s RNase and male SLF sequences.

Repeat this step until *N* offspring are formed.

Steps 1-3 are a single generation. Repeat these steps until the desired number of generations is reached.

### Continuous match probability

In the main model, the match between RNase and SLF proteins is deterministic with only two options: match if *E*^*L*^ (*R*_*i*_, *F*_*j*_) *< E*_th_, and no match otherwise. We have generalized this model to incorporate a probabilistic match, given by 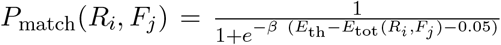, providing a smooth step function, where *β* is the step steepness parameter, taken to be 3 in the simulation.

As the match is probabilistic, we modified the reproduction algorithm accordingly. To test whether a pollen can detoxify an RNase *R*_*i*_, we now determine the total interaction energy (*E*_tot_), and accordingly the match probability *P*_match_(*R*_*i*_, *F*_*j*_) between the RNase *R*_*i*_ and each of the pollen SLFs *F*_*j*_. For each SLF *F*_*j*_, we toss a coin with probability *P*_match_(*R*_*i*_, *F*_*j*_) to determine whether it detoxifies the RNase. If successful, detoxification occurs. If unsuccessful, we proceed to the next SLF, *F*_*j*+1_, until the last one. If none is successful, the pollen does not detoxify the RNase. To determine whether a pollen can produce offspring with a diploid maternal plant, we test both female RNases in this way. Only if both are successfully detoxified does an offspring form.

In contrast to the basic model in which an S-haplotype is either SI or SC, here it is assigned a continuous ‘SI-value’ in the range [0,1]. The SI-value of an S-haplotype with RNase *R* and SLFs *F*_1_, …, *F*_*n*_ equals:

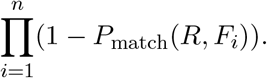

The basic model is the special case in which *P*_match_ equals either 0 or 1. An S-haplotype is then SI if none of the SLFs match, namely ∀*i, P*_match_(*R, F*_*i*_)) = 0, and it is sufficient that a single SLF *F*_*j*_ matches to render the S-haplotype SC. Instead of calculating the SI population proportion, we now average over the population’s SI-values. Because individuals are no longer purely SI or SC, we do not employ the computationally more efficient female picking scheme of Eqs. (6).

### Simulation run-time

We simulated the model for 1 × 10^5^ generations (unless otherwise specified).

## Data analysis

### Determining compatibility classes

#### Binary match between proteins

We classified the population’s S-haplotypes according to their compatibility phenotype. The classification should obey the following two requirements: a S-haplotype affiliated with a class should not be compatible with any of the other members of the same class, either as male or as female, but should be compatible with all members of all other classes, with which it is not affiliated. Following this definition, it is not guaranteed that all S-haplotypes are classified into any of the classes. Specifically, self-compatible S-haplotypes cannot be associated with any class, because they can fertilize their own class members. The division of S-haplotypes into classes is not unique. To ensure that the most frequent S-haplotypes are classified, we first ordered all self-incompatible S-haplotypes in the population by descending copy number. We then defined the most frequent S-haplotype to be the first class. Then, for every unclassified S-haplotype in turn (in descending copy-number order), we tested its compatibility with the existing classes. If it was bidirectionally incompatible with all members of exactly one class and bidirectionally compatible with all members of all the remaining classes, we associated it with the class it was incompatible with. If it was bidirectionally compatible with all members of all existing classes, we defined a new class and associated this S-haplotype with that class. Otherwise, this S-haplotype remained unclassified, and we moved on to the next one. Importantly, our class definition refers only to the compatibility phenotype of S-haplotypes, rather than to their genotype. As our model allows for multiple partners per RNase/SLF, classes could be genetically heterogeneous. Classification was applied every 25 generations, always independently of previous classifications. The full classification algorithm is provided in the supplementary information.

### Effective number of compatibility classes

Since class sizes are often unequal, we use the ‘effective number of classes’, which gives different weights to classes of different sizes. It is defined as:

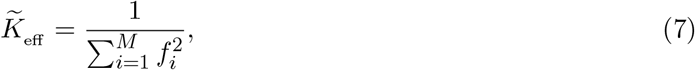

where *f*_*i*_ is the fraction of S-haplotypes associated with the *i*-th class, and *M* is the total number of distinct classes. These fractions are calculated out of the classified S-haplotypes, rather than out of the entire population (that includes unclassified S-haplotypes as well), thus _*i*_ *f*_*i*_ = 1. As the effective number of classes could vary over time, we use in the text its mean 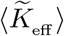 and standard deviation 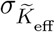 calculated over multiple time points and multiple runs.

#### Direct calculation of the effective number of classes

Alternatively, we determined the effective number of classes directly, without having to classify them first, using the following formula:

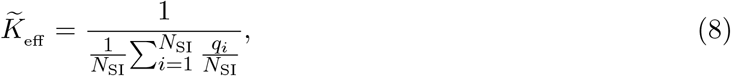

where *N*_SI_ is the number of SI S-haplotypes at a given generation, and *q*_*i*_ represents the number of SI S-haplotypes that are incompatible with S-haplotype *i* both as male and as female. This calculation is significantly faster than the previous one because it does not require applying the population classification first. This calculation can also be applied to the case of continuous match probability by summing over the entire population, and weighing each S-haplotype by its SI-value. Instead of counting the number of S-haplotypes incompatible with *i*, sum over the products of match probabilities: S-haplotype *i* as female with S-haplotype *j* as male times the reciprocal match of S-haplotype *j* as female with S-haplotype *i* as male, namely 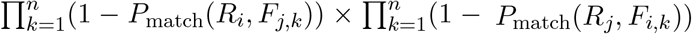.

### The phase diagram

In Fig. 3, for each set of *E*_th_ and *δ* values, we ran a total of four independent simulations, each run for up to 10^5^ generations, of which we discarded the first 8.5 × 10^4^ generations. In the part analyzed, we calculated the population fractions of SC and SI S-haplotypes every 25 generations. For every *E*_th_ and *δ* combination, we combined all data points from the four independent runs and calculated the average fraction of SC and SI S-haplotypes. If the fraction of SI S-haplotypes *<* 5% of the population, such that the diploid genotype proportions were ≈ 0% SI-SI genotype, and 10% SC-SI genotype, the population was considered to be in the self-compatible phase. If the SI S-haplotype fraction was ≥ 5%, the system was considered to be in the mixed phase. Alternatively, if the SI fraction was ≥ 90%, the population was considered to be in the self-incompatible phase. In the phase diagram, *E*_th_ varied from −10 to −2 with a step size of Δ*E*_th_ = 0.25, and *δ* varied from 0.75 to 1.0 with a step size of Δ*δ* = 0.01. The class numbers denoted on the right-side y-axis in Fig. 3a are the average class numbers calculated at *δ* = 1.0, and rounded to the closest integer value. Additional phase diagrams with different parameter values are provided in the supplementary materials.

### Fitting the simulation data to the analytically calculated boundaries

We ran the simulation up to 1 × 10^5^ generations for each combination of *δ* and *E*_th_ values. To obtain the boundary between the SC and mixed phases for a given fraction of HSC genotypes *x* = 0.1 with negligible self-incompatible genotypes, we discarded the first 50 × 10^3^ generations, and took into account the remaining generations with a 25-generation interval between consecutive time points from each run. In the simulation, we checked for which value of *δ*, we obtained *x* = 0.1, and observed that to be *δ*^∗^=0.87 (black dots in Fig. 4c) which is very close to the analytically calculated value *δ*^∗^ = 0.868, Eq. (3) (black dashed line in Fig. 4c). For the calculation of the boundary between the mixed and SI phases. First, we checked for which combination of (*δ, E*_th_), the proportion of FSC genotypes is ≈ 0.01. From that, we took the average class number obtained from the simulation for each (*δ, E*_th_), and used Eq. 5 to calculate *δ*^∗∗^ – shown as red dots in Fig. 4c. Note: Each combination of (*δ, E*_th_) is associated with a distribution of effective class numbers, not just a single value – see Fig. 4e.

### Unbiased autocorrelation

The unbiased temporal correlation *γ*(Δ*t*) of the SI population proportion *r*(*t*), for a given set of parameters within the mixed phase, was calculated as below,

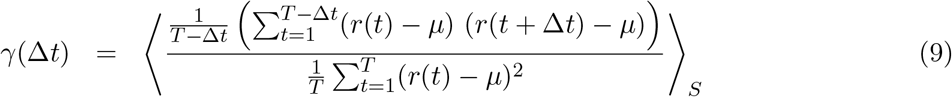

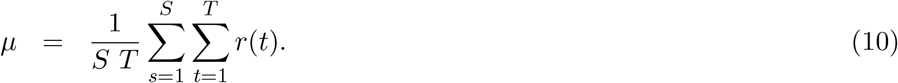

Here *µ* is the average value of the SI population proportion over all the data points lumped together from all the independent runs. *T* = 2000 is the total number of points in each time series used, *S* = 40 is the number of independent runs, and ⟨·⟨_*S*_ is the average over the independent runs.

### Calculation of male-compatibility between SC and SI S-haplotypes in the mixed phase

The average male-compatibilities between SI and SC sub-populations (both ways), as well as amongst the SC sub-population, were calculated using simulation data obtained with parameter values *E*_th_ = −7, *α* = 0.60, and *δ* = 0.92, while the first 75,000 generations of the simulations were discarded. The male-compatibility distributions are shown in Figs. S1, and S2.

## Data availability

The authors declare that all data supporting the findings of this study are available within the article and its Supplementary Information file.

## Code availability

The code is available on gitHub repository: https://github.com/Tamar-Friedlander-Lab/CNSR-based-SI-system-phase-diagram.git.

