## Supplementary_information for "Volatile but persistent co-existence of self-compatibility and self-incompatibility in plants"

[1]The Robert H. Smith Institute of Plant Sciences and Genetics in Agriculture
Faculty of Agriculture, The Hebrew University of Jerusalem,
P.O. Box 12 Rehovot 7610001, Israel

[2]The Einstein Institute of Mathematics, Faculty of Natural Sciences,
The Hebrew University of Jerusalem, Jerusalem 9190401, Israel.

**Supplementary information file contains the following,**

1. **Four biochemical categories and their equilibrium frequencies**

   1.1 Classification of 20 amino acids into four biochemical categories.

   1.2 Equilibrium frequencies of four biochemical categories of amino acids.

2. **Algorithm for compatibility classes**

   2.1 Algorithm to classify the S-haplotypes into the compatibility classes.

3. **Diploid model**

   Supplementary text

   3.1 Probability of choosing a female plant as self-pollinated full self-compatible (FSC), out-crossed full self-compatible, self-pollinated half self-compatible (HSC), out-crossed half self-compatible, and self-incompatible.

   3.2 Supplementary figures.

   S1. Distributions of male-compatibilities between SI and SC sub-populations in the mixed phase vary with the number of SI classes.

   S2. Compatibility distribution of SI (male/female) and SC (female/male) for an individual haplotype when a SC can occur due to sequence mutations only.

   S3. Compatibility genotype map of SI, half SC (HSC), and full SC (FSC) for the main model.

   S4. Class number and self-incompatible population fraction maps when both sequence and RNase inactivation mutations can cause self-compatibility.

   S5. The number of classes exhibits a phase transition in the $\delta$-$E_{\text{th}}$ plane for $2N = 2000$ with a shift in the phase boundaries.

   S6. The number of classes exhibits a phase transition in the $\delta$-$E_{\text{th}}$ plane for $\mu' = \mu/5$ with a shift in the phase boundaries.

   S7. The number of classes exhibits a phase transition in the $\delta$-$E_{\text{th}}$ plane for $\mu' = 2 \cdot \mu$ with no shift in the phase boundaries.

   S8. The mixed phase shrinks when selfing is high, $\alpha = 0.90$, while the number of non-self fertilization attempts per female remains high.

   S9. Starting from a fully self-compatible population, most of the phase diagram is retained, except for high $E_{\text{th}}$, where no SI phase emerges.

   S10. Recovery of SI phase: Introducing a few SI haplotypes, when the model starts with SC haplotypes, the population quickly recovers to the SI phase.

   S11. The proportion of S-haplotypes that are not affiliated with any of the compatibility classes.

   S12. The mixed phase persists for a long time, starting from an initially self-incompatible population.

   S13. The mixed phase persists for a long time, starting from an initially self-compatible population.

   S14. The mixed phase is significantly more fluctuating compared to the full SI phase.

   S15. Applying a continuous match probability between RNase and SLFs yields a qualitatively similar phase diagram to the main model.

   S16. The phase diagram of mating regimes projected on the ($\delta$,$\alpha$) parameter plane.

   S17. The fitness values of SI and SC S-haplotypes in the mixed phase are stable and highly similar, despite large-amplitude temporal fluctuations in the SI population proportion.

   S18. Distributions of the population SI fraction under different parameter combinations in the phase plane.

4. **Haploid model**

   Supplementary text

4.1 Probability of choosing a female S-haplotype as self-pollinated, self-compatible, out-crossed self-compatible, and self-incompatible.

4.2 Analytical calculation of boundaries between SC - mixed phase, and mixed - SI phase.

4.3 The stochastic simulation.

4.4 The phase diagram.

4.5 Fitting the simulation data to the fitness equations.

4.6 Supplementary figures.

S19. Model description.

S20. The population life cycle is implemented in a stochastic simulation.

S21. The model exhibits three separate phases: a self-compatible population, a self-incompatible population partitioned into classes, or a mixture of self-compatible and self-incompatible sub-populations.

S22. Self-incompatible population fraction and number of classes map when only sequence mutations can cause self-compatibility.

S23. Distributions of male-compatibilities between SI and SC sub-populations in the mixed phase vary with the number of SI classes.

S24. In most of the parameter range, except for the highest $E_{\text{th}}$, the phase diagram is independent of the population's initial condition: whether it starts as fully self-compatible or fully self-incompatible.

S25. The three different phases: self-compatible, self-incompatible, and mixed, are obtained for various threshold values defining the boundaries between the phases.

S26. The three different phases: self-compatible, self-incompatible, and mixed, are also obtained for different values of the self-pollination rate $\alpha$.

S27. The population proportion of unclassified S-haplotypes decreases, and the proportion of self-compatible ones increases with $E_{\text{th}}$.

S28. Beginning from a fully self-compatible population, self-incompatibility can emerge under a high interaction energy threshold if SI S-haplotypes are manually introduced.

S29. Distribution of population proportion of unclassified SI S-haplotypes in the mixed phase.

S30. There are multiple distinct genotypes in the SC phase.

S31. The mixed phase persists for a long time, starting from an initially self-incompatible population.

S32. The mixed phase persists for a long time, starting from an initially self-compatible population.

S33. Class number and self-incompatible population fraction maps when both sequence and RNase inactivation mutations can cause self-compatibility.

S34. The class number increases, and the class size decreases under pollen limitation.

S35. The population proportion of unclassified S-haplotypes decreases, and the proportion of self-compatible ones increases under pollen limitation.

S36. Both self-compatibility and self-incompatibility population states persist even under pollen limitation, whereas the mixed phase nearly vanishes.

S37. The mixed-phase region becomes broader as the number of fertilization attempts per female, $k$, increases.

### 1. Four biochemical categories of amino acids and their equilibrium frequencies

1.1 Classification of 20 amino acids into four biochemical categories

Table S1: Four biochemical categories of amino acid [1].

| Biochemical categories | H (%) | P (%) | + (%) | - (%) |
|---|---|---|---|---|
| Amino acids | F (3.86) | C (1.37) | E (6.74) | R (5.53) |
| | W (1.09) | Y (2.92) | D (5.46) | K (5.82) |
| | M (2.41) | T (5.35) | | |
| | L (9.65) | S (6.60) | | |
| | I (5.93) | N (4.06) | | |
| | V (6.86) | H (2.27) | | |
| | G (7.08) | Q (3.93) | | |
| | P (4.72) | | | |
| | A (8.26) | | | |

1.2 Equilibrium frequencies of four biochemical categories of amino acids

Table S2: Equilibrium frequencies [2].

| Biochemical categories | H (%) | P (%) | + (%) | - (%) |
|---|---|---|---|---|
| Threshold energy | | | | |
| $E_{\mathrm{th}}$ = -10 | 0.3403 | 0.3567 | 0.1596 | 0.1434 |
| $E_{\mathrm{th}}$ = -9 | 0.3350 | 0.3314 | 0.1616 | 0.1720 |
| $E_{\mathrm{th}}$ = -8 | 0.3046 | 0.2887 | 0.1907 | 0.2161 |
| $E_{\mathrm{th}}$ = -7 | 0.2710 | 0.2950 | 0.2045 | 0.2295 |
| $E_{\mathrm{th}}$ = -6 | 0.2209 | 0.2946 | 0.2290 | 0.2555 |
| $E_{\mathrm{th}}$ = -5 | 0.2031 | 0.2681 | 0.2376 | 0.2912 |
| $E_{\mathrm{th}}$ = -4 | 0.1717 | 0.2576 | 0.2619 | 0.3087 |
| $E_{\mathrm{th}}$ = -3 | 0.1297 | 0.2392 | 0.2985 | 0.3326 |
| $E_{\mathrm{th}}$ = -2 | 0.0958 | 0.2042 | 0.3178 | 0.3822 |

### 2. Algorithm for determination of compatibility classes

We classify the population of S-haplotypes into compatibility classes based on their mutual compatibility property. The classification algorithm fulfills two conditions. First, every S-haplotype associated with a compatibility class is incompatible with all other members, as a male and as a female. Second, every classified S-haplotype is compatible with all the members of all the other classes again, both as a male and as a female. To ensure that the most frequent S-haplotype is classified, we order the S-haplotype in descending order and apply the algorithm starting from the most frequent S-haplotype and ending in the least frequent one. We describe the classification algorithm in detail below.

In the compatibility matrix $C$ the element $C_{[i,j]}$ is 1 if S-haplotype $\mathrm{H}_i$ as a female is compatible with S-haplotype $\mathrm{H}_j$ as a male, otherwise it is 0.

**Notations**:
$range(i,j)$: $(i, i+1 \ldots j)$
$F[1]$: first element of the sequence $F$
$F_{\mathrm{C}}$: $cumsum(F)$, cumulative sum of $F$
$n\{F\}$: number of elements in sequence $F$

**input** : Copy number of all distinct S-haplotypes $F = (\tilde{f}_1 \dots \tilde{f}_n) \; \forall \; \tilde{f}_i > \tilde{f}_{i+1}, i = 1 \dots n-1$.
Corresponding compatibility matrix $C_{n \times n} \; \forall \; C[i, i] = 0$, all S-haplotypes are SI.
$C[\text{female}_i, \text{male}_j] = 1$ $\text{male}_j$ is compatible with $\text{female}_i$
$C[\text{female}_i, \text{male}_j] = 0$ $\text{male}_j$ is incompatible with $\text{female}_i$.

**output:** Compatibility classes with each class having different S-haplotypes.
Unclassified S-haplotypes.

```
class_dict = {1 : [1]}     /* structure: {class id:[indices of all haplotype]};
unclassified_haps = [ ];
for i to range(2, n{F}) do
    current_classes = [class ids of all current classes from class_dict];
    no_of_classes_compatible_with = 0;
    no_of_classes_incompatible_with = 0;
    class_id_incompatible_with = 0;

    for each_class_id to current_classes do
        no_of_haps_compatible_with = 0;
        no_of_haps_incompatible_with = 0;

        /* checking compatibility and incompatibility over all S-haplotypes of a class;
        for hap_ind to class_dict[each_class_id] do
            if C[i, hap_ind] == 1 and C[hap_ind, i] == 1 then
                no_of_haps_compatible_with += 1;
            end
            if C[i, hap_ind] == 0 and C[hap_ind, i] == 0 then
                no_of_haps_incompatible_with += 1;
            end
        end

        if no_of_haps_compatible_with == len(class_dict[each_class_id]) then
            no_of_classes_compatible_with += 1;
        end
        if no_of_haps_incompatible_with == len(class_dict[each_class_id]) then
            no_of_classes_incompatible_with += 1;
            class_id_incompatible_with = each_class_id;
        end
    end

    /* if a new distinct S-haplotype is bidirectionally compatible with all the members of existing
       classes, this haplotype joins a new class;
    if no_of_classes_compatible_with == len(current_classes) then
        class_dict[max(current_classes)+1] = [i];
    else
        /* if a new distinct S-haplotype is bidirectionally compatible with all the members of
           existing n − 1 classes and bidirectionally incompatible with the members of only one class,
           it joins the incompatible class ;
        if no_of_classes_compatible_with == len(current_classes) − 1 and
          no_of_classes_incompatible_with == 1 then
            class_dict[class_id_incompatible_with].append(i);
        else
            /* the new distinct S-haplotype cannot join any of the classes, hence discarded;
            unclassified_haps.append(i);
        end
    end
end
```

### 3. Diploid model

**3.1 Probability of choosing a female plant as self-pollinated full self-compatible (FSC), out-crossed full self-compatible, self-pollinated half self-compatible (HSC), out-crossed half self-compatible, and self-incompatible.**

Consider $x$, $y$, and $z$ are the fractions of FSC, HSC, and SI S-haplotypes at a given generation G, where $x + y + z = 1$. Hence, the probability of choosing a female plant as self-pollinated full self-compatible $f^{\mathrm{s}}_{\mathrm{FSC}}$, out-crossed full self-compatible $f^{\mathrm{ns}}_{\mathrm{FSC}}$, self-pollinated half self-compatible $f^{\mathrm{s}}_{\mathrm{HSC}}$, out-crossed half self-compatible $f^{\mathrm{ns}}_{\mathrm{HSC}}$ and self-incompatible $f_{\mathrm{SI}}$ are as,

$$
\begin{aligned}
f^{\mathrm{s}}_{\mathrm{FSC}} &= x\ \alpha\ (1-\delta) + \overbrace{\underbrace{(x\ \alpha\ \delta + y\ \alpha'\ \delta)}_{\text{dying fraction}}\ \underbrace{x\ \alpha\ (1-\delta)}_{\text{self-pollinated}}}^{\text{round 1}} + \overbrace{\underbrace{(x\ \alpha\ \delta + y\ \alpha'\ \delta)^2}_{\text{dying fraction}}\ \underbrace{x\ \alpha\ (1-\delta)}_{\text{self-pollinated}}}^{\text{round 2}} + \dots \\
f^{\mathrm{ns}}_{\mathrm{FSC}} &= x\ (1-\alpha) + \overbrace{\underbrace{(x\ \alpha\ \delta + y\ \alpha'\ \delta)}_{\text{dying fraction}}\ \underbrace{x\ (1-\alpha)}_{\text{out-crossed}}}^{\text{round 1}} + \overbrace{\underbrace{(x\ \alpha\ \delta + y\ \alpha'\ \delta)^2}_{\text{dying fraction}}\ \underbrace{x\ (1-\alpha)}_{\text{out-crossed}}}^{\text{round 2}} + \dots \\
f^{\mathrm{s}}_{\mathrm{HSC}} &= y\ \alpha'\ (1-\delta) + \overbrace{\underbrace{(x\ \alpha\ \delta + y\ \alpha'\ \delta)}_{\text{dying fraction}}\ \underbrace{y\ \alpha'\ (1-\delta)}_{\text{self-pollinated}}}^{\text{round 1}} + \overbrace{\underbrace{(x\ \alpha\ \delta + y\ \alpha'\ \delta)^2}_{\text{dying fraction}}\ \underbrace{y\ \alpha'\ (1-\delta)}_{\text{self-pollinated}}}^{\text{round 2}} + \dots \\
f^{\mathrm{ns}}_{\mathrm{HSC}} &= y\ (1-\alpha') + \overbrace{\underbrace{(x\ \alpha\ \delta + y\ \alpha'\ \delta)}_{\text{dying fraction}}\ \underbrace{y\ (1-\alpha')}_{\text{out-crossed}}}^{\text{round 1}} + \overbrace{\underbrace{(x\ \alpha\ \delta + y\ \alpha'\ \delta)^2}_{\text{dying fraction}}\ \underbrace{y\ (1-\alpha')}_{\text{out-crossed}}}^{\text{round 2}} + \dots \\
f_{\mathrm{SI}} &= z + \overbrace{\underbrace{(x\ \alpha\ \delta + y\ \alpha'\ \delta)}_{\text{dying fraction}}\ \underbrace{y}_{\text{self-incompatible}}}^{\text{round 1}} + \overbrace{\underbrace{(x\ \alpha\ \delta + z\ \alpha'\ \delta)^2}_{\text{dying fraction}}\ \underbrace{z}_{\text{self-incompatible}}}^{\text{round 2}} + \dots
\end{aligned}
$$

The first term in the right-hand side of the equations is the fraction of $f^{\mathrm{s}}_{\mathrm{FSC}}$, $f^{\mathrm{ns}}_{\mathrm{FSC}}$, $f^{\mathrm{s}}_{\mathrm{HSC}}$, $f^{\mathrm{ns}}_{\mathrm{HSC}}$, and $f_{\mathrm{SI}}$. The second term represents the fraction of $f^{\mathrm{s}}_{\mathrm{FSC}}$, $f^{\mathrm{ns}}_{\mathrm{FSC}}$, $f^{\mathrm{s}}_{\mathrm{HSC}}$, $f^{\mathrm{ns}}_{\mathrm{HSC}}$, and $f_{\mathrm{SI}}$ female plant being chosen after $x\ \alpha\ \delta + y\ \alpha'\ \delta$ fraction of SC plant died - could not make up to adulthood due to inbreeding depression. The third term is the same as the second, but the fraction of plants that could not make it to adulthood is $(x\ \alpha\ \delta)^2$. We have assumed that each female plant is definitely fertilized, so that pollen is abundant. Self-pollination rate $\alpha'$ for HSC female plant is given as $\alpha' = (\alpha/2)/(1-\alpha/2)$. These equations lead to,

$$
\begin{aligned}
f^{\mathrm{s}}_{\mathrm{FSC}} &= x\ \alpha\ (1-\delta)\ (1 + (x\ \alpha\ \delta + y\ \alpha'\ \delta) + (x\ \alpha\ \delta + y\ \alpha'\ \delta)^2 + \dots) && (1) \\
f^{\mathrm{ns}}_{\mathrm{FSC}} &= x\ (1-\alpha)\ (1 + (x\ \alpha\ \delta + y\ \alpha'\ \delta) + (x\ \alpha\ \delta + y\ \alpha'\ \delta)^2 + \dots) && (2) \\
f^{\mathrm{s}}_{\mathrm{HSC}} &= y\ \alpha'\ (1-\delta)\ (1 + (x\ \alpha\ \delta + y\ \alpha'\ \delta) + (x\ \alpha\ \delta + y\ \alpha'\ \delta)^2 + \dots) && (3) \\
f^{\mathrm{ns}}_{\mathrm{HSC}} &= y\ (1-\alpha')\ (1 + (x\ \alpha\ \delta + y\ \alpha'\ \delta) + (x\ \alpha\ \delta + y\ \alpha'\ \delta)^2 + \dots) && (4) \\
f_{\mathrm{SI}} &= z\ (1 + (x\ \alpha\ \delta + y\ \alpha'\ \delta) + (x\ \alpha\ \delta + y\ \alpha'\ \delta)^2 + \dots) && (5)
\end{aligned}
$$

The probability of choosing a female plant as $f^{\mathrm{s}}_{\mathrm{FSC}}$, $f^{\mathrm{ns}}_{\mathrm{FSC}}$, $f^{\mathrm{s}}_{\mathrm{HSC}}$, $f^{\mathrm{ns}}_{\mathrm{HSC}}$, and $f_{\mathrm{SI}}$ are given as,

$$
\begin{aligned}
f^{\mathrm{s}}_{\mathrm{FSC}} &= \frac{x\ \alpha\ (1-\delta)}{1 - x\ \alpha\ \delta - y\ \alpha'\ \delta} && (6) \\
f^{\mathrm{ns}}_{\mathrm{FSC}} &= \frac{x\ (1-\alpha)}{1 - x\ \alpha\ \delta - y\ \alpha'\ \delta} && (7) \\
f^{\mathrm{s}}_{\mathrm{HSC}} &= \frac{y\ \alpha'\ (1-\delta)}{1 - x\ \alpha\ \delta - y\ \alpha'\ \delta} && (8) \\
f^{\mathrm{ns}}_{\mathrm{HSC}} &= \frac{y\ (1-\alpha')}{1 - x\ \alpha\ \delta - y\ \alpha'\ \delta} && (9) \\
f_{\mathrm{SI}} &= \frac{1 - x - y}{1 - x\ \alpha\ \delta - y\ \alpha'\ \delta} && (10)
\end{aligned}
$$

#### 3.2 Supplementary figures.

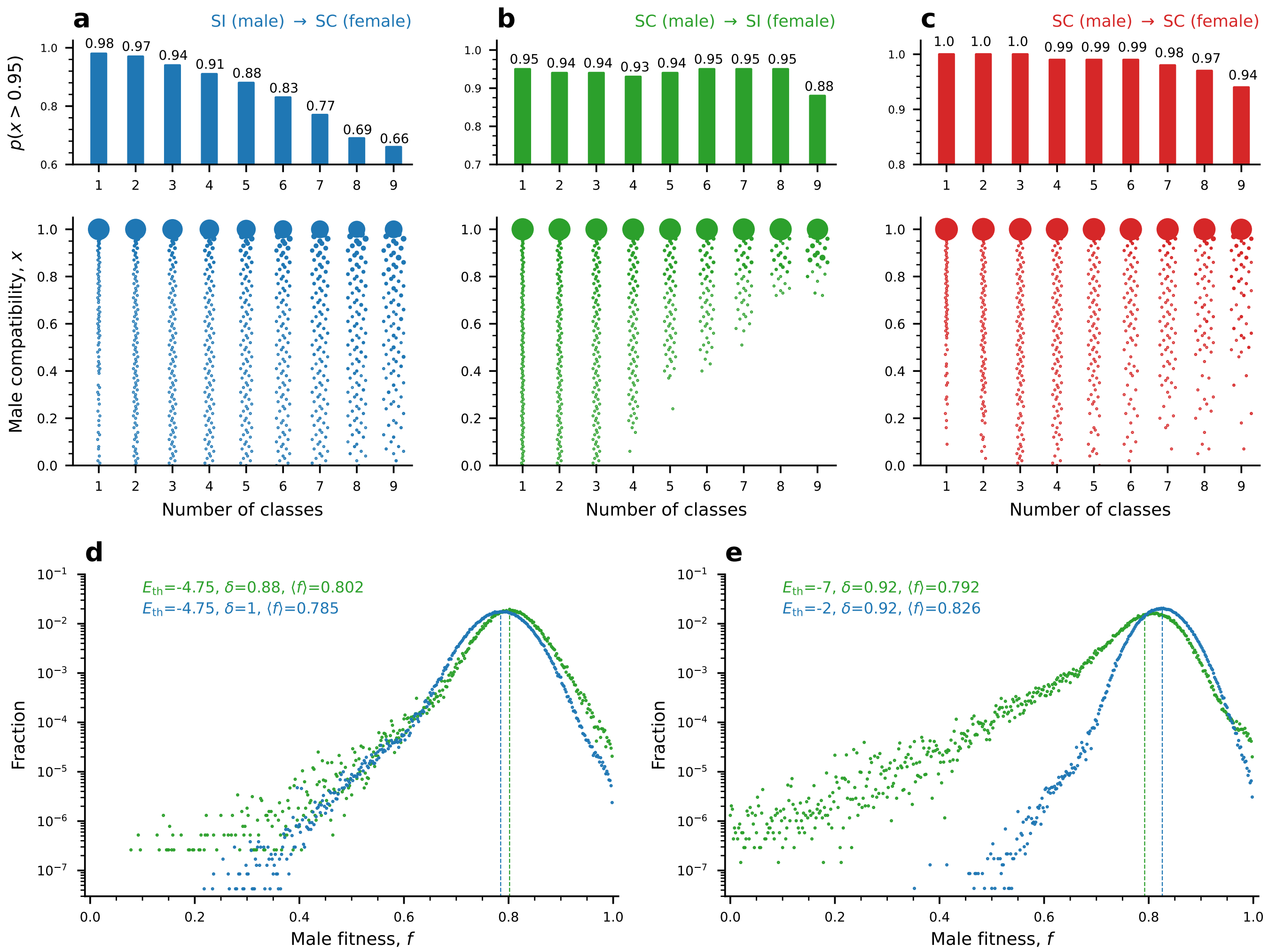

Figure S1: **Distributions of male-compatibilities between SI and SC sub-populations in the mixed phase vary with the number of SI classes.** Distributions of male-compatibilities, namely the population proportion an individual is compatible with as male, between SI as a male and SC as a female **(a)**, SC as a male and SI as a female **(b)**, and SC in both male and female roles **(c)** in the mixed phase, dissected by the number of SI classes in the population (x-axis). Note that the number of classes is also affected by the varying proportion of the SI sub-population. The bar graphs (top) show the frequency of the relevant male haplotypes compatible with at least 0.95 of the target female haplotypes. The scatter plots (bottom) show histograms (vertically aligned) of the compatible proportion. Larger circles represent higher frequencies. Compatibility is highest amongst SC genotypes (c) and lowest between SI as male and SC as female (a). Generally, the lower the compatibility, the higher the number of classes. **(d-e)** The proportion of diploid females (both SI and SC) with which an SI haploid pollen is compatible, for different parameter combinations. The blue and green curves show parameter combinations located in the SI phase and mixed phases, respectively. (d) compares equal $E_{\rm th}$ and different $\delta$ values; (e) compares equal $\delta$ and different $E_{\rm th}$ values. In the latter, the mixed phase has a slightly lower mean compatibility and a broader distribution, with greater representation at lower compatibilities, compared to the SI phase. The figure is based on 128 independent runs, each with 400 data points, and a 25-generation interval between consecutive time points used for the analysis. Total run time is $10^5$. Parameters: $E_{\rm th}$ = -7, $\alpha$ = 0.60, and $\delta$ = 0.92. The values of the remaining parameters are as in Table 2, main text.

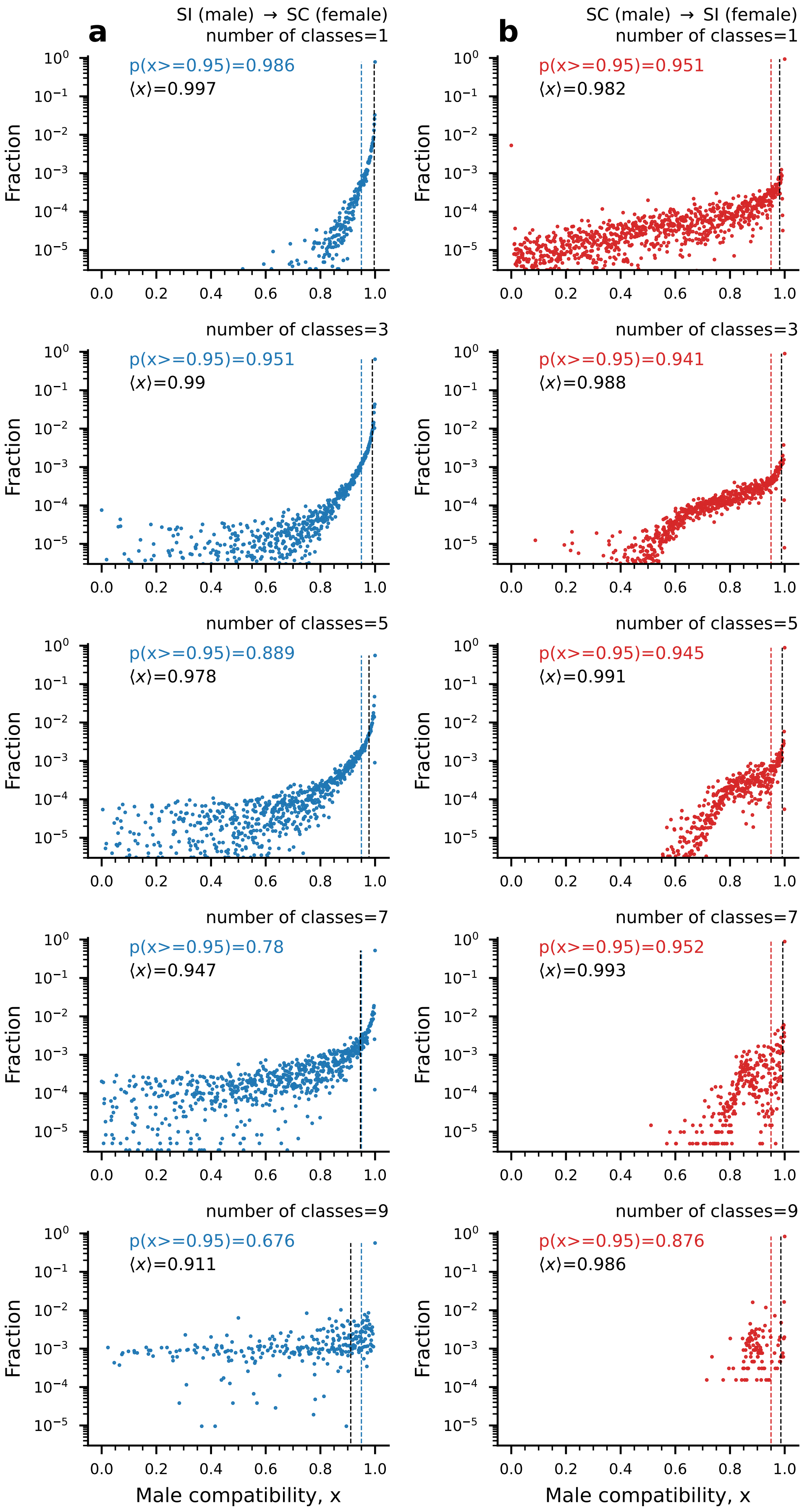
a
b
SI (male) → SC (female)
SC (male) → SI (female)
number of classes=1
p(x>=0.95)=0.986
⟨x⟩=0.997
p(x>=0.95)=0.951
⟨x⟩=0.982
number of classes=3
p(x>=0.95)=0.951
⟨x⟩=0.99
p(x>=0.95)=0.941
⟨x⟩=0.988
number of classes=5
p(x>=0.95)=0.889
⟨x⟩=0.978
p(x>=0.95)=0.945
⟨x⟩=0.991
number of classes=7
p(x>=0.95)=0.78
⟨x⟩=0.947
p(x>=0.95)=0.952
⟨x⟩=0.993
number of classes=9
p(x>=0.95)=0.676
⟨x⟩=0.911
p(x>=0.95)=0.876
⟨x⟩=0.986
Fraction
Male compatibility, x

Figure S2

Figure S2 *(previous page)*: **Compatibility distribution of SI (male/female) and SC (female/male) for an individual S-haplotype when a SC can occur due to sequence mutations only.** The first column shows compatibility between SI (male) and SC (female), and the second shows the reverse. A vertically colored dashed line is the threshold $x$ above which $p(x)$ is calculated. The vertical black dashed line corresponds to the mean male compatibility. SI (male) compatibility reduces as the class number increases because the chances of being selected as male increase as the SI classes increase. Whereas SC male compatibility remains almost the same because these SC haplotypes are just mutants of these SI from different SI classes. All data are calculated over a total of 128 runs, and from each run, we have taken 600 generation points at 25-generation intervals. Each class contains at least 10 haplotypes. Parameters: $E_{\mathrm{th}}$ =-7, and $\delta = 0.92$. The remaining parameter values are as in Table 2 main text. Related to Figs. 3, and 4 in the main text.

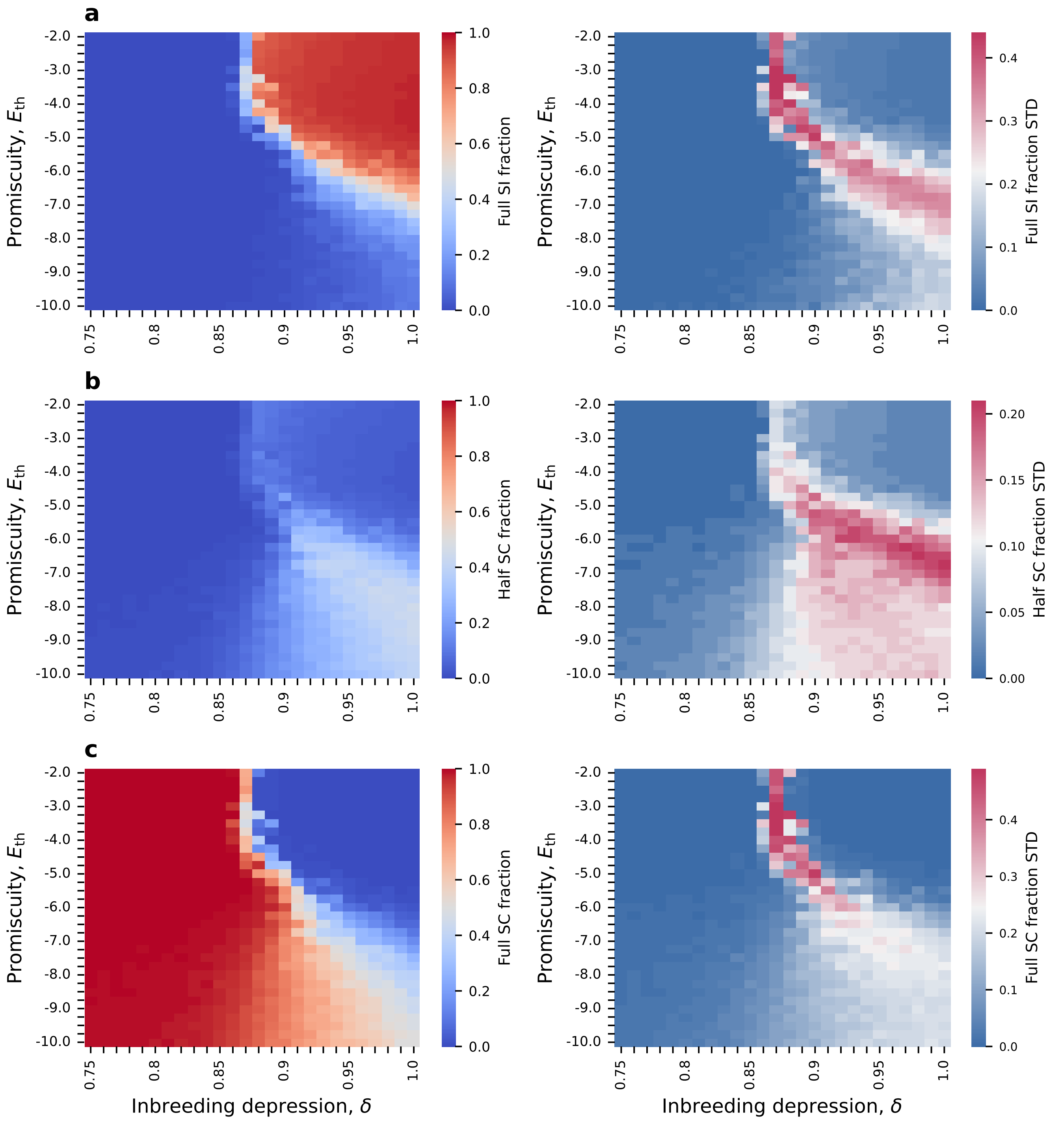

Figure S3: **Frequencies of full SI, half SC (HSC), and full SC (FSC) genotypes as a function of $\delta$ and $E_{\text{th}}$ for the main model.** Mean (left column) and standard deviation (right column) of the population frequencies of full SI (a), half-SI-half-SC (b), and full SC (c) genotypes, in the $\delta$-$E_{\text{th}}$ plane. The SC phase is almost exclusively full SC, the full SI phase is mostly full SI (with a small proportion of HSC, and the mixed phase contains all three genotypes in varying proportions, depending on the parameter values. Even if both S-haplotypes are SC, it is possible that one of them (or both) is unable to self-fertilize due to a lack of compatibility with the RNase of the homologous S-haplotype. We use a 'practical' definition here. For example, a genotype is considered HSC if exactly one of its two S-haplotypes can self-fertilize and the other cannot, etc. The figures are based on simulation results of 4 independent runs for each point in $\delta$-$E_{\text{th}}$ plane for $\delta \in$ [0.75,0.85] and [0.89,1.0], and 30 runs for $\delta \in [0.86, 0.9]$. From each run, we extracted 2000 data points in 25-generation intervals. The parameter values are as in Table 2 main text.

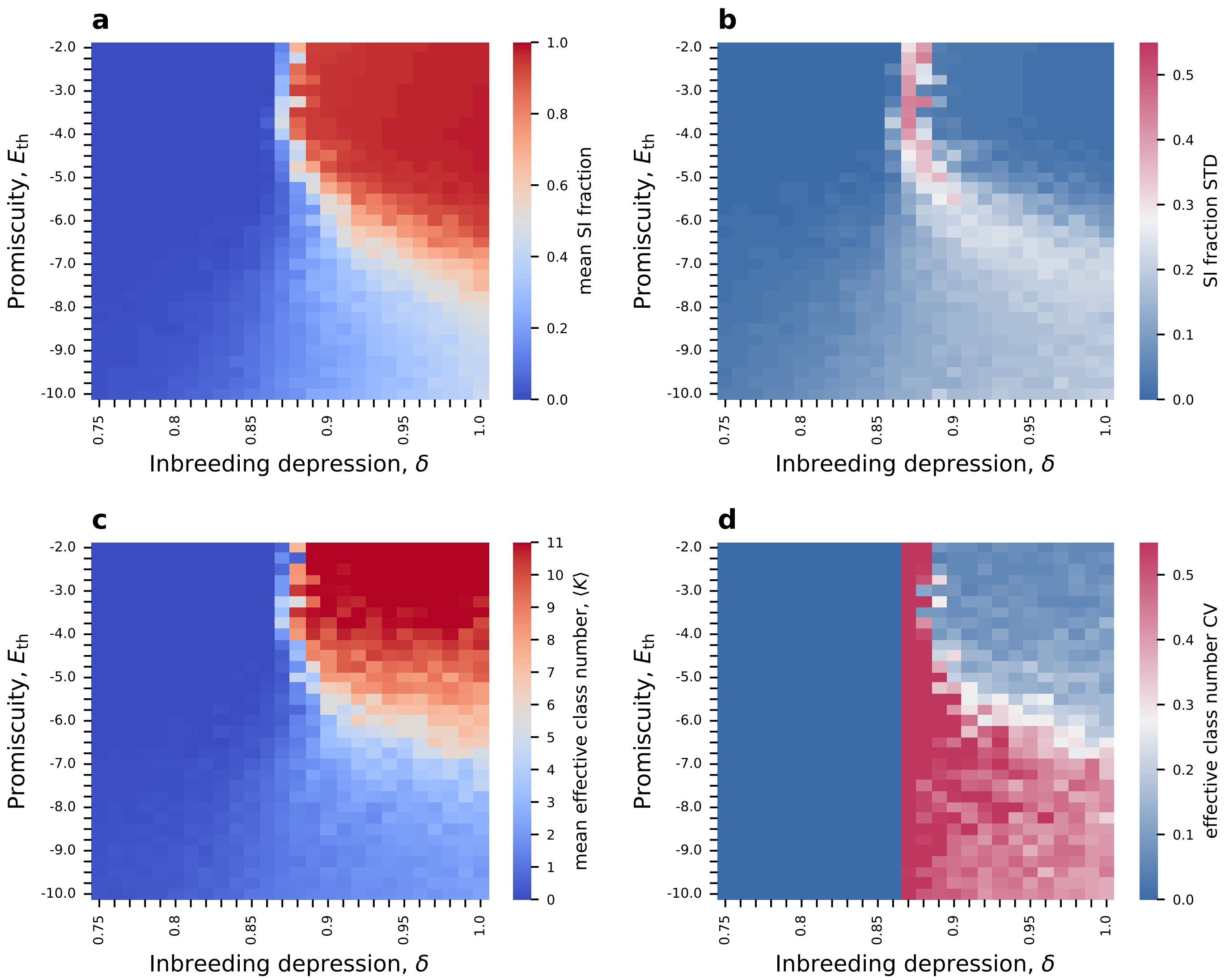

Figure S4: **Class number and self-incompatible population fraction maps when both sequence and RNase inactivation mutations can cause self-compatibility. (a-b)** Maps of the self-incompatible population fraction's mean and standard deviation. **(c-d)** The mean and the CV (std/mean) of the effective number of classes. Here, we observe an unstable mixed phase, similar to the case in which only sequence mutations could cause self-compatibility (compared to Fig. 4 in the main text). Parameter values: $\mu_R = \mu \cdot L$. The figures are based on simulation results of 4 independent runs for $\delta \in$ [0.75,0.85], and [0.89,1.0], and 30 runs for $\delta \in [0.86, 0.88]$. From each run, we have taken 2000 generation points in 25-generation intervals for (a-b) and (c-d), respectively. The minimal class size is 10 S-haplotypes. The remaining parameter values are as in Table 2 main text.

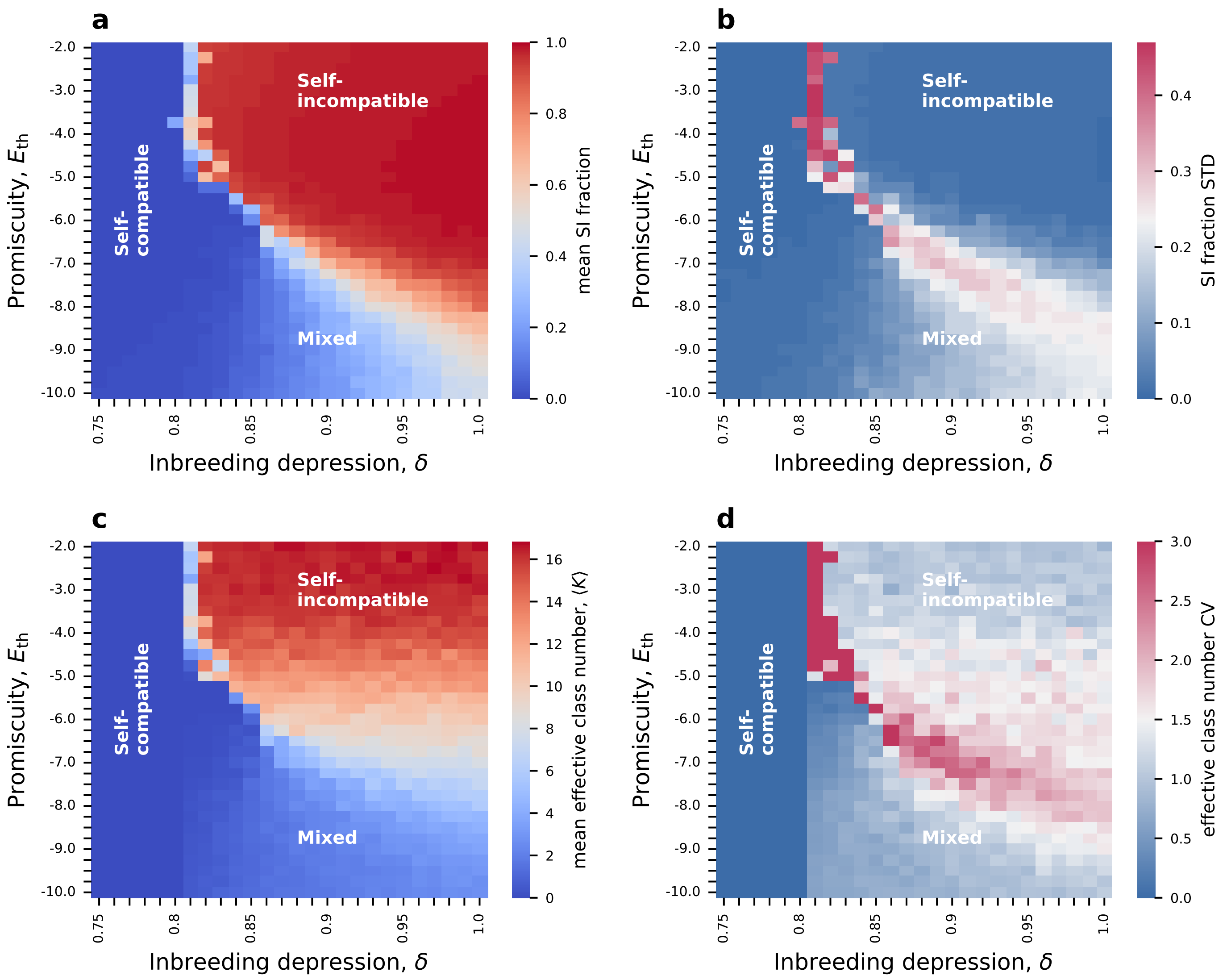

Figure S5: **The number of classes exhibits a phase transition in the $\delta$-$E_{\mathrm{th}}$ plane for $2N = 2000$ with a shift in the phase boundaries. (a-b)** Maps of the self-incompatible population fraction's mean and standard deviation. **(c-d)** The mean and the CV (std/mean) of the effective number of classes. Here we observe the unstable mixed phase, similar to the case for $2N = 1000$ (compared to Fig. 4 in the main text). Parameter values: $2N = 2000, \mu_{\mathrm{R}} = 0$. The figures are based on simulation results of 4 independent runs for $\delta \in$ [0.75,0.80], and [0.83,1.0], and 30 runs for $\delta \in [0.81, 0.82]$ plane. From each run, we extracted 2000 data points in 25-generation intervals. The minimal class size is 10 S-haplotypes. The remaining parameter values are as in Table 2 main text.

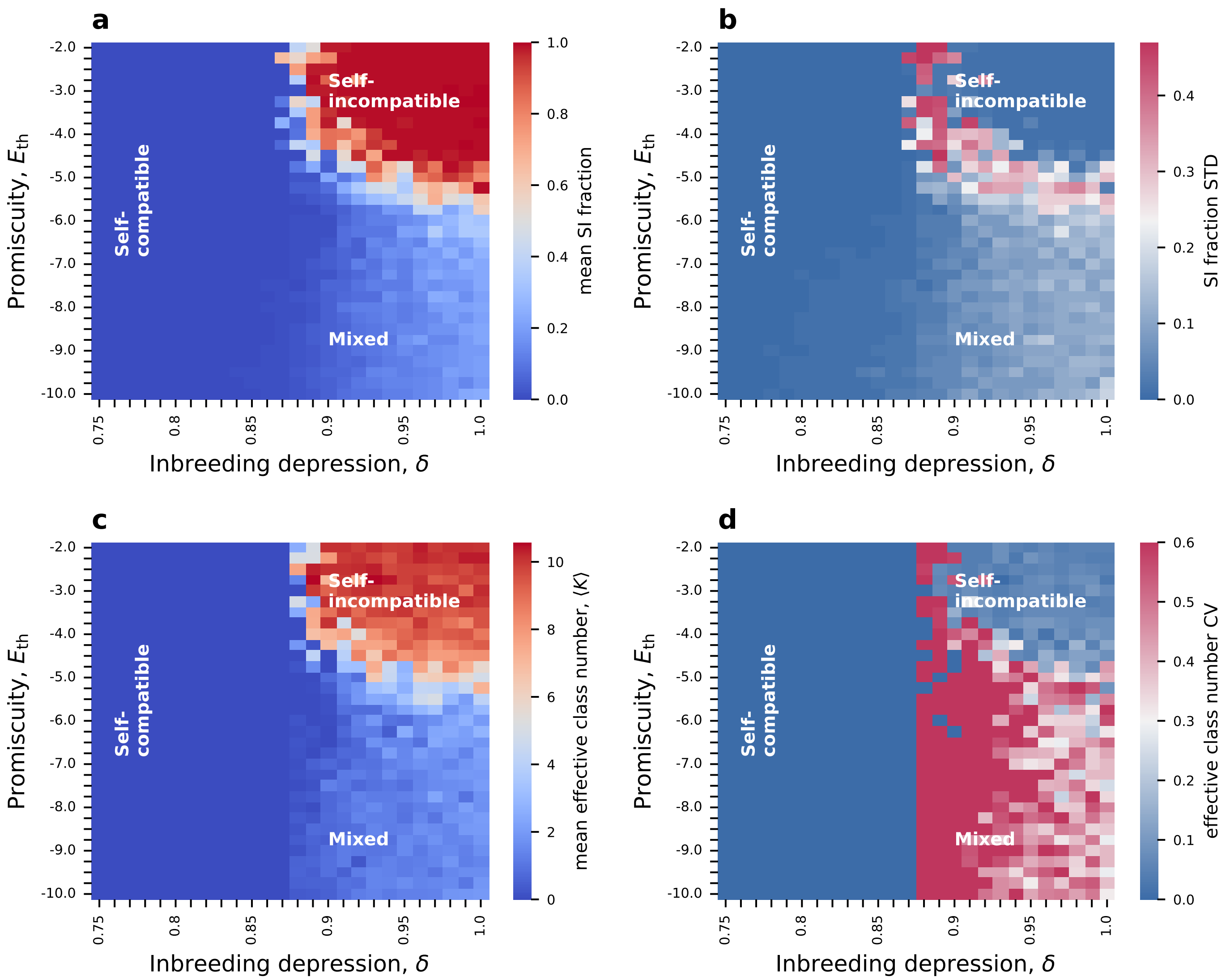

Figure S6: **The number of classes exhibits a phase transition in the $\delta$-$E_{\rm th}$ plane for $\mu' = \mu/5$ with a shift in the phase boundaries. (a-b)** Maps of the self-incompatible population fraction's mean and standard deviation. **(c-d)** The mean and the CV (std/mean) of the effective number of classes. Here we observe the unstable mixed phase, similar to the case for $2N = 1000$ (compared to Fig. 4 in the main text). Parameter values: $\mu' = \mu/5$, $\mu_{\rm R} = 0$. The figures are based on simulation results of 4 independent runs for $\delta \in [0.75, 1.0]$ plane. From each run, we extracted 2000 data points in 25-generation intervals. The minimal class size is 10 S-haplotypes. The remaining parameter values are as in Table 2 main text.

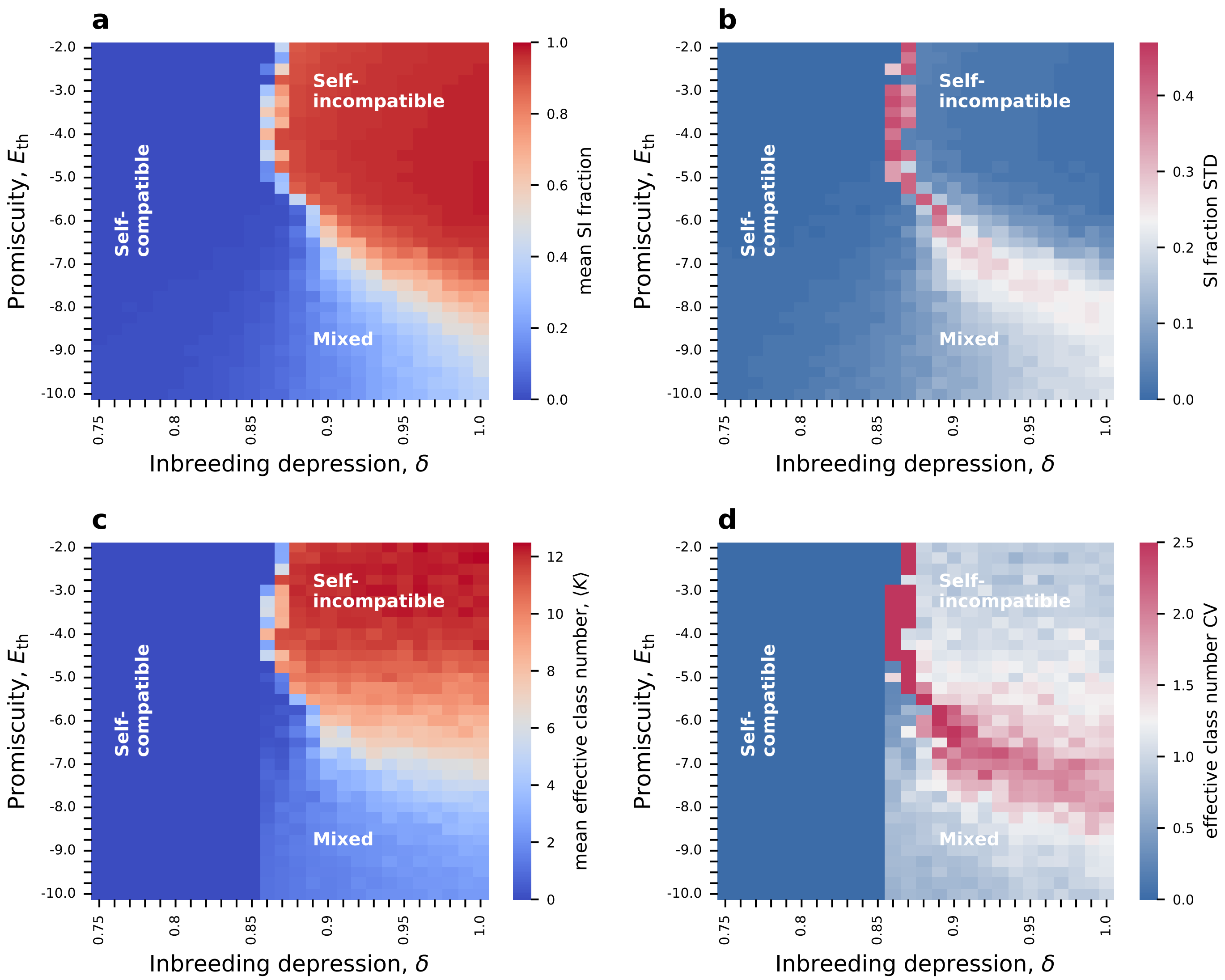

Figure S7: **The number of classes exhibits a phase transition in the $\delta$-$E_{\mathrm{th}}$ plane for $\mu' = 2 \cdot \mu$ with no shift in the phase boundaries.** **(a-b)** Maps of the self-incompatible population fraction's mean and standard deviation. **(c-d)** The mean and the CV (std/mean) of the effective number of classes. Here we observe the unstable mixed phase, similar to the case for $2N = 1000$ (compared to Fig. 4 in the main text). Parameter values: $\mu' = 2\mu, \mu_{\mathrm{R}} = 0$. The figures are based on simulation results of 4 independent runs for $\delta \in [0.75,1.0]$ plabe. From each run, we extracted 2000 data points in 25-generation intervals. The minimal class size is 10 S-haplotypes. The remaining parameter values are as in Table 2 main text.

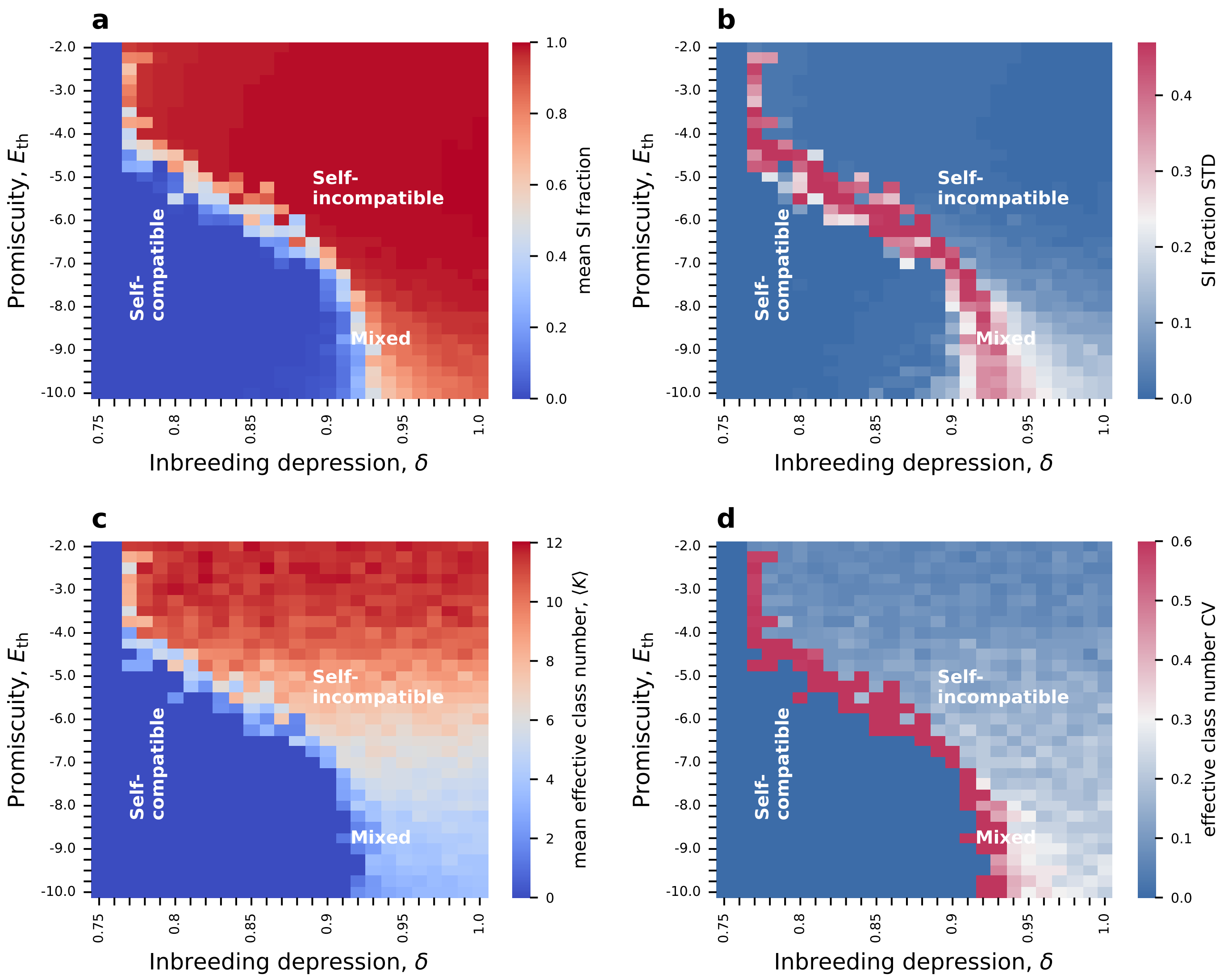

Figure S8: **The mixed phase shrinks when the selfing rate is high, $\alpha = 0.90$, while the number of non-self fertilization attempts per female remains high. (a-b)** Maps of the self-incompatible population fraction's mean and standard deviation. **(c-d)** The mean and the CV (std/mean) of the effective number of classes. Here we observe the unstable mixed phase, but for a very narrow range of $\delta$ values (compared to Fig. 4 in the main text). Parameter values: $\alpha = 0.90$, $k = 1000$ (number of non-self fertilization attempts per female). The figures are based on simulation results from 4 independent runs for $\delta \in [0.75,1.0]$. From each run, we extracted 2000 data points in 25-generation intervals. The minimal class size is 10 S-haplotypes. The remaining parameter values are as in Table 2 main text.

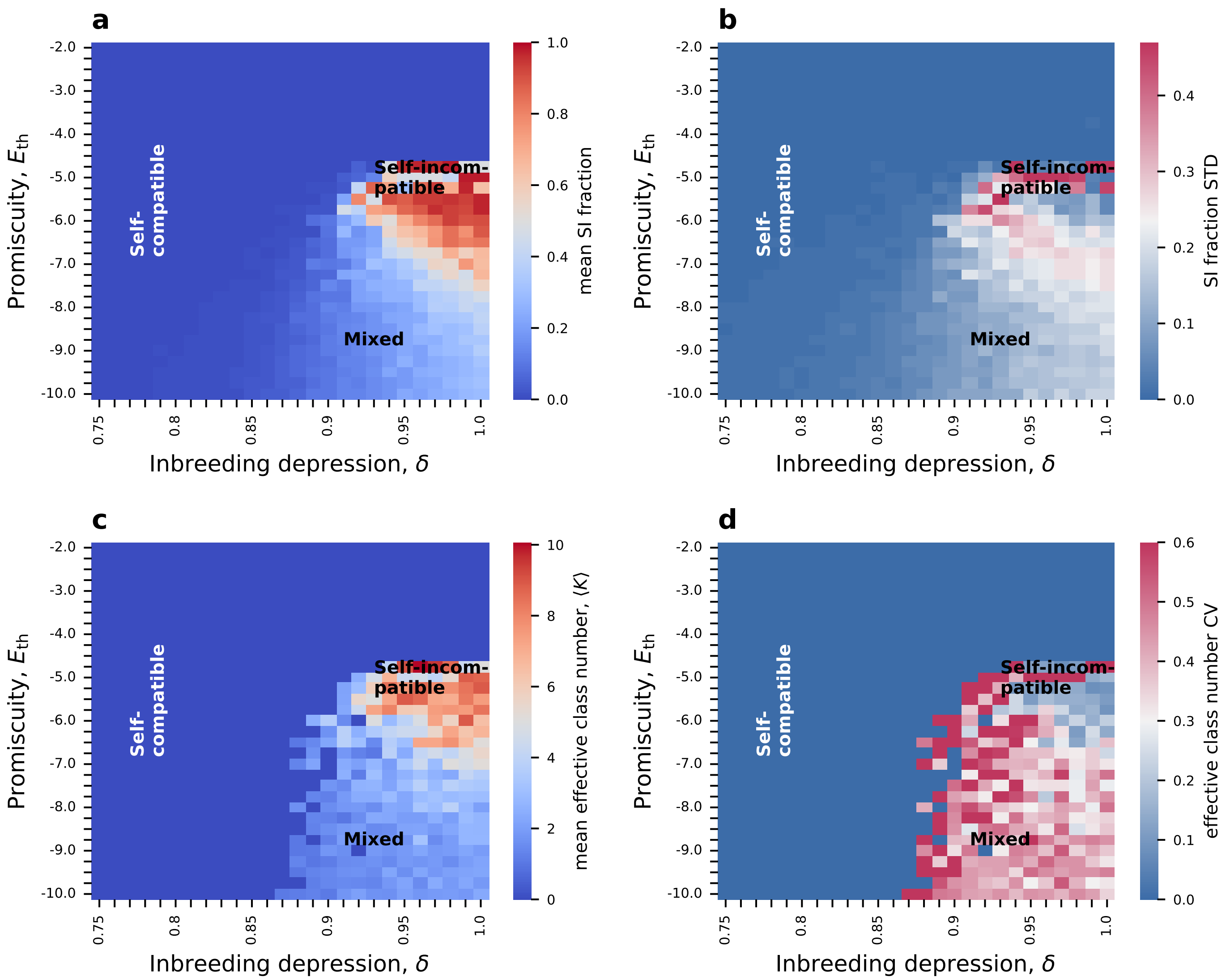

Figure S9: **Starting from a fully self-compatible population, most of the phase diagram is retained, except for high $E_{th}$, where no SI phase emerges.** This figure shows the data shown schematically in 6a in the main text. **(a-b)** Maps of the self-incompatible population fraction mean (a) and standard deviation (b). **(c-d)** The mean and the CV (std/mean) of the effective number of classes. We observe that the SI phase is not recovered for $E_{th} \geq -4.5$, because the waiting time for a mutation that turns an SC S-haplotype to SI is very large. Parameter values: $\alpha = 0.6$. The figures are based on simulation results of 4 independent runs for $\delta \in [0.75,1.0]$. From each run, we extracted 2000 data points in 25-generation intervals. The minimal class size is 10 S-haplotypes. The remaining parameter values are as in Table 2 main text.

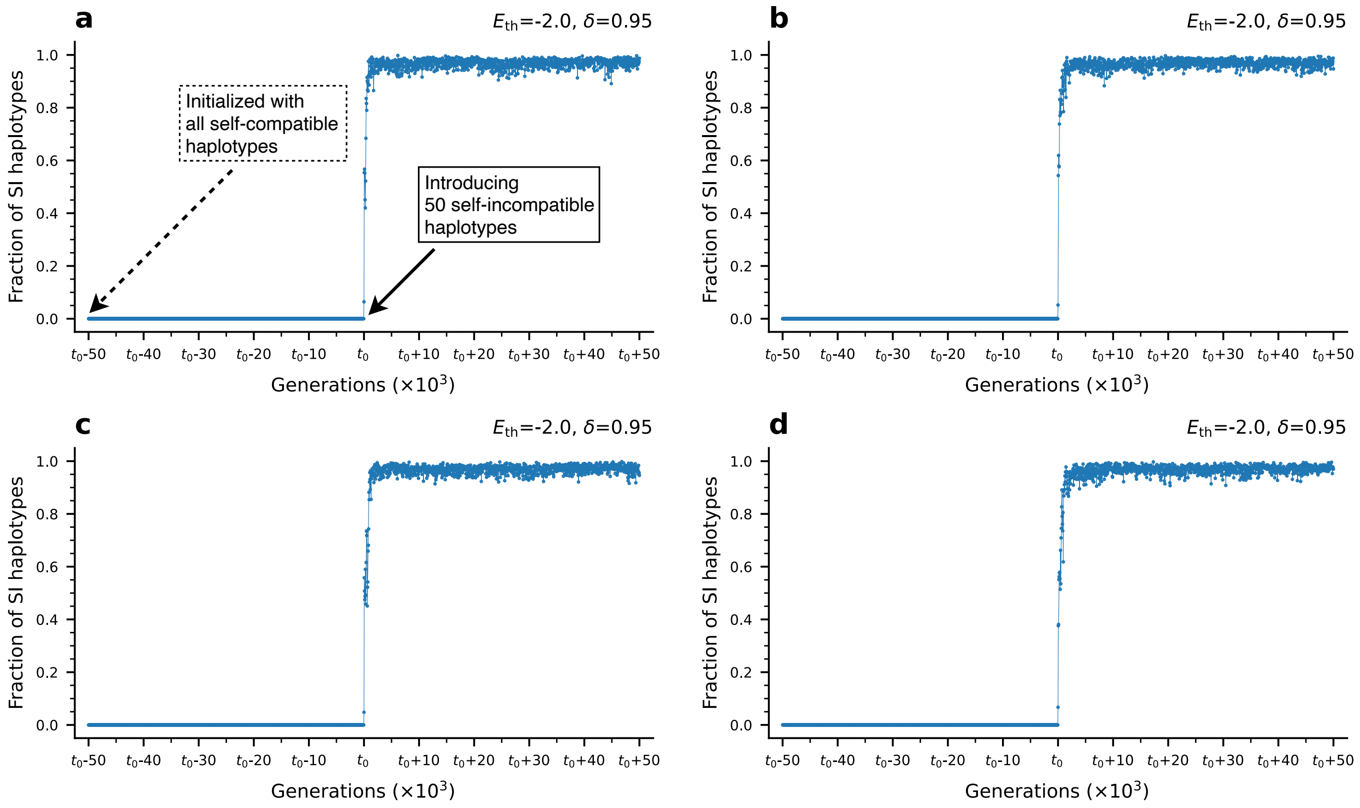

Figure S10: **Recovery of SI phase: Introducing a few SI S-haplotypes, when the population is initialized as fully SC, allows for a quick takeover of the SI population state. (a-d)** The population proportion of SI S-haplotypes in 4 independent simulation runs under $E_{\rm th}$ =-2, $\delta = 0.95$ (parameter values associated with the SI phase) initialized with a fully self-compatible population. Initially, no emergence of self-incompatibility is observed (solid dashed line) after waiting for $5 \cdot 10^4$ generations. SI quickly stabilizes within $5 \cdot 10^3$ generations (solid dotted line) after introducing 50 self-incompatible S-haplotypes into the population at generation $t_0$. Once it emerged, the self-incompatibility population state persists - we have tested that for an additional $5 \cdot 10^4$ generations. This demonstrates that the transition from self-compatibility to self-incompatibility under parameter values supporting the stable existence of self-incompatibility was not observed under high $E_{\rm th}$ only because the probability of mutations converting self-compatible S-haplotypes into self-incompatible ones is extremely low under high $E_{\rm th}$, hence the waiting time for such mutations is infeasibly long. Parameter values: $E_{\rm th} = -2, \delta = 0.95$. The remaining parameters are as in Table 2 main text.

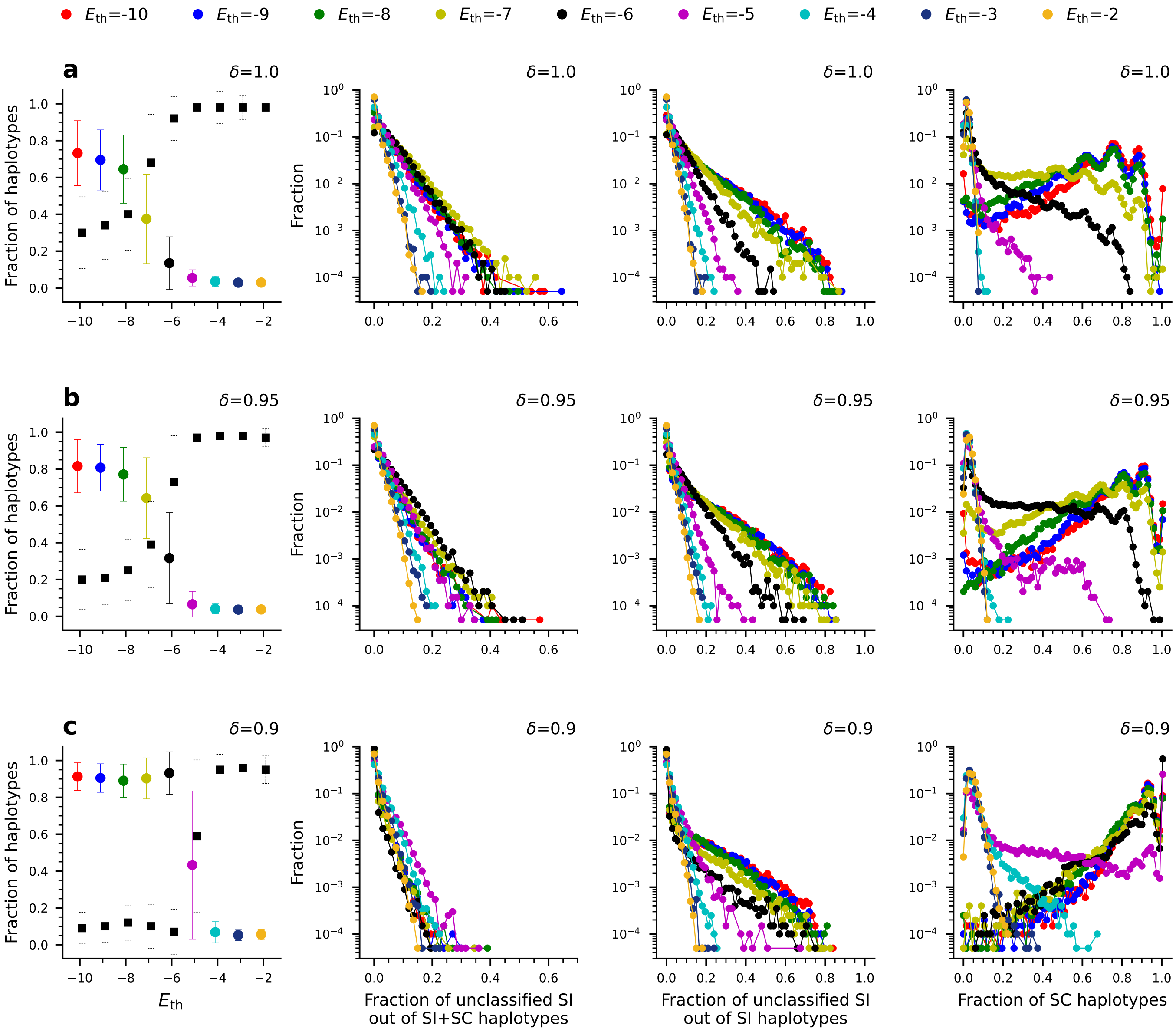

Figure S11: **The proportion of S-haplotypes that are not affiliated with any of the compatibility classes. Leftmost column:** The total proportion of unclassified S-haplotypes (both SI+SC) is shown in colored circles representing the mean value, and bars show the STD. Different colors represent different $E_{\text{th}}$ values, as labeled on top of the figure. Since only SI S-haplotypes can be classified, we also show the population proportion of SI S-haplotypes (both classified and unclassified): Black squares - mean value, bars - the STD. **Second column:** Distributions of the proportion of unclassified S-haplotypes out of the total population of S-haplotypes (SI+SC). We find that the proportion of unclassified S-haplotypes decreases as $E_{\text{th}}$ increases. **Third column:** Distributions of the proportion of unclassified S-haplotypes out of the self-incompatible sub-population of S-haplotypes. Here too, we find that the proportion of unclassified S-haplotypes decreases as $E_{\text{th}}$ increases. **Rightmost column:** Distribution of the proportion of self-compatible S-haplotypes within the entire population. In separate rows, these plots show different values of inbreeding depression, $\delta$.

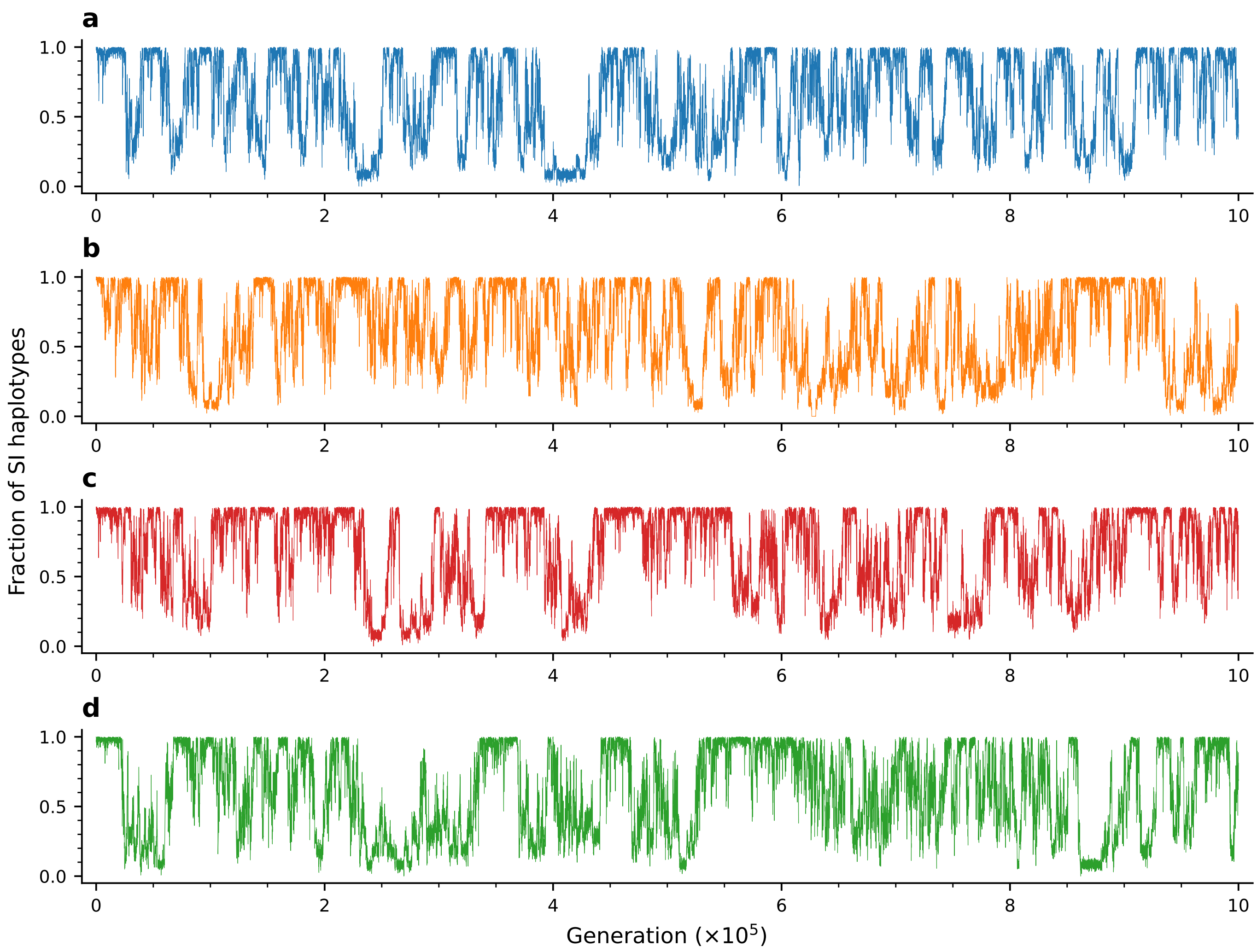

Figure S12: **The mixed phase persists for a long time, starting from an initially self-incompatible population**. We show here time traces from 4 independent runs of the full model under a parameter combination associated with the mixed-phase ($E_{\mathrm{th}} = -6$, $\alpha = 0.6$, and $\delta = 0.94$), each run for $10^6$ generations. All runs were initialized with a fully self-incompatible population. The self-incompatible population proportion is plotted here in 25 generation time intervals. Parameter values are as in Table 2 main text.

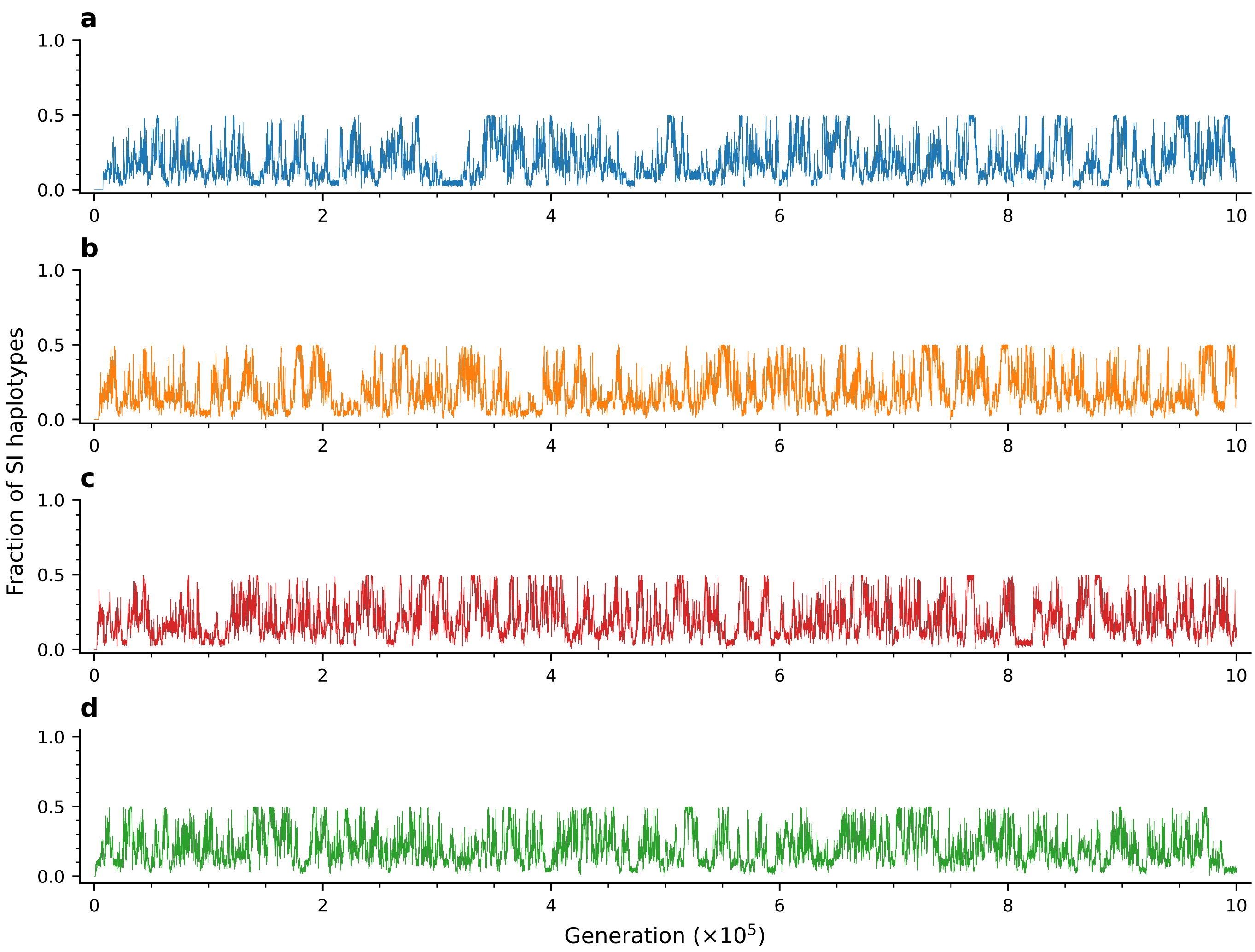

Figure S13: **The mixed phase persists for a long time, starting from an initially self-compatible population**. We show here time traces of 4 independent runs of the full model under a parameter combination associated with the mixed-phase ($E_{\rm th} = -7$, $\alpha = 0.6$ and $\delta = 0.95$), ran up to $10^6$ generations. All runs were initialized with a fully self-compatible population. The self-incompatible population proportion is plotted here in 25 generation time intervals. Parameter values are as in Table 2 main text.

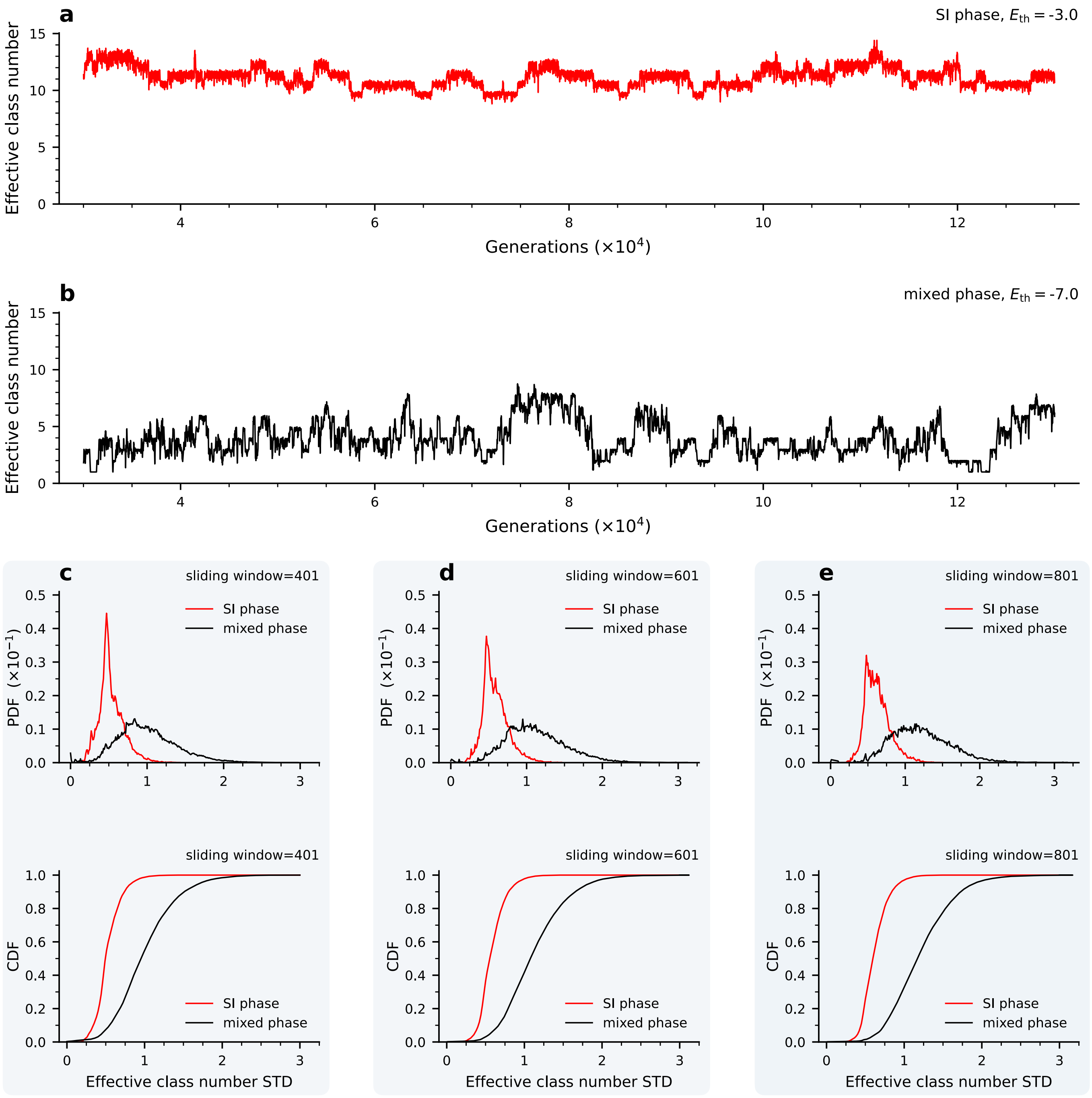

Figure S14: **The mixed phase is significantly more fluctuating compared to the full SI phase.** Examples of time series of the effective number of compatibility classes calculated at 10 generation intervals (as explained in Methods) in the SI phase **(a)** and mixed phase **(b)**. **(c)** We calculated the standard deviation in the effective class number over sliding windows of 401 time points. The class number was calculated every 10 generations, after discarding the initial $3 \times 10^4$ generations, for a total run time of $13 \times 10^4$ generations. The total number of time points used in the calculation is $5 \times 10^5$ from 50 independent runs. Here we illustrate the distributions of these temporal fluctuation sizes, shown as pdf (top) and cdf (bottom). A similar analysis was repeated in different window sizes **(d)** 601 and **(e)** 801 time points, yielding qualitatively similar results. Evidently, the mixed phase exhibits significantly larger temporal fluctuations in class numbers, as seen both in the distributions (c-e) and demonstrated in the time traces (a-b). Energy thresholds for SI (red) and mixed phase (black) are -3 and -7, respectively. $\delta$=0.97, and $\alpha$=0.6 are the same for both phases. Remaining parameters are the same as in Table 2 main text.

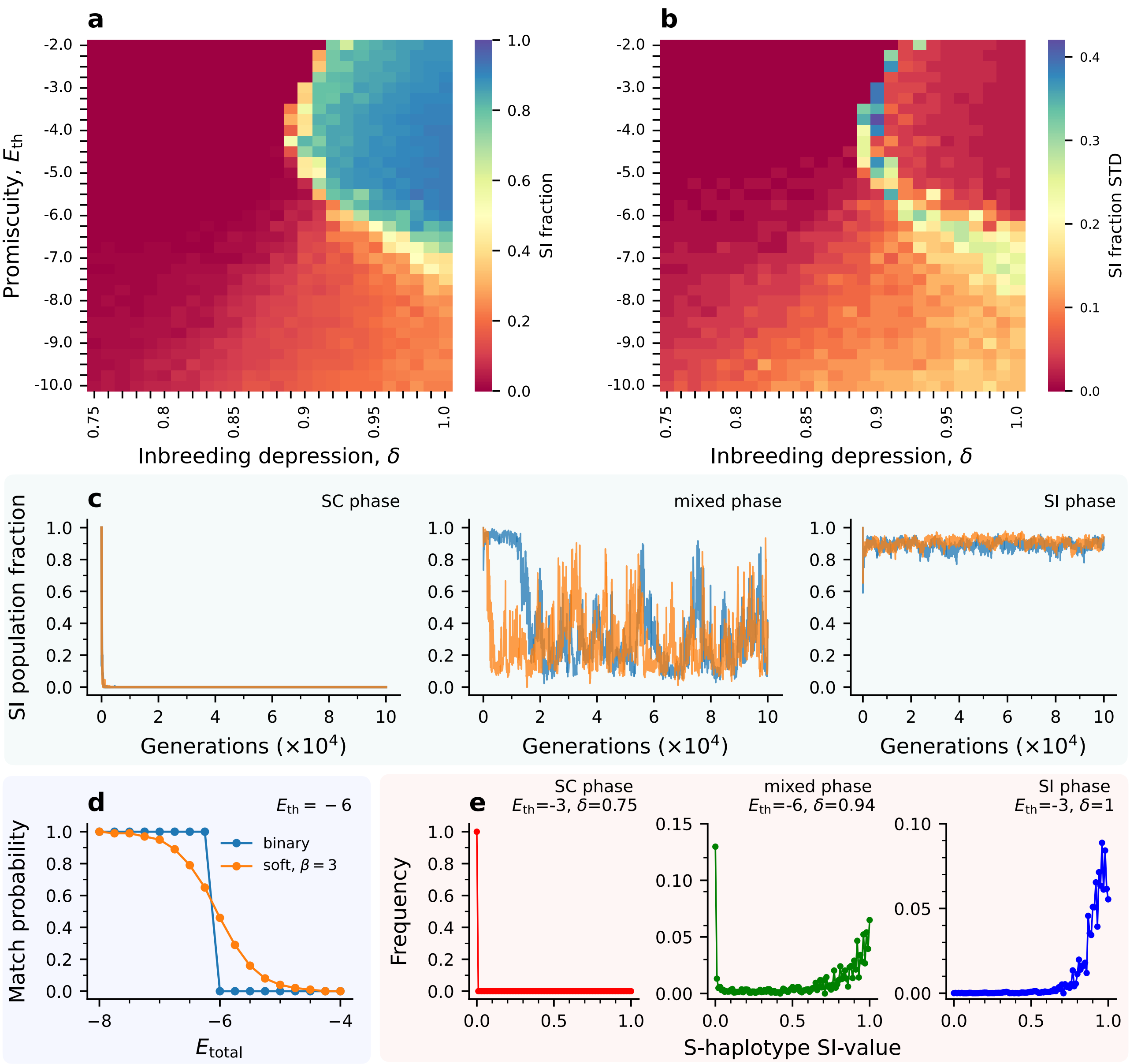

Figure S15: **Applying a continuous match probability between RNases and SLFs yields a qualitatively similar phase diagram to the main model.** Here, the compatibility between two proteins is probabilistic, rather than deterministic, as in the main model. A pair of RNase and SLF proteins match with probability $P_{\text{match}} = \frac{1}{1+e^{-\beta\ (E_{\text{th}}-E_{\text{tot}}-0.05)}}$, where $\beta$ was taken to be 3. See the Methods section for details of the reproduction algorithm in this case. **(a-b)** Maps of the self-incompatible population fraction's mean and standard deviation. We observe the three phases, including the volatile mixed phase, as in the main model (compare with Fig. 4 in the main text). **(c)** Examples of time-series of SI population fraction in the three mating regimes, with two independent runs shown for each. The SI fraction was calculated by summing the 'SI-value' of all S-haplotypes in the population. Similar to the main model, the SC and SI regimes show stable SI fractions (0 or 1, respectively) with only little fluctuation around this value in the SI case. The mixed phase again shows large temporal fluctuations spanning most of the range between full SI and full SC. **(d)** The binary and the continuous match probabilities for $\beta = 3$ and $E_{\text{th}} = -6$ are shown. **(e)** Unlike the main model, in which an S-haplotype is either SI or SC, here this trait can take any value in the range [0,1], which we call the 'SI-value'. We show examples of distributions of SI values in the population across the three phases. While in the SC phase, the S-haplotypes are entirely SC; in the SI phase, they are not entirely SI, as they used to be in the main model. The mixed phase shows bimodality with a 'pure' SC and partial SI sub-populations. The figures are based on simulation results of 4 independent runs for $\delta \in$ [0.75,1.0]. From each run, we extracted 5000 data points in 10-generation intervals. **(a-b)** The parameter values are as in Table 2 main text.

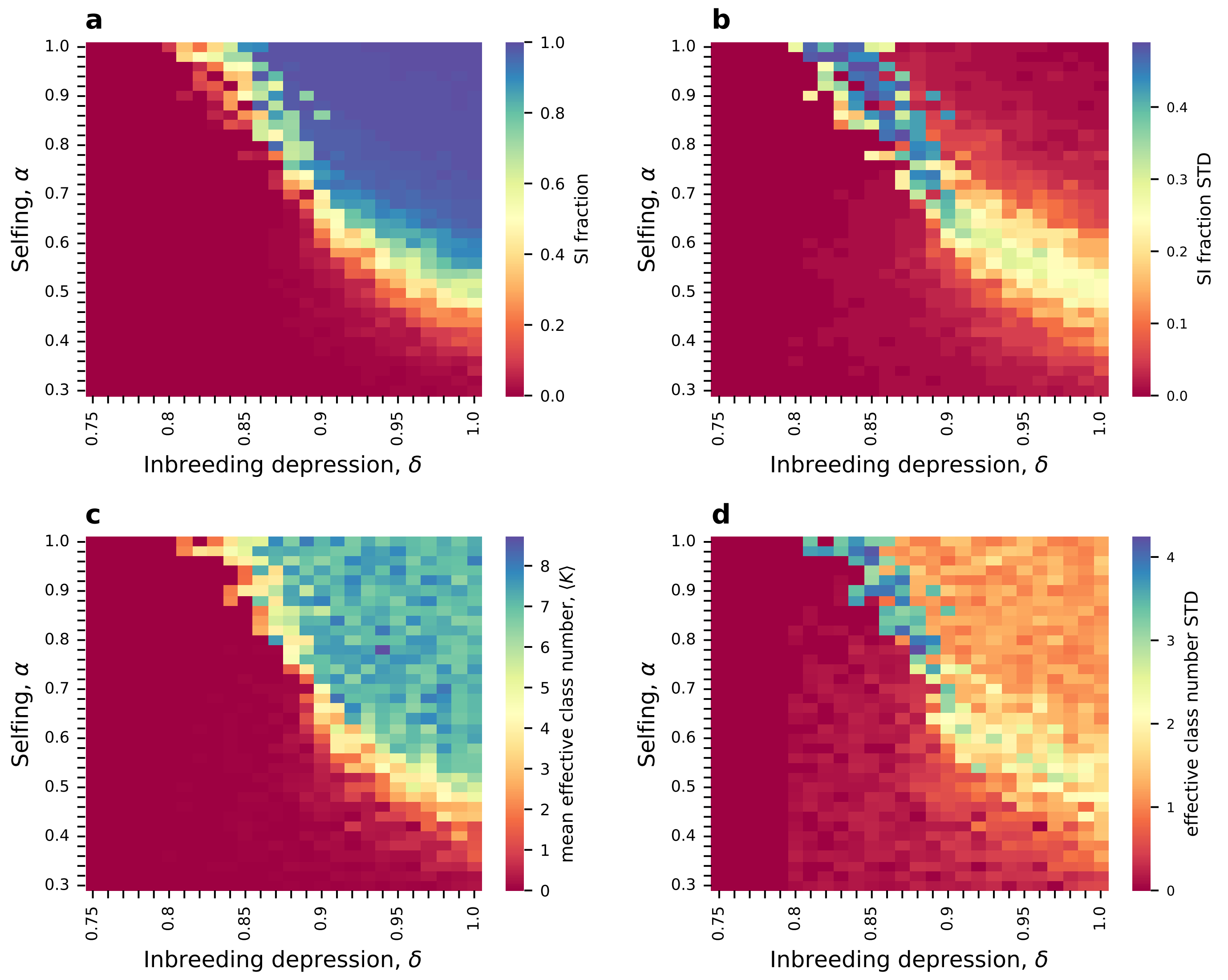

Figure S16: **The phase diagram of mating regimes projected on the ($\delta$,$\alpha$) parameter plane.** Unlike previous figures showing the phase diagram projected on the ($\delta$,$E_{\text{th}}$) plane with a fixed value of $\alpha$ (the proportion of self-pollen received), here we show this diagram in the ($\delta$,$\alpha$) plane with fixed $E_{\text{th}}$ = -6. **(a-b)** Maps of the self-incompatible population fraction's mean and standard deviation. **(c-d)** The mean and standard deviation of the effective number of classes. In this parameter plane we also observe the three mating regimes as before, including the unstable mixed phase, similar to the main model (compared to Fig. 4 in the main text). We find that the mating regime depends on both $\alpha$ and $\delta$: Self-incompatibility is obtained only if both $\delta$ and $\alpha$ are not too small, whereas self-compatibility is obtained if either $\delta < 0.8$ (regardless of $\alpha$) or if $\alpha < 0.3$ (regardless of $\delta$). The figures are based on simulation results of 4 independent runs. From each run, we extracted 2000 data points in 10-generation intervals. The minimal class size is 10 S-haplotypes. The remaining parameter values are as in Table 2, main text.

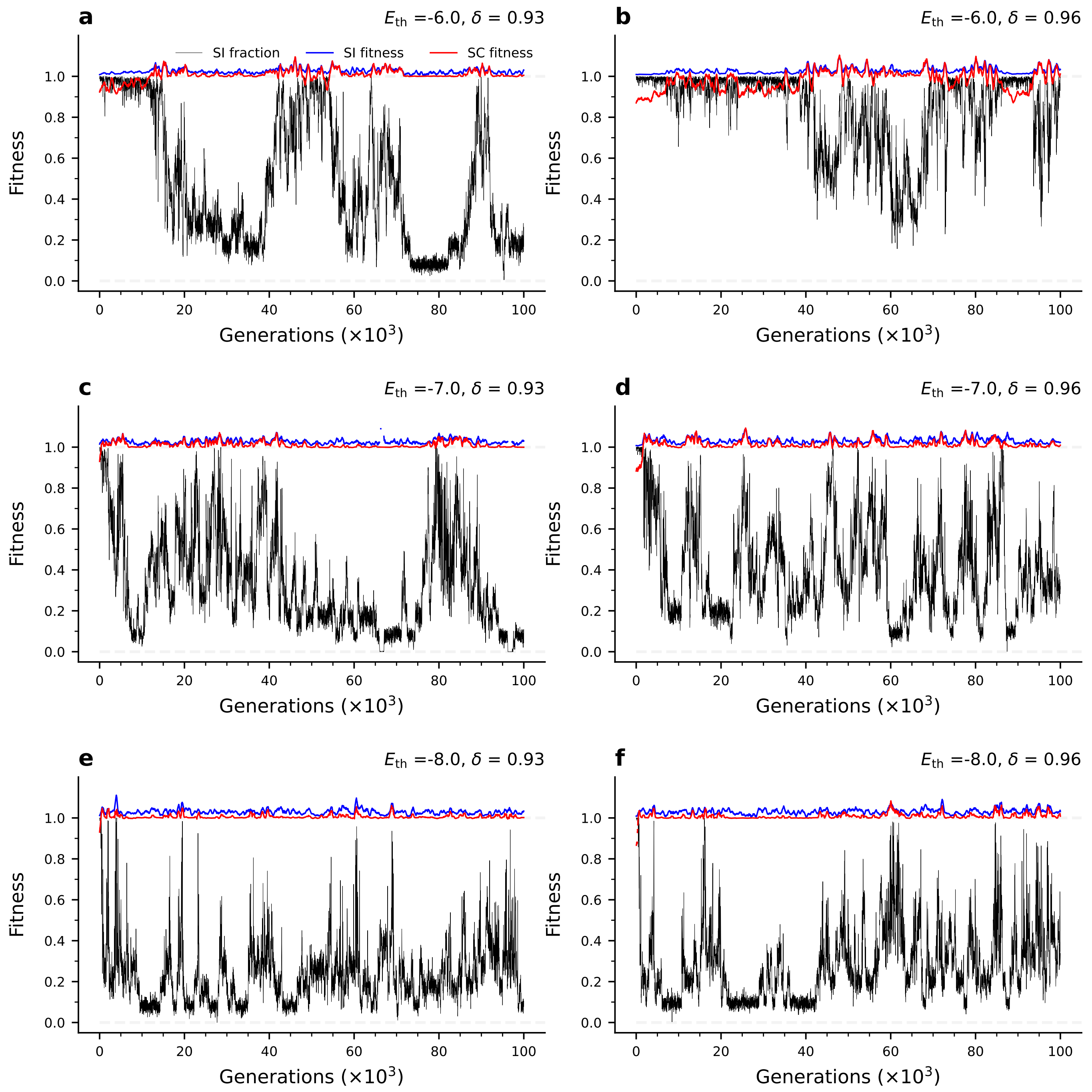

Figure S17: **The fitness values of SI and SC S-haplotypes in the mixed phase are stable and highly similar, despite large-amplitude temporal fluctuations in the SI population proportion.** Time series of SI (blue) and SC (red) S-haplotypes fitness values, and the SI population fraction (black) for different sets of parameters within the mixed phase. The fitness of the individual S-haplotype accounts for its reproductive contributions as both male and female. The formulas used to calculate fitness are provided in Table 3, main text. The remaining parameter values are as in Table 2, main text.

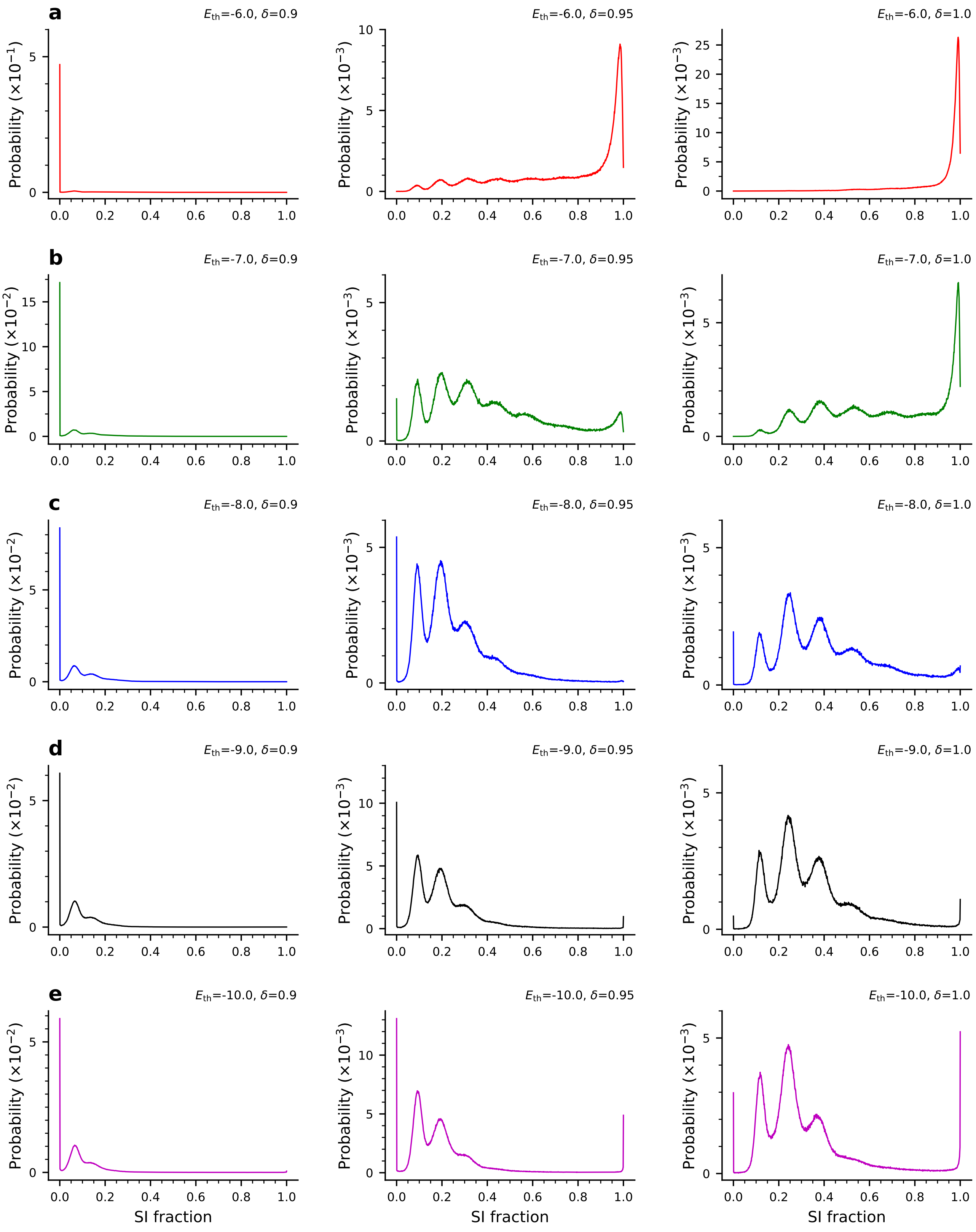

Figure S18: **Distributions of the population SI fraction under different parameter combinations in the phase plane.** This figure demonstrates a variety of population compositions: only SI as a single stable state (a right), only SC as a single stable state (a left), transitions between full SI and different SI-SC mixtures (a middle, b right), and similarly transitions between SC and different SI-SC mixtures (for example, c middle). There are also parameter combinations allowing all three behaviors: full SI, full SC, and different SI-SC mixtures (for example: b middle, e middle, and right). The figures are based on simulation results of 25 independent runs. From each run, we extracted 5000 data points in 10-generation intervals after discarding the initial 50,000 generations. The remaining parameter values are as in Table 2, main text.

### 4. Haploid model

#### 4.1 Probability of choosing a female S-haplotype as self-pollinated, self-compatible, out-crossed self-compatible, and self-incompatible

Consider $x$ and $y$ are the fraction of SC and SI S-haplotypes at a given generation G, where $x + y = 1$. Hence, the probability of choosing a female S-haplotype as self-pollinated self-compatible $f^{\rm s}_{\rm SC}$, out-crossed self-compatible $f^{\rm ns}_{\rm SC}$, and self-incompatible $f_{\rm SI}$ are as,

$$
\begin{aligned}
f^{\rm s}_{\rm SC} &= x\,\alpha\,(1-\delta) + \overbrace{\underbrace{(x\,\alpha\,\delta)}_{\text{dying fraction}}\ \underbrace{x\,\alpha\,(1-\delta)}_{\text{self-pollinated}}}^{\text{round 1}} + \overbrace{\underbrace{(x\,\alpha\,\delta)^2}_{\text{dying fraction}}\ \underbrace{x\,\alpha\,(1-\delta)}_{\text{self-pollinated}}}^{\text{round 2}} + \dots \\
f^{\rm ns}_{\rm SC} &= x\,(1-\alpha) + \overbrace{\underbrace{(x\,\alpha\,\delta)}_{\text{dying fraction}}\ \underbrace{x\,(1-\alpha)}_{\text{out-crossed}}}^{\text{round 1}} + \overbrace{\underbrace{(x\,\alpha\,\delta)^2}_{\text{dying fraction}}\ \underbrace{x\,(1-\alpha)}_{\text{out-crossed}}}^{\text{round 2}} + \dots \\
f_{\rm SI} &= y + \overbrace{\underbrace{(x\,\alpha\,\delta)}_{\text{dying fraction}}\ \underbrace{y}_{\text{self-incompatible}}}^{\text{round 1}} + \overbrace{\underbrace{(x\,\alpha\,\delta)^2}_{\text{dying fraction}}\ \underbrace{y}_{\text{self-incompatible}}}^{\text{round 2}} + \dots
\end{aligned}
$$

The first term in the right-hand side of the equations is the fraction of $f^{\rm s}_{\rm SC}$, $f^{\rm ns}_{\rm SC}$, and $f_{\rm SI}$. The second term represents the fraction of $f^{\rm s}_{\rm SC}$, $f^{\rm ns}_{\rm SC}$, and $f_{\rm SI}$ S-haplotypes being chosen after $x\ \alpha\ \delta$ fraction of SC S-haplotypes died - could not make up to adulthood due to inbreeding depression. The third term is the same as the second, but the fraction of S-haplotypes that could not make up to adulthood is $(x\ \alpha\ \delta)^2$. We have assumed that each maternal haplotype is definitely fertilized such that pollen is available in abundance. These equations lead to,

$$
\begin{aligned}
f^{\rm s}_{\rm SC} &= x\,\alpha\,(1-\delta)\,(1 + (x\,\alpha\,\delta) + (x\,\alpha\,\delta)^2 + \dots) && (11)\\
f^{\rm ns}_{\rm SC} &= x\,(1-\alpha)\,(1 + (x\,\alpha\,\delta) + (x\,\alpha\,\delta)^2 + \dots) && (12)\\
f_{\rm SI} &= y\,(1 + (x\,\alpha\,\delta) + (x\,\alpha\,\delta)^2 + \dots) && (13)
\end{aligned}
$$

The probability of choosing a female S-haplotype as $f^{\rm s}_{\rm SC}$, $f^{\rm ns}_{\rm SC}$, and $f_{\rm SI}$ are given as,

$$
\begin{aligned}
f^{\rm s}_{\rm SC} &= \frac{x\,\alpha\,(1-\delta)}{1 - x\,\alpha\,\delta} && (14)\\
f^{\rm ns}_{\rm SC} &= \frac{x\,(1-\alpha)}{1 - x\,\alpha\,\delta} && (15)\\
f_{\rm SI} &= \frac{1-x}{1 - x\,\alpha\,\delta} && (16)
\end{aligned}
$$

#### 4.2 Analytical calculation of boundaries between SC - mixed phase, and mixed - SI phase.

To provide an analytic interpretation of the boundaries between these distinct population states, we examine the reproductive capacity of each as the mean number of per-capita offspring that carry the same genotype as their parent. Recall that in our haploid model, the offspring inherits the genotype of one of its two parents with equal probability. In all cases, we assume sufficient pollen abundance, such that each S-haplotype is guaranteed to produce offspring as a maternal parent. The fitness of a self-compatible S-haplotype occupying a population proportion $0 \le A \le 1$ in the presence of $K$ equally-sized compatibility classes, each occupying a population proportion $x$, is:

$$
f_{\rm SC-K} = \underbrace{\frac{Kx}{2(1-x)}}_{\text{male}} + \underbrace{(1-\alpha)(A + \frac{1-A}{2})}_{\text{female non-self}} + \underbrace{\alpha(1-\delta)}_{\text{female self}}\ , \tag{17}
$$

where the first term stands for offspring it produces as male by fertilizing an SI maternal S-haplotype. The second and third terms stand for offspring it produces as a female by either outcrossing (with probability

$1-\alpha$) or by selfing (with probability $\alpha$), where the latter offspring survive with probability $1-\delta$. Outcrossing (2nd term) can be by either non-self SC or by SI pollen, whereas if both parents are SC, the offspring is also SC with probability 1. The factor 1/2 for outcrossing is needed because the offspring receives either parental genotype with equal probabilities. Similarly, the fitness of an SI individual from one of the $K$ classes reads:

$$f_{\mathrm{SI-K}} = \underbrace{\frac{1}{2}\left(A(1-\alpha) + \frac{(K-1)x}{1-x}\right)}_{\text{male}} + \underbrace{\frac{1}{2}}_{\text{female non-self}} . \tag{18}$$

The first term refers to offspring it produces as male by oucrossing either an SC or SI maternal S-haplotype of any of the other $K-1$ classes. The second term stands for offspring it produces as a maternal parent, in which case it is guaranteed to be fertilized owing to pollen abundance. Substituting $x = \frac{1-A}{K}$, and equating $f_{\mathrm{SC-K}} = f_{\mathrm{SI-K}}$, the boundary between the mixed and SI regimes is obtained at:

$$K^{**}(\delta) = 1 - A + \frac{1-A}{2\alpha(\delta - \frac{1}{2})}, \qquad \delta > \frac{1}{2}, \tag{19}$$

or alternatively at

$$\delta^{**}(K) = \frac{1}{2} + \frac{1-A}{2\alpha(K-1+A)} > \frac{1}{2}. \tag{20}$$

For $A = 0.1$ and $\alpha = 0.6$ (as in Fig. S21) we obtain the boundary at $K(\delta) = 0.9 + 0.75/(\delta - 0.5)$. As (19) shows, for a larger number of compatibility classes $K$, a lower inbreeding depression $\delta$ is sufficient to switch to the self-incompatibility mating mode, and vice versa. The reason is that as $K$ grows, self-incompatibility becomes more advantageous, because the population proportion with which a self-incompatible individual cannot mate (its own class) shrinks.

The transition between the SC and mixed mode occurs when SI mutants establish in the SC population, forming the first SI class. Thus, to obtain the second boundary between the SC and mixed mating modes, we substitute $K = 1$ in Eqs. (17)-(18) and obtain that this boundary is at a fixed $\delta$ value:

$$\delta^{*} = \frac{1}{2} + \frac{1-A}{2A\alpha} > \frac{1}{2}. \tag{21}$$

This boundary depends solely on $\delta$, in contrast to the boundary of (21) which depends on both $\delta$ and $K$. In Fig. S21 we used $A = 0.9$ and $\alpha = 0.6$ yielding $\delta^{*} = 0.593$. Since we use an arbitrary SC population proportion $A$ to calculate the boundary, this proportion could occasionally support more than one SI class. The boundary must then be calculated by substituting the exact number of classes. For example, if $K = 2$, we obtain a lower value of $\delta^{*} = 0.54$, because the more SI classes exist, the more advantageous the SI state becomes, hence a lower inbreeding depression suffices to maintain self-incompatibility. To fit the boundary in Fig. S21a we substituted the average number of classes obtained in simulations for the SC population proportion $A$. The two boundaries, calculated using Eqs. (20)-(21) are marked in Fig. S21a using gray and red dots, showing excellent agreement with the simulation results.

#### 4.3 The stochastic simulation

##### Initialization of the S-haplotype population

We initialize the S-haplotype population using $n$ distinct types, such that each type is self-incompatible and bidirectionally compatible with the remaining $n-1$ types. We initialize the population with equal frequencies of all types. We employ the following three steps to draw these initial types:

1. **Draw RNases sequences**: First, generate $n$ distinct RNases $\mathrm{R} = \{\mathrm{R}_1 \dots \mathrm{R}_n\}$ by randomly drawing sequences from the prior distribution $\nu$.

2. **Draw SLFs sequences**: Construct $n$ pools where each pool $\mathrm{P}_i = \{\mathrm{S}_m\}$, $i \in (1 \dots n)$, contains $m$ SLF sequences, that can detoxify the $i$-th RNase, and potentially others as well. To construct the SLF pools, we generate SLFs, one at a time, check which RNase(s) it detoxifies, and add that SLF into the corresponding pool(s). Any SLF that does not detoxify any RNase or that detoxifies all of them is discarded. Repeat this step until each pool has $m$ number of SLF genes.

3. **Form $N$ self-incompatible and complete S-haplotypes**: For each RNase $R_i \in R$, choose in total $n-1$ SLF genes, one from each pool $P_j$, $j \in (1 \ldots n)$, $j \neq i$ and make sure none of these SLFs detoxifies $R_i$. If it does, replace it with another SLF from the same pool until an SLF that does not detoxify $R_i$ is found. After generating $n$ distinct S-haplotypes, for an initial population of size, $N$ using $N/n$ copies of each S-haplotype, and shuffle their order.

The above procedure produces SLF that can potentially match multiple RNases each. Alternatively, to construct an initial population with only one-to-one RNase-SLF interactions, each pool $P_i$ should have only those SLFs that detoxify only $i^{\text{th}}$ RNase.

#### Main model: mutation and reproduction

1. **Mutation**: Each amino acid in each gene can mutate with probability $\mu$ per generation. If a mutation occurs at an amino acid, it is replaced by a randomly chosen amino acid drawn from the prior distribution $\nu$.

2. **Maternal S-haplotype picking**: Maternal SC S-haplotypes can be fertilized by non-self paternal S-haplotype or by the self one. In the latter case, only a proportion $1-\delta$ of this self-offspring survives. Maternal S-haplotypes chosen to reproduce can be either SC (with either self or non-self fertilization) or SI (only non-self fertilization). We compute the frequencies of either group forming offspring: $f^{\text{s}}_{\text{SC}}$, $f^{\text{ns}}_{\text{SC}}$, and $f_{\text{SI}}$, respectively:

$$f^{\text{s}}_{\text{SC}} = \frac{x\ \alpha\ (1-\delta)}{(1 - x\ \alpha\ \delta)} \tag{22}$$

$$f^{\text{ns}}_{\text{SC}} = \frac{x\ (1-\alpha)}{(1 - x\ \alpha\ \delta)} \tag{23}$$

$$f_{\text{SI}} = \frac{1-x}{(1 - x\ \alpha\ \delta)}, \tag{24}$$

where $x$, and $1-x$ are the population fraction of SC and SI S-haplotypes in the current generation, $\alpha$ is the proportion of self-pollen received and $\delta$ is the inbreeding depression, namely the proportion of self-fertilization offspring that do not survive. Full derivation of $f^{\text{s}}_{\text{SC}}$, $f^{\text{ns}}_{\text{SC}}$, and $f_{\text{SI}}$ is provided in the supplementary information file.

3. **Offspring formation**: Pick a maternal S-haplotype $H_i$ from either group with probabilities $\{f^{\text{s}}_{\text{SC}}, f^{\text{ns}}_{\text{SC}}, f_{\text{SI}}\}$. If the maternal S-haplotype $H_i$ belongs to $f^{\text{s}}_{\text{SC}}$, the offspring is identical to the parental one. If the maternal S-haplotype $H_i$ belongs to either $f^{\text{ns}}_{\text{SC}}$ or $f_{\text{SI}}$, it is fertilized by non-self pollen. In this case, it is given $k$ to match a randomly chosen non-self S-haplotype partner, in the paternal role. If such a match is found, the offspring would be one of the two parental S-haplotypes with equal probability. Repeat this step until $N$ offspring are formed.

Steps 1-3 are a single generation. Repeat these steps until the desired number of generations is reached.

#### Model-I: sequence mutation and RNase inactivation mutation

1. **Mutation**: Each amino acid in each gene (including RNase) can mutate with probability $\mu$ per generation. If a mutation occurs at an amino acid, it is replaced by a randomly chosen amino acid drawn from the prior distribution $\nu$. In addition to these sequence mutations, each RNase can alternate between active and inactive states at a rate $\mu_{\text{R}}$. In this model, an S-haplotype is considered self-compatible either if its RNase is inactive or if it is equipped with an SLF that can detoxify self-RNase.

2. **Maternal S-haplotype picking**: Same as in the section "Maternal S-haplotype picking".

3. **Offspring formation**: Pick a maternal S-haplotype $H_i$ from either group with probabilities $\{f^{\text{s}}_{\text{SC}}, f^{\text{ns}}_{\text{SC}}, f_{\text{SI}}\}$. If the maternal S-haplotype $H_i$ belongs to $f^{\text{s}}_{\text{SC}}$, the offspring is identical to the parental one. If the maternal S-haplotype $H_i$ belongs to either $f^{\text{ns}}_{\text{SC}}$ with an active RNase or $f_{\text{SI}}$, it is fertilized by non-self pollen. In this case, it is given $k$ attempts to match a randomly chosen non-self S-haplotype partner, in the paternal role. If such a match is found, the offspring would be one of the two parental S-haplotypes

with equal probability. If the maternal S-haplotype $H_i$ belongs to $f^{\text{ns}}_{\text{SC}}$ with an inactive RNase, any chosen non-self S-haplotype in the first attempt is the male partner, and the offspring would be any of these two.

Repeat this step until $N$ offspring are formed.

Steps 1-3 are a single generation. Repeat these steps until the desired number of generations is reached.

#### 4.4 Phase diagram

In Fig. 3, for each set of $E_{\text{th}}$ and $\delta$ values, we ran a total of four independent simulations, each ran for up to $10^5$ generations, of which we discarded the first $8.5 \times 10^4$ generations. In the part analyzed, we calculated the population fractions of SC and SI S-haplotypes every 25 generations. For every $E_{\text{th}}$ and $\delta$ combination, We lumped together all these ata points from all four independent runs and calculated the average fraction of SC and SI S-haplotypes. If the fraction of SI S-haplotypes $< 10\%$ (the threshold value used there) of the population, the system was considered to be in a self-compatible phase. If the SI fraction was $\geq 10\%$ and $< 90\%$, the system was considered to be in the mixed phase. Alternatively, if the SI fraction was $\geq 90\%$, it was considered to be in the self-incompatible phase. In the phase diagram, $E_{\text{th}}$ varied from -10 to -2 with a step size of 0.25, and $\delta$ varied from 0.0 to 0.99 with a step size of 0.03. The class numbers denoted on the right-side y-axis in Fig. 3a are the most probable class numbers calculated at $\delta = 0.90$. Phase diagrams with different thresholds defining the boundaries are also shown in supplementary Fig. S25, and for different $\alpha$ in supplementary Fig. S26.

#### 4.5 Fitting the simulation data to the fitness equations

Eqs. (17)-(18) state the fitness of SC and SI S-haplotypes, while the SI sub-population is comprised of $K$ equally-sized classes, and a proportion $A$ of the population is SC. To compare the boundary obtained in simulations, where classes may have different sizes (Fig. 3a), to this analytical result, we had to calculate the effective number of classes achieved with SC population proportions $A = 10\%$ and $A = 90\%$. To do that, we simulate the full model for $E_{\text{th}}$ from -10 to -4 at $\delta = 0.57$ – four independent runs for each $E_{\text{th}}$ value. We discarded the first $10^5$ generations and then calculated the effective number of classes for SI fractions between 7.5% and 12.5%, with up to 1000 time points per run. We averaged over the effective number of classes obtained in these multiple simulation instances for $E_{\text{th}}$ values between -10 to -4 and substituted these values as the class number $K$ in (21) to find the corresponding $\delta$ value. These points are shown as red circles in Fig. 3a.

Similarly, to fit the boundary between the mixed and SI phases, that was defined as the curve upon which $A = 90\%$ of the population, we calculated the effective number of classes at simulation data points for which the SI fraction was between 87.5% to 92.5% of the population and substituted the averge class number in the (20) to obtain the corresponding $\delta$ value. These points are shown as grey circles in Fig. 3a. For $E_{\text{th}}$ = -3 and -2, we found a direct transition between the SC and SI phases, with no mixed phase in between, such that the two boundaries overlap.

#### 4.6 Supplementary figures

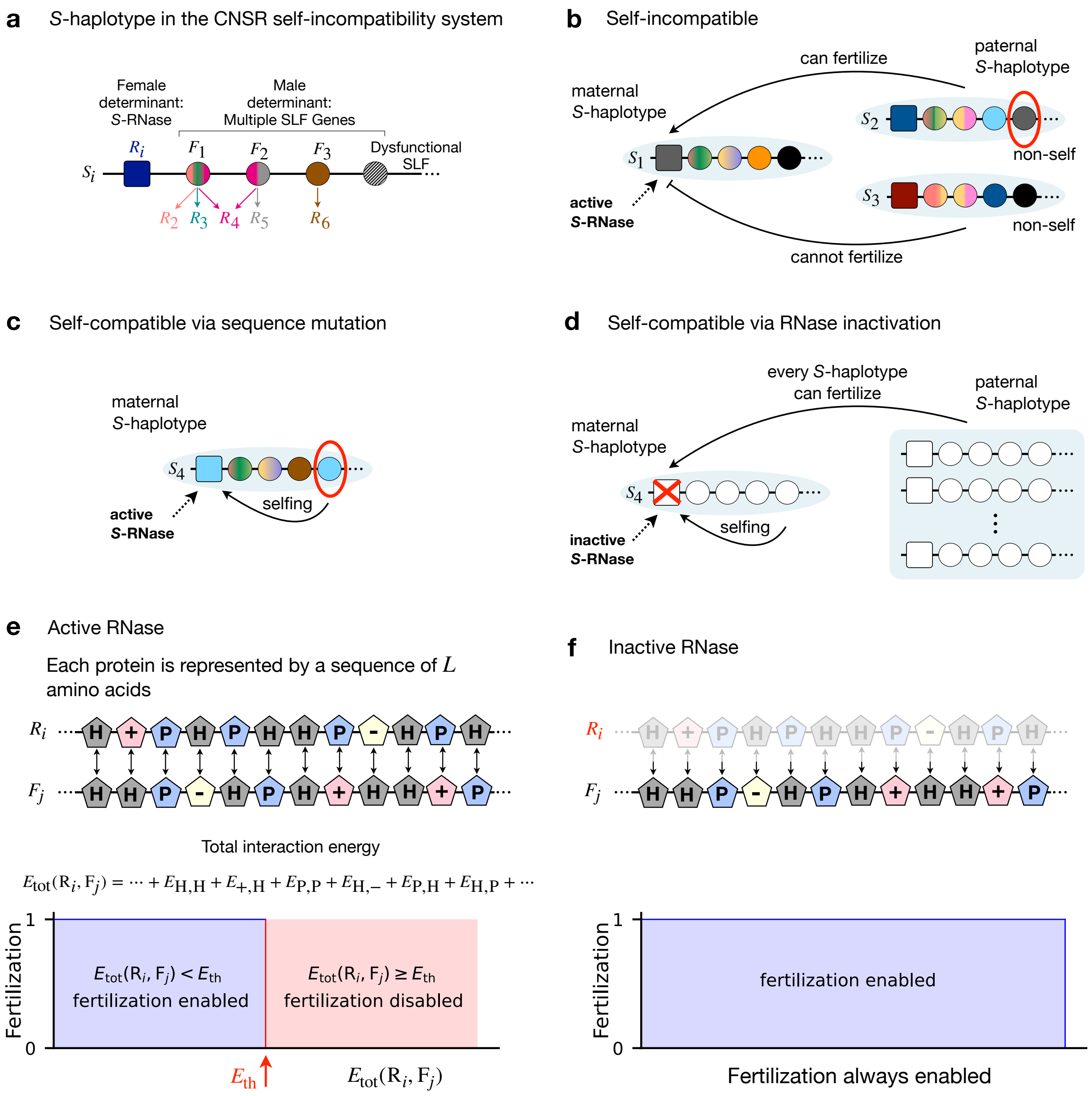

Figure S19: **Model description**. **(a)** Each S-haplotype consists of a single female determinant gene (RNase $R_i$, square) and multiple male-determinant genes (SLFs $F_1, F_2, \ldots$, circles). Each SLF could match one or more different RNases, or none. **(b)** For a paternal S-haplotype to successfully fertilize a maternal S-haplotype, it must be equipped with an SLF matching the maternal RNase. A maternal S-haplotype can be fertilized by either self (genetically identical) or non-self pollen. An S-haplotype is considered 'self-incompatible' if it does not contain an SLF matching its own RNase, hence fertilization by self-pollen is impossible. **(c-d)** An S-haplotype is considered 'self-compatible' either if it does contain an SLF matching its RNase and facilitating fertilization by self-pollen **(c)** or if its RNase is inactive **(d)**, in which case any S-haplotype can fertilize it, including the self one. **(e)** Each protein encoded in the S-haplotype is represented by a sequence of $L$ amino acids of the four biochemical categories. The total interaction energy $E_{\text{tot}}$ between an active RNase $R_i$ and an SLF $F_j$ is defined as the sum of the pairwise interaction energies between their corresponding amino acids. Two such proteins are considered matching if $E_{\text{tot}}$ is smaller than an energy threshold $E_{\text{th}}$, and otherwise they are non-matching. **(f)** If the RNase is inactive, the maternal S-haplotype carrying it can be fertilized by any SLF, regardless of its sequence.

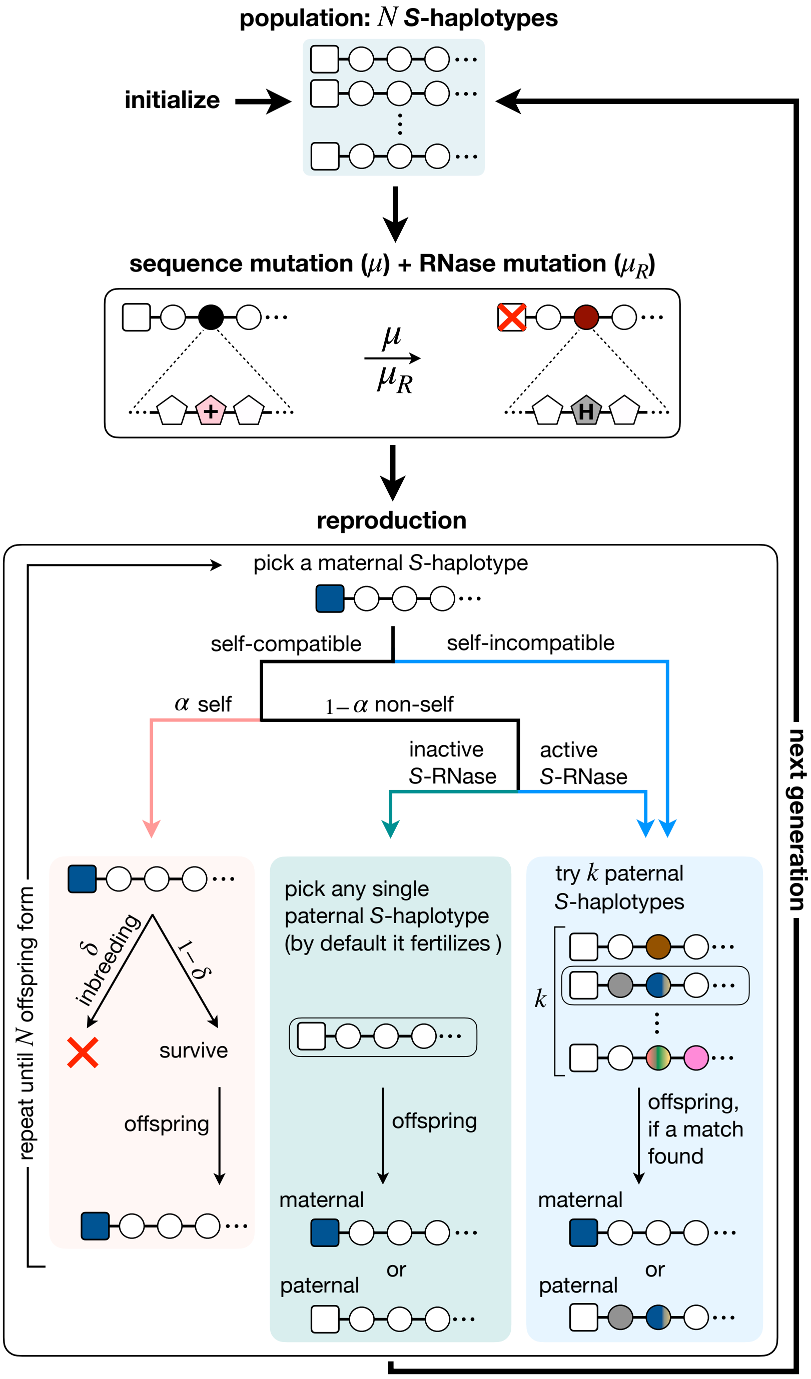

Figure S20: **The population life cycle implemented in a stochastic simulation.** The population consists of a fixed number $N$ of S-haplotypes, as shown in Fig. S19. Every generation, each of the amino acids in each of the proteins can mutate with probability $\mu$, and each RNase could alternate between active and inactive states with probability $\mu_R$. We assume non-overlapping generations. Every generation, maternal S-haplotypes are randomly selected for reproduction. If the maternal S-haplotype chosen is self-compatible, it can be fertilized by either non-self-pollen (with probability $1-\alpha$) or by self-pollen (with probability $\alpha$), where self-fertilization offspring survive with probability $1-\delta$ relative to offspring formed by non-self fertilization. Alternatively, if the maternal S-haplotype is self-incompatible, only non-self pollination is possible. Under non-self pollination, the maternal S-haplotype is given up to $k$ attempts to find a compatible paternal S-haplotype, by randomly picking S-haplotypes from the entire population. If the maternal S-haplotype is self-compatible due to an inactive RNase, the first paternal S-haplotype will fertilize. Following a successful fertilization, an offspring forms, inheriting either the paternal or the maternal S-haplotype, with equal probability. Maternal S-haplotype picking and offspring formation are repeated until $N$ offspring are formed. The offspring population then replaces the parental population, thereby completing one generation.

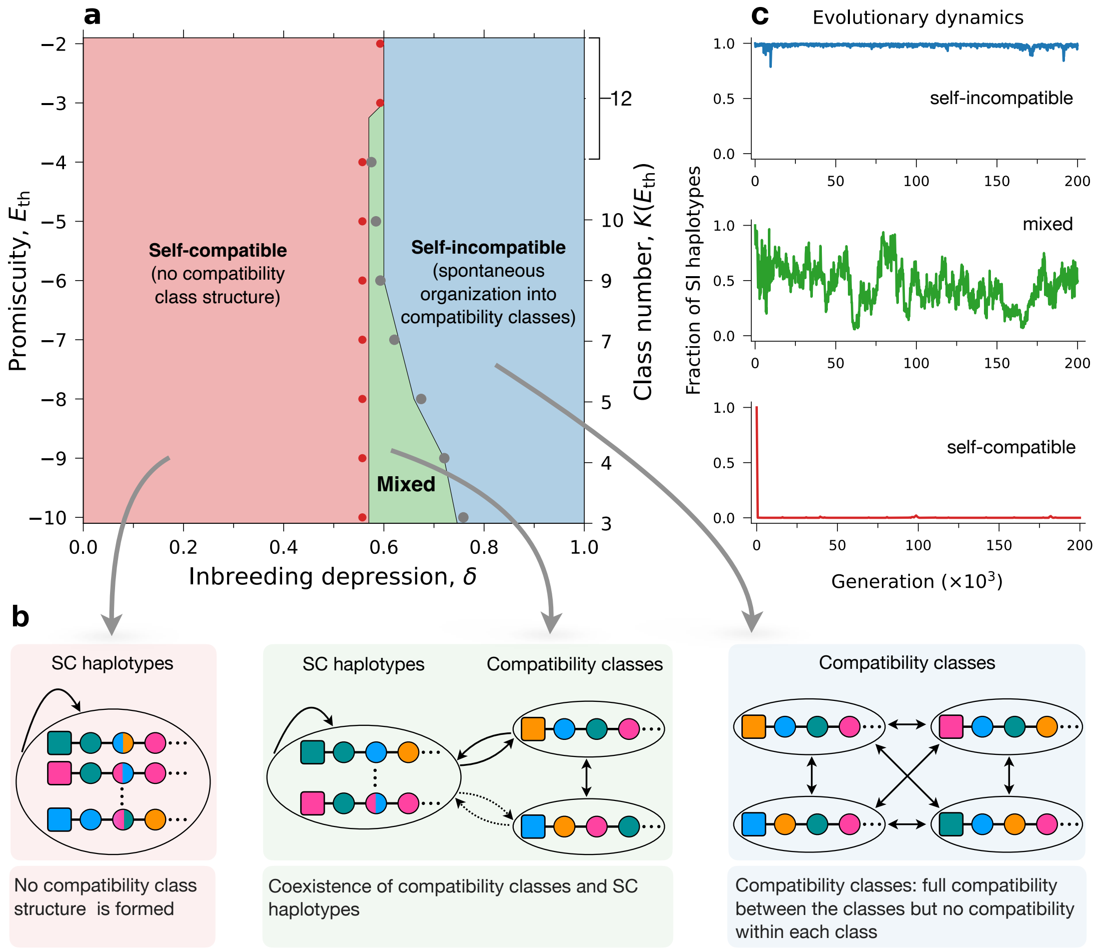

Figure S21: **The model exhibits three different phases: self-compatible, self-incompatible population partitioned into classes, or a mixture of self-compatible and incompatible sub-populations.** **(a)** The model phase diagram as a function of the inbreeding depression $\delta$ and the energy threshold $E_{\mathrm{th}}$ (promiscuity) parameters as obtained in simulations. The red and grey dots mark the boundaries between the phases, as predicted by Eqs. (20)-(21), using the arithmetic mean over the effective number of classes obtained in simulations under the relevant parameter values (see Methods). **(b)** Schematic illustration of the population structure in the three phases. Under high inbreeding depression, the population clusters into distinct compatibility classes, each of which is fully compatible with all the others. At the other extreme of low inbreeding depression, the population is fully self-compatible, namely, each individual can self-fertilize, and no class structure forms. In between, the population forms a mixture of classes fully compatible with each other and a self-compatible sub-population that could potentially be compatible with the classes. **(c)** Examples of temporal dynamics of the population self-incompatible fraction in each of the three phases. The self-compatible (top, blue) and self-incompatible (bottom, red) are dynamically stable, whereas the mixed phase (middle, green) exhibits vigorous fluctuations – see also Fig. S22d.

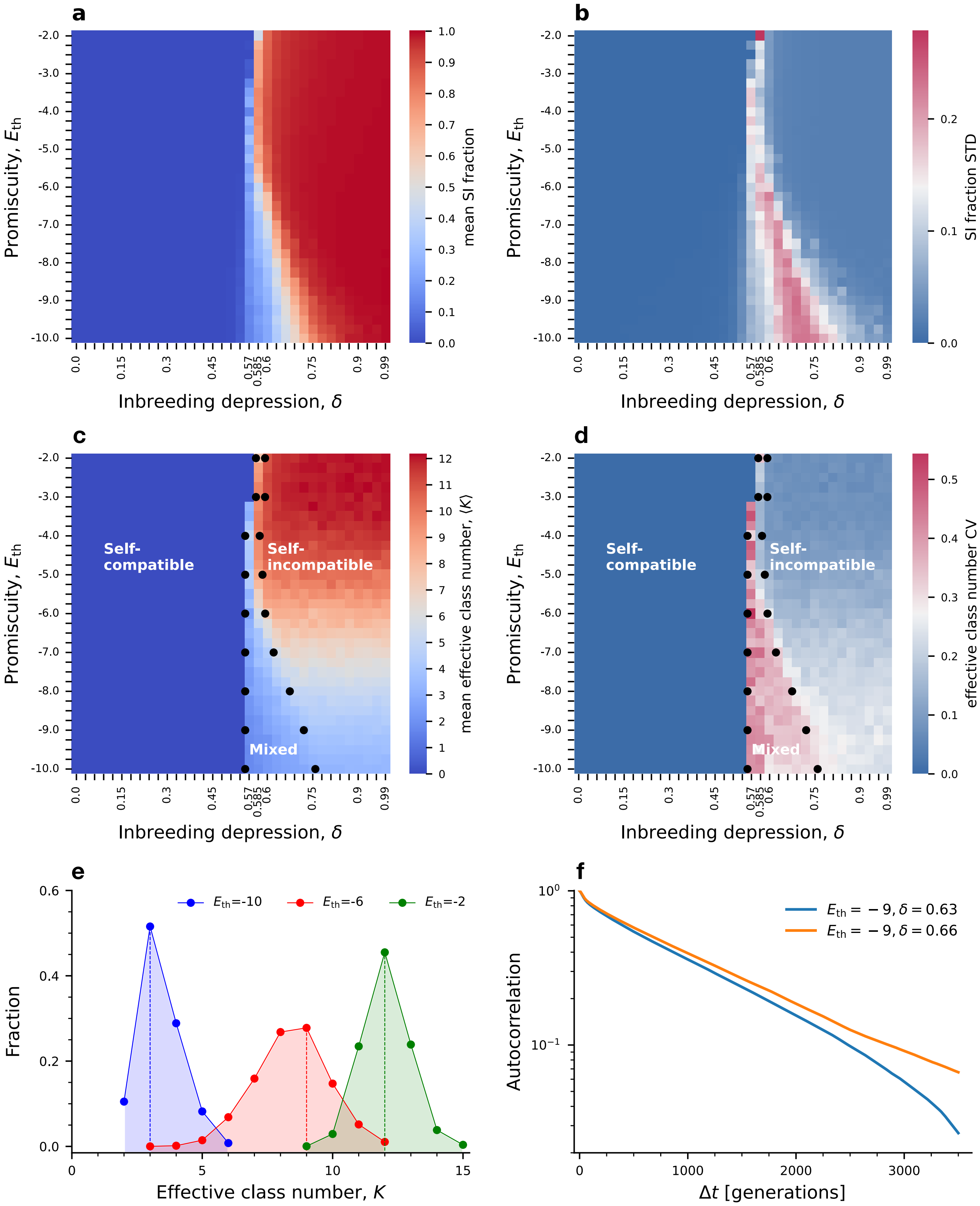

a
b
c
d
e
f
Promiscuity, $E_{th}$
Inbreeding depression, $\delta$
mean SI fraction
SI fraction STD
mean effective class number, $\langle K \rangle$
effective class number CV
Self-compatible
Self-incompatible
Mixed
Fraction
Effective class number, $K$
$E_{th}$=-10
$E_{th}$=-6
$E_{th}$=-2
Autocorrelation
$\Delta t$ [generations]
$E_{th} = -9, \delta = 0.63$
$E_{th} = -9, \delta = 0.66$

Figure S22

Figure S22 *(previous page)*: **Self-incompatible population fraction and number of classes map when only sequence mutations can cause self-compatibility. (a-b)** Maps of the mean and the standard deviation of the self-incompatible population fraction. **(c)** A map of the mean effective class number (shown by color-code) obtained in simulations for different combinations of the inbreeding depression $\delta$ and promiscuity $E_{\text{th}}$. In the left half-plane $\delta < \delta^*$, we find zero classes, in accord with the full self-compatibility regime, as shown in Fig. S21. The right half occupies the mixed and self-incompatible phases observed in Fig. S21 and indeed shows a non-zero number of classes, where for a fixed $\delta$ value their number increases with $E_{\text{th}}$. The change in class number at the right boundary of the self-compatibility phase is moderate for low $E_{\text{th}}$ and becomes more and more steep, as $E_{\text{th}}$ gets higher. **(d)** Coefficient of variation (standard deviation divided by the mean) in the number of classes clearly highlights the mixed phase as having a higher variation than the full SI and full SC phases. Black dots in (a-b) were calculated using Eqs. (17)-(18). **(e)** Effective class number distributions for different $E_{\text{th}}$ values ($\delta = 0.90$). The dashed vertical lines mark the most probable class number for each $E_{\text{th}}$. **(f)** The mixed phase exhibits a short temporal correlation compared to the SI and SC phases. We plot the unbiased temporal correlation $\gamma(\Delta t) = \left\langle \frac{\frac{1}{T-\Delta t}\left(\sum_{t=1}^{T-\Delta t}(r(t)-\mu)\ (r(t+\Delta t)-\mu)\right)}{\frac{1}{T}\sum_{t=1}^{T}(r(t)-\mu)^2} \right\rangle_S$, where $\mu = \frac{1}{S\ T}\sum_{s=1}^{S}\sum_{t=1}^{T} r(t)$, $T$ is the total number of points in the time series used. Each autocorrelation curve is based on 100 runs. The correlation was calculated over 3500 data points in 1-generation intervals. Total run time is $2 \times 10^5$, and the initial one-fourth of the data points are discarded.

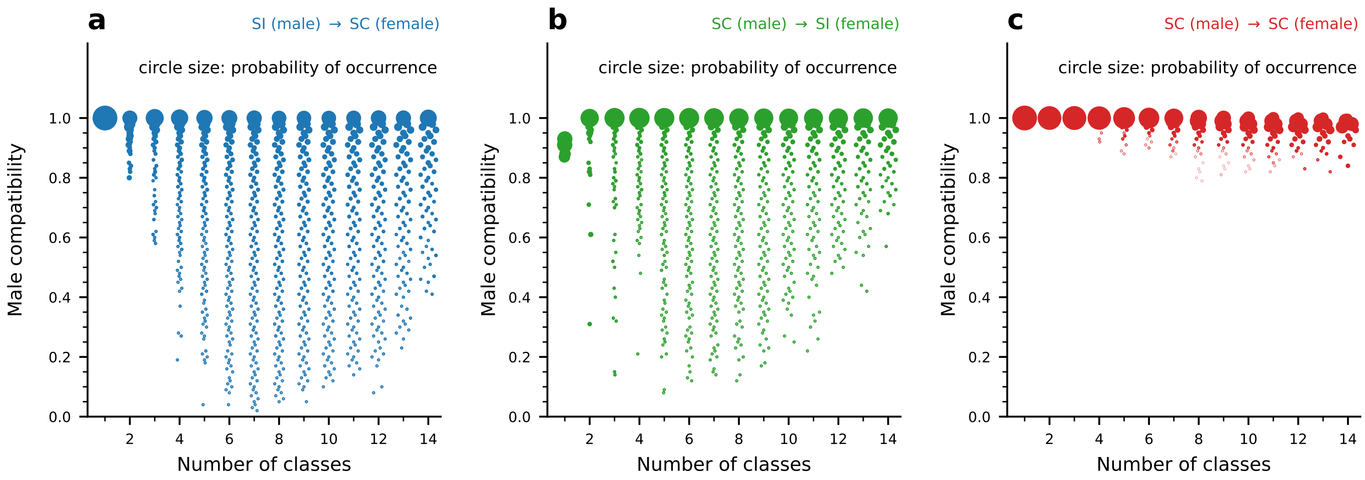

Figure S23: **Distributions of male-compatibilities between SI and SC sub-populations in the mixed phase vary with the number of SI classes.** Distributions of male-compatibilities, namely the population proportion an individual is compatible with as male, between SI (as a male) and SC (as a female) **(a)**, SC (as a male) and SI (as a female) **(b)** and SC in both male and female roles **(c)** in the mixed phase, dissected by the number of SI classes in the population (x-axis). Note that the number of classes is also affected by the varying proportion of the SI sub-population. The circle size corresponds to the probability of occurrence. Compatibility is highest amongst SC genotypes (c), but lower between SI and SC genotypes, both as males and as females (a-b). In the case of SC sub-population co-existing with only a single SI class, we find full SI-to-SC (a), but incomplete SC-to-SI male-compatibility (b). Once the number of classes increases, we observe a broadening of the distribution in both. An SI S-haplotype is incompatible with its class member; hence, if only a single class exists, it must be compatible with the SC sub-population to survive. This requirement is relieved once additional SI classes emerge. The figure is based on 4 independent runs, each with 10000 data points, and a 25-generation interval between consecutive time points used for the analysis. Total run time is $2 \times 10^5$. Parameters: $E_{\mathrm{th}}$ = -6, $\alpha = 0.60$, and $\delta = 0.6$, and the values of the remaining parameters are as in Table 2 main text except $N = 2000$.

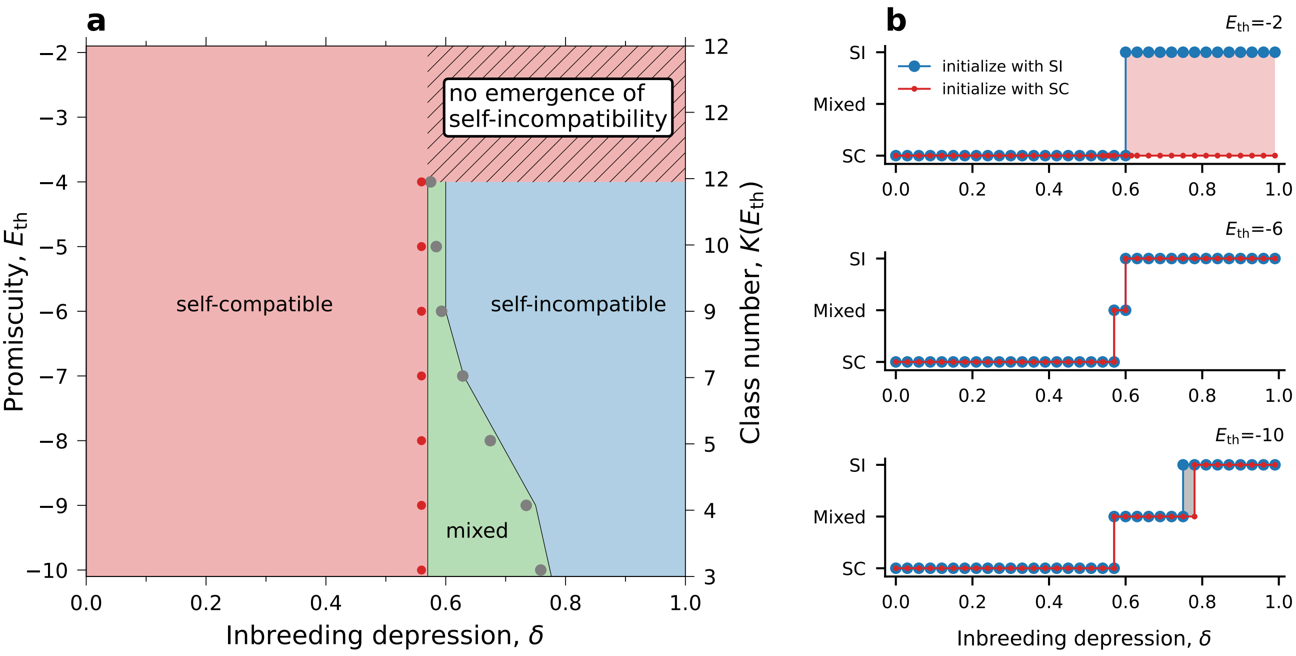

Figure S24: **In most of the parameter range, except for the highest $E_{\text{th}}$, the phase diagram is independent of the population's initial condition: whether it starts as fully self-compatible or fully self-incompatible**. **(a)** Phase diagram similar to Fig. S21 obtained when the population was initialized as fully self-compatible. Over most of the parameter range, we observe the same three phases with boundaries similar to those in Fig. S21, where the population was initiated as fully self-incompatible. Only for high promiscuity $E_{\text{th}} \geq -4$, self-incompatibility does not emerge (region marked by diagonal stripes). The calculation of an effective number of classes $K$ (right-side y-axis) was carried out at $\delta = 0.90$. To obtain this diagram, we ran simulations for different combinations of the parameters $E_{\text{th}}$ (with a step size of 1) and $\delta$ (with a step size of 0.03) – 4 independent runs for each combination. Each simulation was run for $2 \times 10^5$ generations, but we discarded the initial $1.5 \times 10^5$ and analyzed only the last 50,000 generations. We used 2000 data points with 25-generation intervals between consecutive time points from each run. The boundaries between the phases were set as the parameter combination for which, on average over these multiple data points, the population SI proportions equal the threshold values. **(b)** The population final state (SC, SI or mixed) as a function of the inbreeding depression $\delta$ for three different $E_{\text{th}}$ values, starting from either fully self-incompatible (red) or fully self-compatible (blue) population state. This graph is a cross-section of the phase diagrams of Fig. S21a starting from full SI, and the analogous one in (a) starting from full SC. Parameter values: $\alpha = 0.60$.

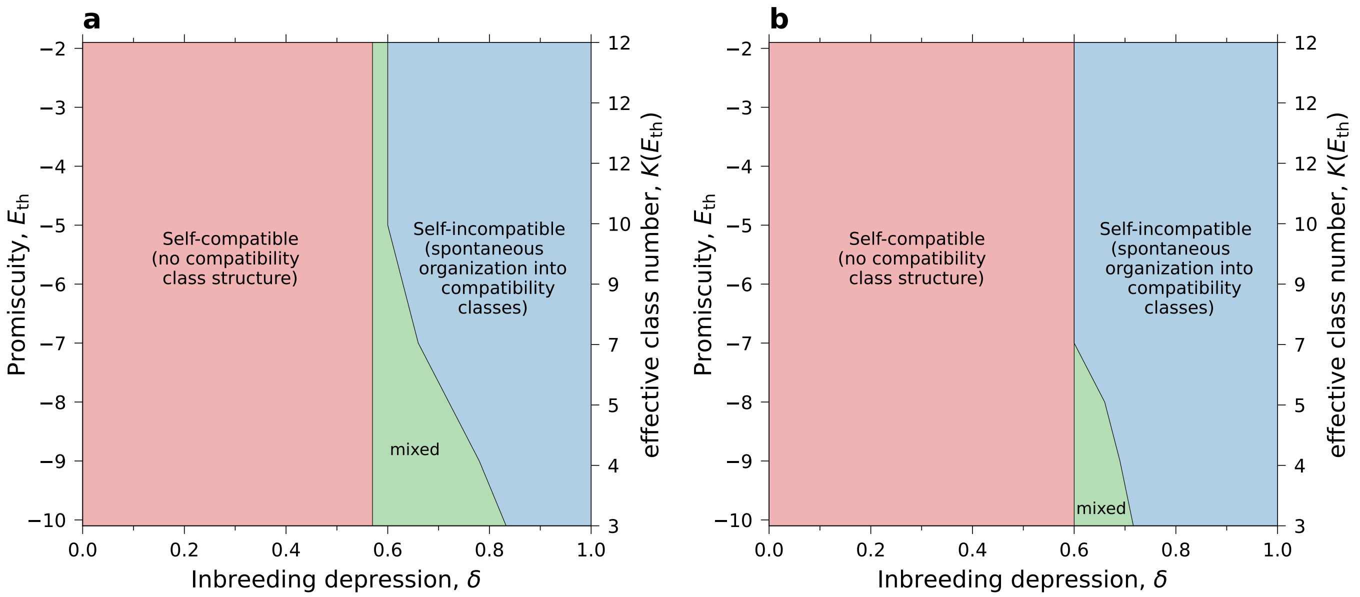

Figure S25: **The three different phases: self-compatible, self-incompatible, and mixed are obtained for various threshold values defining the boundaries between the phases**. The phase diagram, similar to Fig. 3a (main text) but with threshold values of 5% and 95% **(a)** or 20% and 80% **(b)** of the SI S-haplotype population proportions defining the mixed phase boundaries (compared to 10% and 90% in the main text). The plot is based on 4 independent runs for each set of ($E_{th}$, $\delta$) parameter values. We have taken 2000 data points from the end of each run in 25-generation intervals (total run time $2 \times 10^5$ generations). The boundaries between the phases were set as the parameter combination for which, on average over these multiple data points, the population SI proportions equal the threshold values. The interaction energy threshold $E_{th}$ varies from -10 to -2 with a step size of +1, and the inbreeding depression value varies from 0.0 to 0.99 with a step size of 0.03. The total run time for each parameter set is $2 \times 10^5$ generations. The calculation of an effective number of classes $K$ is carried out at $\delta = 0.90$. Parameter values: $\alpha = 0.60$. The remaining parameters are as in Table 2, main text except $N = 2000$.

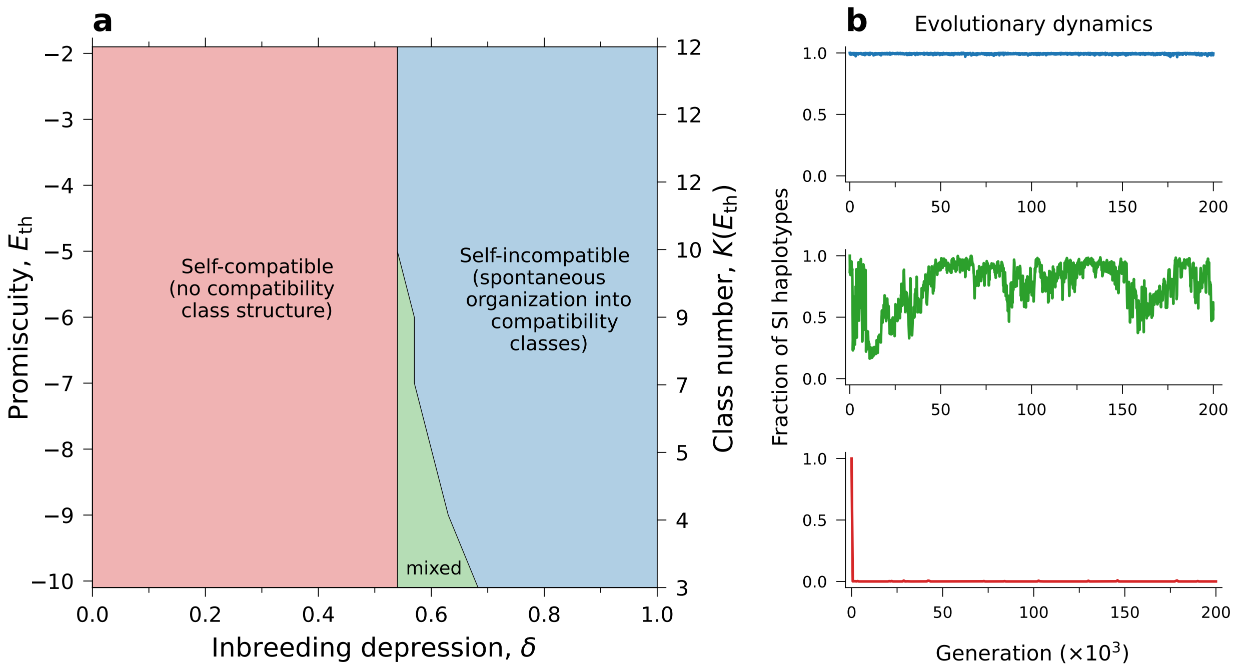

Figure S26: **The three different phases: self-compatible, self-incompatible, and mixed are also obtained for different values of the self-pollination rate** $\alpha$. Similar to Fig. 3 (main text), but with $\alpha = 0.90$. The plot is based on 10 independent runs for each set of ($E_{th}$, $\delta$) parameter values. We have taken 2000 data points from the end of each run in 25-generation intervals (total run time $2 \times 10^5$ generations). The interaction energy threshold $E_{th}$ varies from -10 to -2 with an increase of +1, and inbreeding depression varies from 0.0 to 0.99 with a step size of 0.03. The total run time for each parameter set is $2 \times 10^5$ generations. The boundaries between the phases were set as the parameter combination for which, on average over these multiple data points, the population SI proportions equal the threshold values. The calculation of an effective number of classes $K$ (right-side y-axis) is carried out at $\delta = 0.90$. We used the default parameter values listed in Table 2 main text except $N = 2000$.

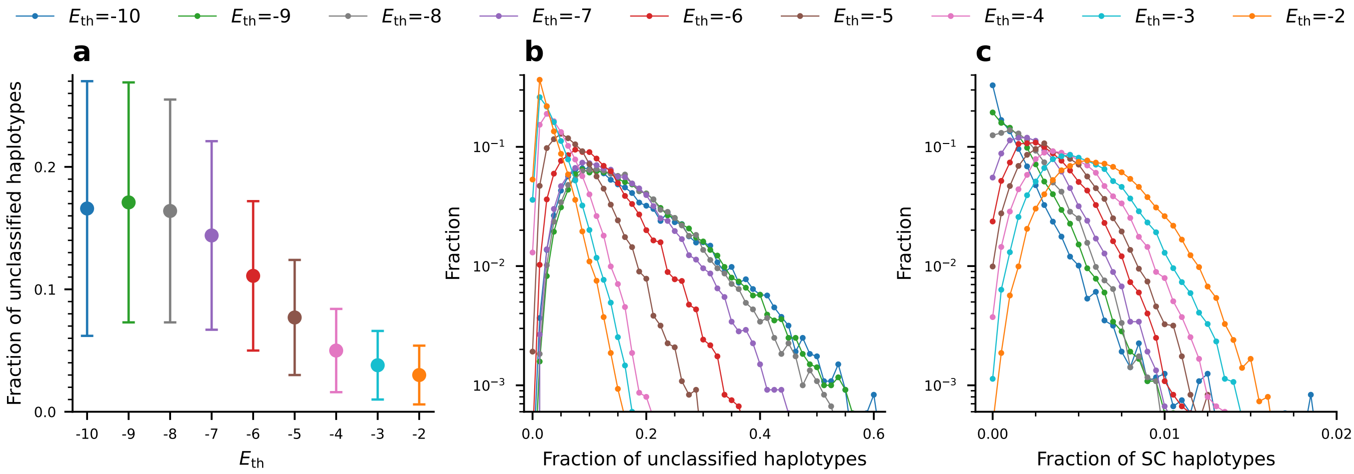

Figure S27: **The population proportion of unclassified S-haplotypes decreases, and the proportion of self-compatible ones increases with $E_{\mathrm{th}}$ in the SI phase**. **(a)** The average population proportion of unclassified S-haplotypes (either SI or SC) for different values of $E_{\mathrm{th}}$, denoted by different colors. The colored circle and bar represent the mean and the $\pm$ STD. **(b-c)** Distributions of the population proportions of unclassified S-haplotypes (either SI or SC) (b) and self-compatible S-haplotypes (c) for different values of $E_{\mathrm{th}}$. We observe that the proportion of unclassified S-haplotypes decreases, while the proportion of self-compatible S-haplotypes increases with $E_{\mathrm{th}}$. The figure is based on 16 independent runs for each $k$ value, with 1200 data points per run and a 25-generation interval between consecutive time points used for the analysis. All these data points were taken after the entire population descended from a single common ancestor, and the amino acid frequencies have already reached their steady-state values. The total runtime for each is $2 \times 10^5$ generations. Parameter values: interaction energy thresholds $E_{\mathrm{th}} = -6$. The remaining parameters are as in Table 2 main text except $N = 2000$.

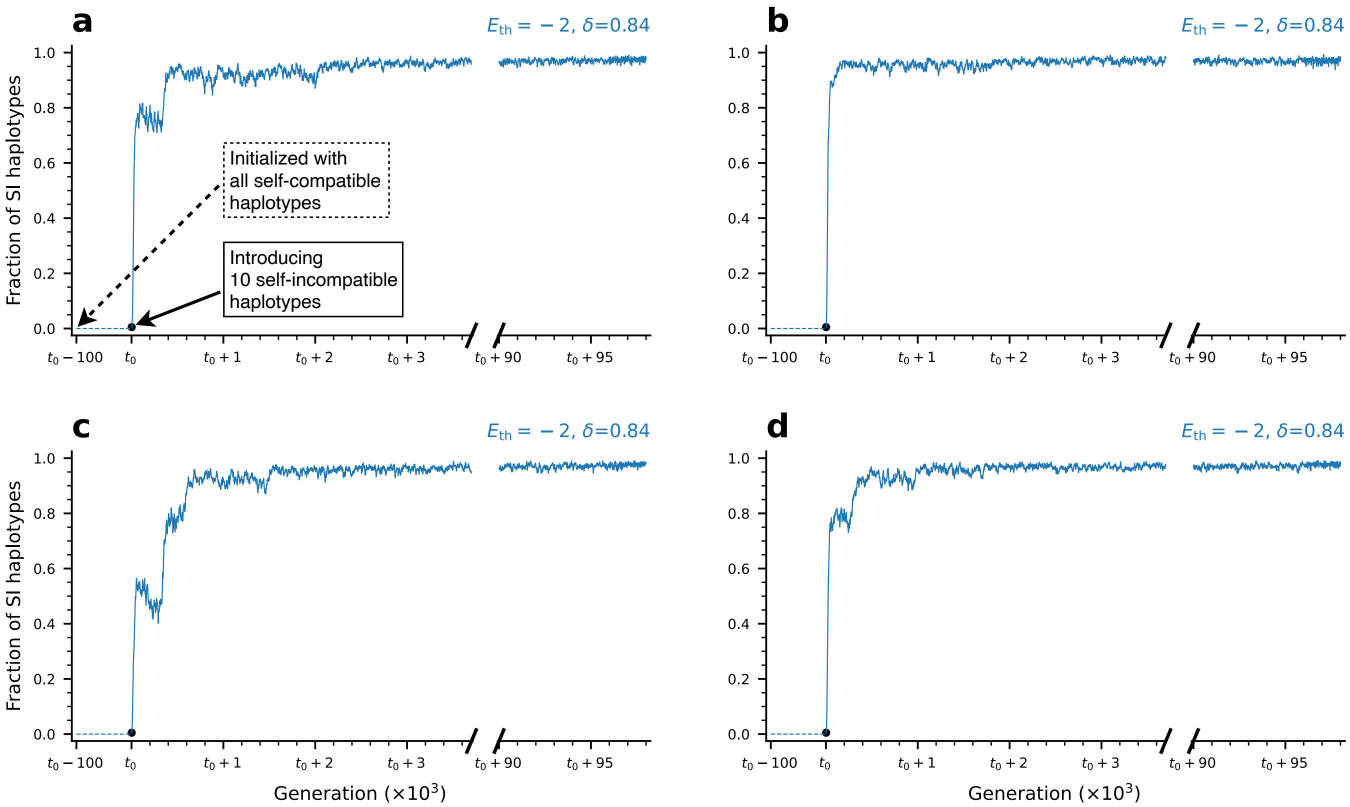

Figure S28: **Beginning from a fully self-compatible population, self-incompatibility can emerge under a high interaction energy threshold if SI S-haplotypes are manually introduced**. **(a-d)** The population proportion of SI S-haplotypes in 4 independent simulation runs under $E_{\text{th}}$ =-2 initialized with a fully self-compatible population. Initially, no emergence of self-incompatibility is observed (blue dashed line) after waiting for $10^5$ generations. SI quickly stabilizes within 4000 generations after introducing 10 self-incompatible S-haplotypes into the population at generation $t_0$. Once it emerged, the self-incompatibility population state persists - we have tested that for an additional 90,000 generations from $t_0$ as $t_0 + 90$. This demonstrates that the transition from self-compatibility to self-incompatibility under parameter values supporting the stable existence of self-incompatibility was not observed under high $E_{\text{th}}$ only because the probability of mutations converting self-compatible S-haplotypes into self-incompatible ones is extremely low under high $E_{\text{th}}$, hence the waiting time for such mutations is infeasibly long. (a-d) show different independent runs. The data from $t_0$ to $t_0 + 4$ ($\times 10^3$) generations are plotted in 2-generation intervals and from $t_0 + 90$ ($\times 10^3$) in 10-generation intervals. Parameter values: $E_{\text{th}} = -2, \delta = 0.84$. The remaining parameters are as in Table 2 main text, $N = 2000$.

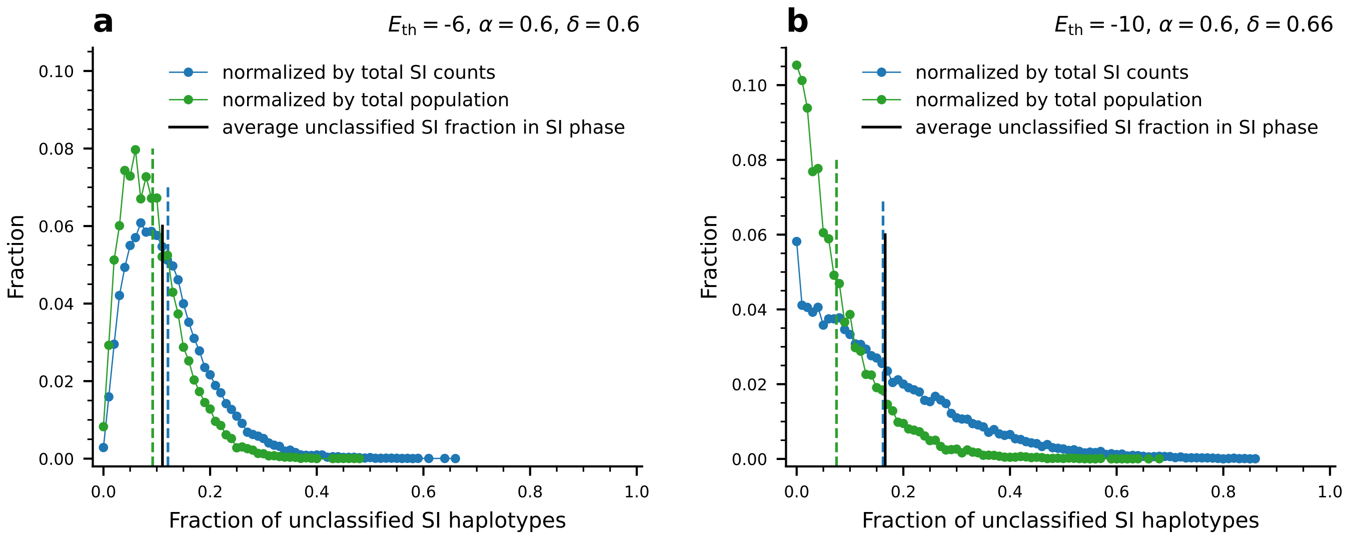

Figure S29: **Distribution of the population proportion of unclassified SI S-haplotypes in the mixed phase**. Distributions of the population proportion of unclassified self-incompatible S-haplotypes in the mixed phase for $E_{\text{th}}$ = -6, $\alpha$ = 0.60, and $\delta$ = 0.60 **(a)**, and for $E_{\text{th}}$ = -10, $\alpha$ = 0.60, and $\delta$ = 0.66 **(b)**. We show both the population proportion among the total self-incompatible (blue) and among the entire population (green). The vertical dashed lines mark the distribution mean values. Compare to the unclassified proportion in the SI phase shown in Fig. S27. The black vertical lines mark the mean fraction of unclassified S-haplotypes in the SI phase, given here for reference. We observe that despite the rapid changes in the self-incompatible population proportion, the unclassified S-haplotype proportion is very close to its value in the self-incompatible phase, in which the self-incompatible population proportion is stable. The figure is based on 4 independent runs, each with 10000 data points, and a 25-generation interval between consecutive time points used for the analysis. Total run time is $2 \times 10^5$. Parameters: the values of the remaining parameters are as in Table 2 main text, $N = 2000$.

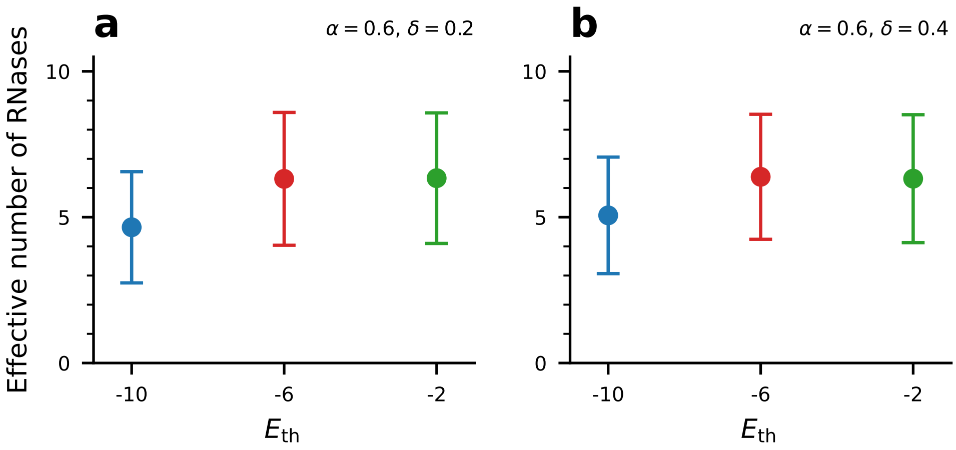

Figure S30: **There are multiple distinct genotypes in the SC phase.** The effective number of distinct RNases in the population under different $E_{\mathrm{th}}$ values for $\delta = 0.20$ **(a)**, and for $\delta = 0.40$ **(b)**. Circles denote the mean values and bars show ±STD relative to the mean. The figure is based on 4 independent runs, with the last 600 data points taken from each run, with a 25-generation interval between consecutive time points used for the analysis. All these data points were taken after the entire population descended from a single common ancestor, discarding the first 87,500 generations. The total runtime is $1 \times 10^5$ generations. Parameter values: $\alpha = 0.60$; the values of the remaining parameters are as in Table 2 main text, $N = 2000$.

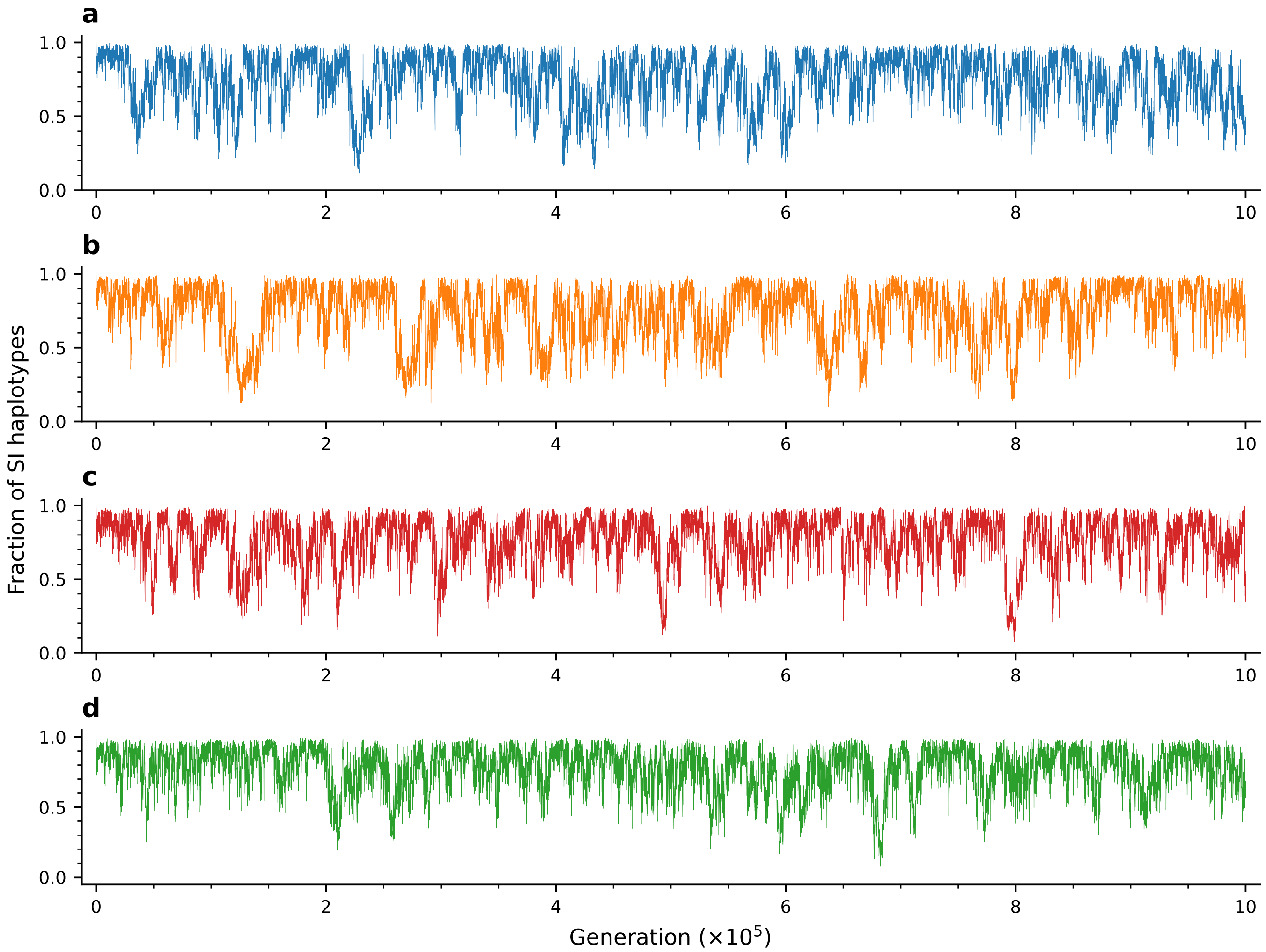

Figure S31: **The mixed phase persists for a long time starting from an initially self-incompatible population**. We show here time traces of 4 independent runs of the full model with the parameter combination associated with the mixed-phase ($E_{\mathrm{th}} = -6$, $\alpha = 0.6$ and $\delta = 0.6$), each run up to $10^6$ generations. All runs were initialized with a fully self-incompatible population. The self-incompatible population proportion is plotted here in 25 generation time intervals. Parameter values are as in Table 2 main text, $N = 2000$.

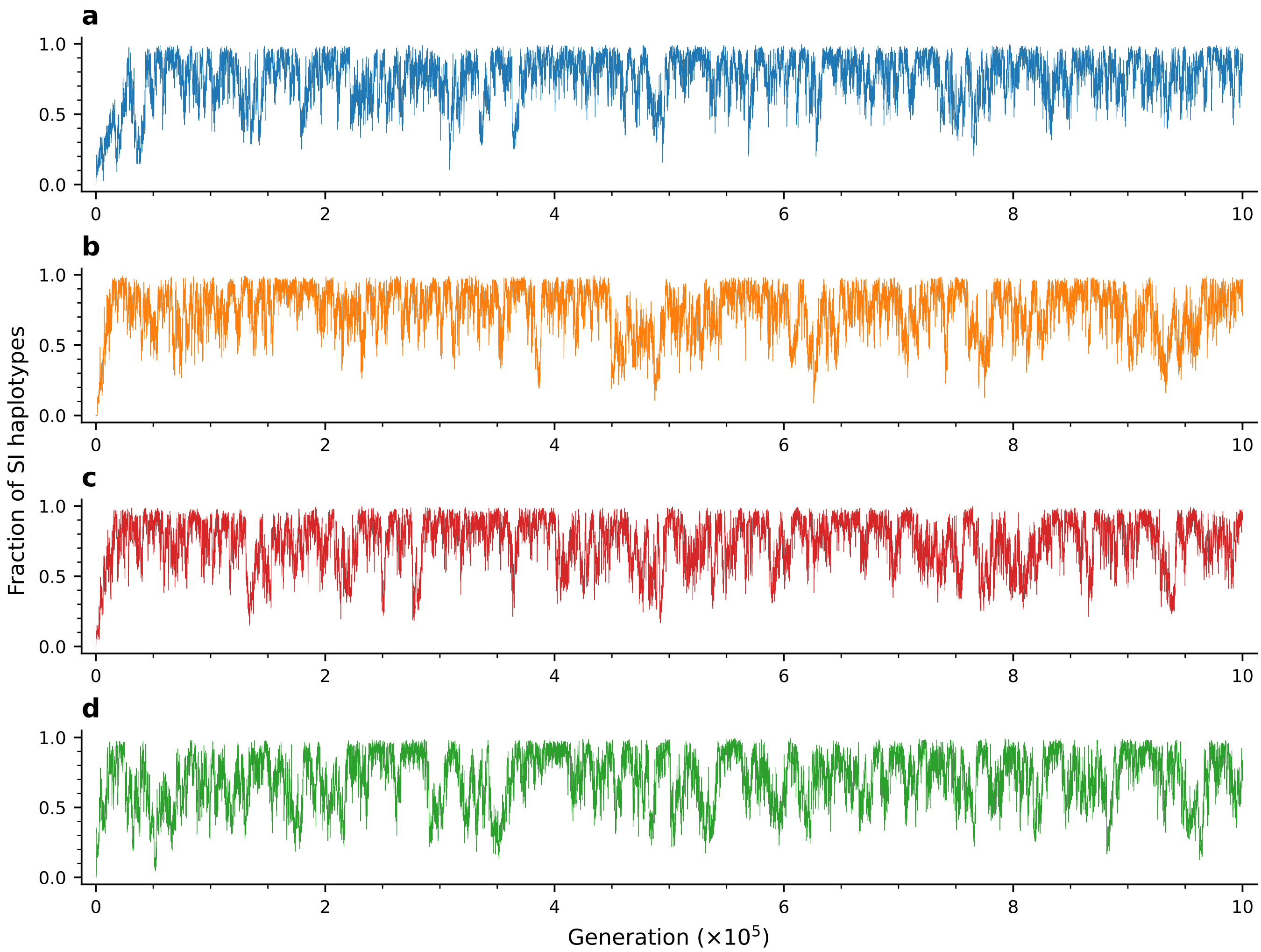

Figure S32: **The mixed phase persists for a long time starting from an initially self-compatible population**. We show here time traces of 4 independent runs of the full model with parameter combination associated with the mixed-phase ($E_{\rm th} = -6$, $\alpha = 0.6$ and $\delta = 0.6$), ran up to $10^6$ generations. All runs were initialized with a fully self-compatible population. The self-incompatible population proportion is plotted here in 25 generation time intervals. Parameter values are as in Table 2 main text, $N = 2000$.

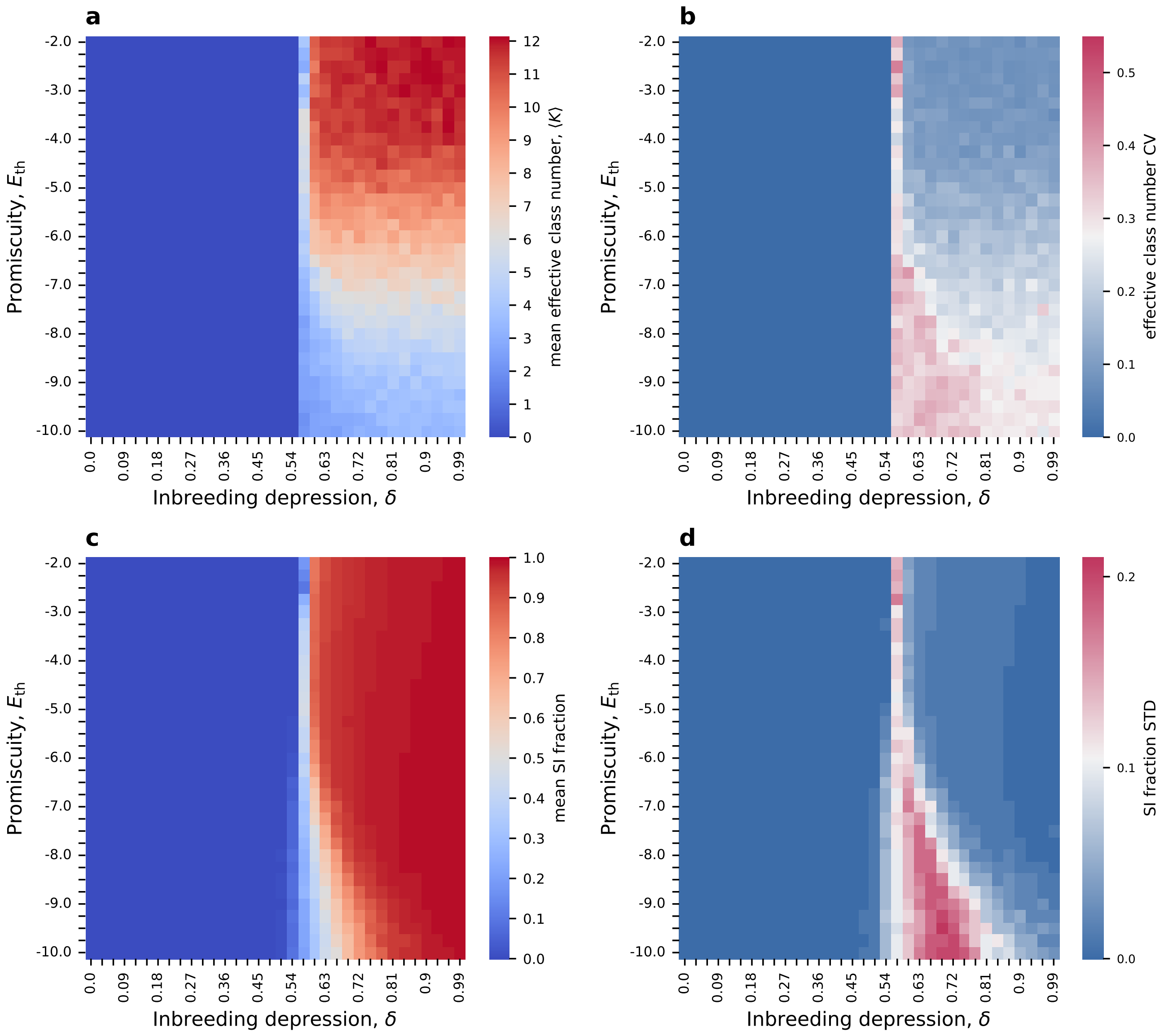

Figure S33: **Class number and self-incompatible population fraction maps when both sequence and RNase inactivation mutations can cause self-compatibility.** **(a-b)** The mean and the CV (std/mean) of the effective number of classes, for SI population fraction greater than 10% (here obtained for $\delta > 0.54$). **(c-d)** Maps of the mean and the standard deviation of the self-incompatible population fraction. Here we observe the unstable mixed phase, similar to the case that only sequence mutations could cause self-compatibility (compare to Fig. S22). Only the boundaries between the different phases are slightly shifted. $\mu_R = 10^{-4} \cdot L$. Other parameter values are as in Table 2 main text, $N = 2000$.

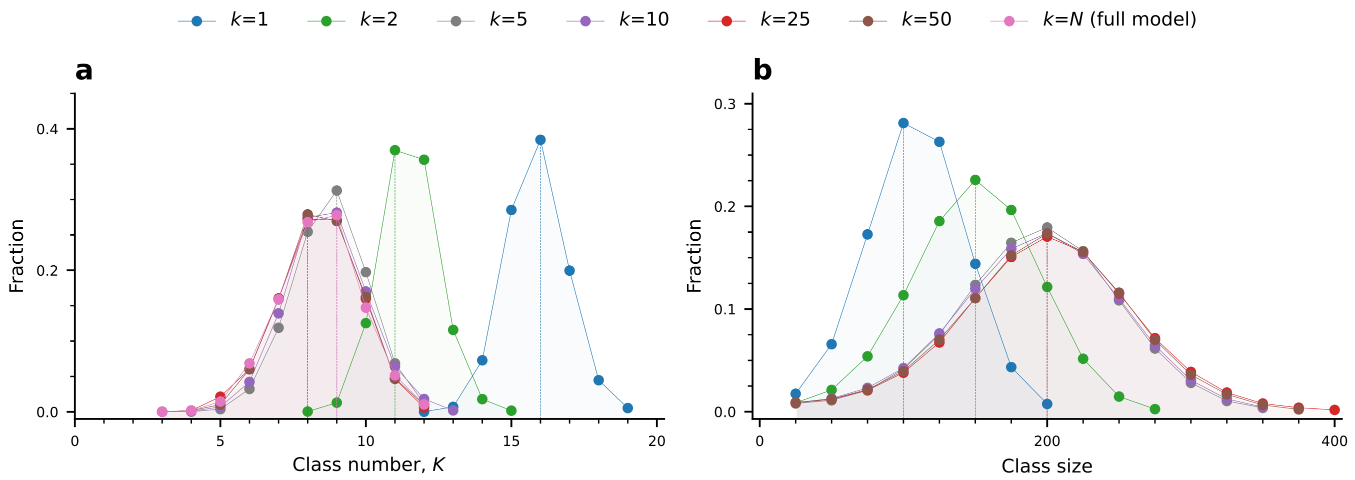

Figure S34: **The class number increases and the class size decreases under pollen limitation. (a-b)** Distributions of the number of classes in the population (a) and the class size (b) for different numbers of fertilization attempts $k$ per maternal S-haplotype. Distributions associated with different values of $k$ are colored differently. We find an increase in the class number and, accordingly, a decrease in the class size under extreme pollen limitation ($k = 1, 2$), but for $k \geq 5$ the distributions overlap, showing insensitivity to further increases in pollen abundance. The figure is based on 16 independent runs for each $k$ value, with 1200 data points per run and a 25-generation interval between consecutive time points used for the analysis. All these data points were taken after the entire population descended from a single common ancestor, and the amino acid frequencies have already reached their steady-state values. The total runtime for each is $2 \times 10^5$ generations. Parameter values: interaction energy thresholds $E_{\mathrm{th}} = -6$. The remaining parameters are as in Table 2 main text, $N = 2000$.

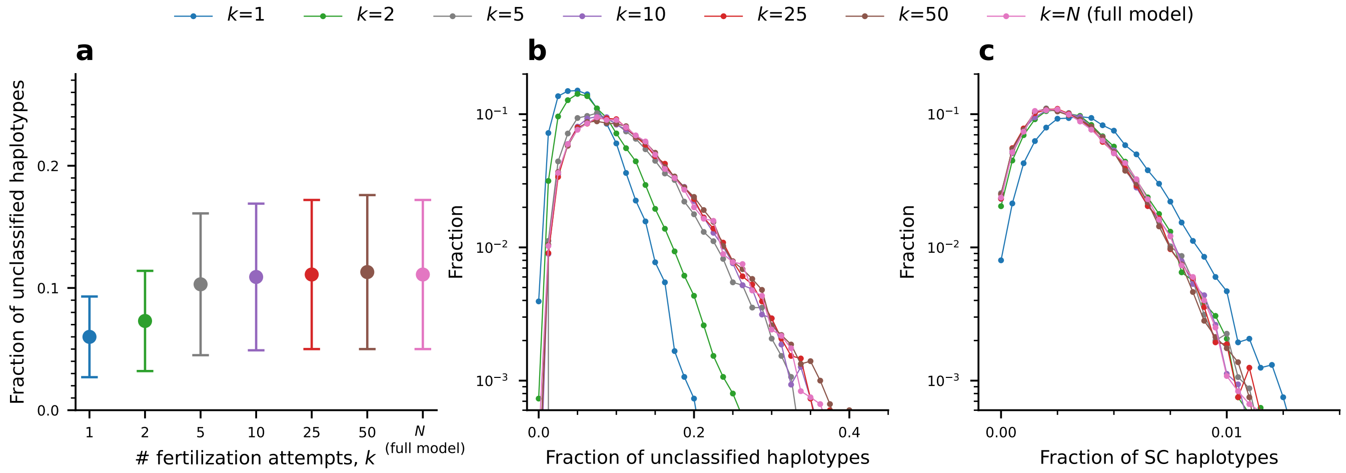

Figure S35: **The population proportion of unclassified S-haplotypes decreases, and the proportion of self-compatible ones increases under pollen limitation**. **(a)** The average population proportion of unclassified S-haplotypes for different numbers of fertilization attempts $k$, denoted by different colors. The bars denote ± STD. **(b-c)** Distribution of the population proportions of unclassified S-haplotypes (b) and self-compatible S-haplotypes (c) for different numbers of fertilization attempts, $k$. We observe that the proportion of unclassified S-haplotypes decreases with $k$, while the proportion of self-compatible ones increases. The figure is based on 16 independent runs for each $k$ value, with 1200 data points per run and a 25-generation interval between consecutive time points used for the analysis. All these data points were taken after the entire population descended from a single common ancestor, and the amino acid frequencies have already reached their steady-state values. The total runtime for each is $2 \times 10^5$ generations. Parameter values: interaction energy thresholds $E_{\text{th}} = -6$. The remaining parameters are as in Table 2 main text, $N = 2000$.

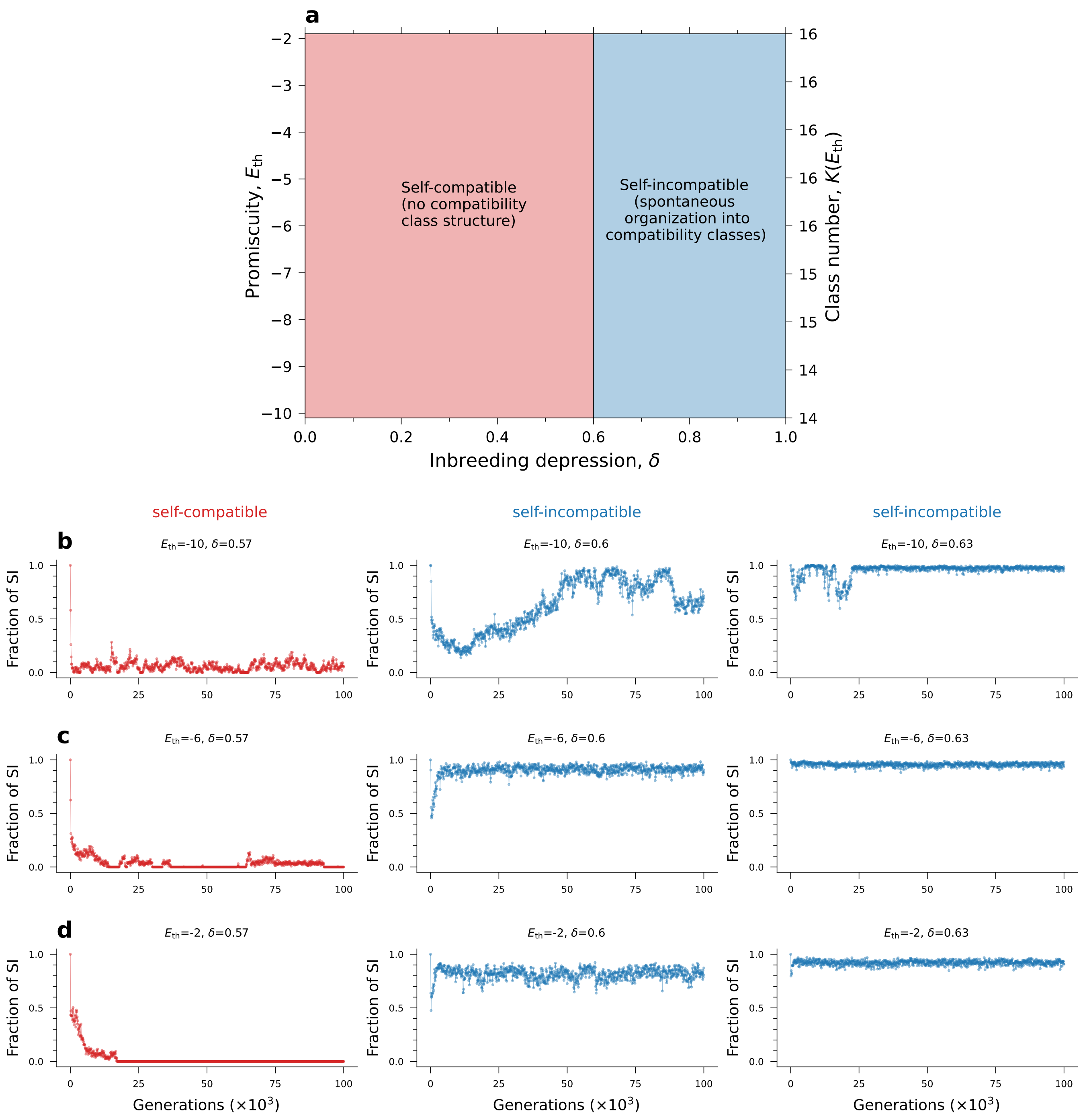

Figure S36: **Both the self-compatibility and self-incompatibility population states exist even under pollen limitation, but the mixed phase nearly vanishes**. We repeated the simulation of the main text, but with only $k = 1$ fertilization attempts per female, to mimic pollen limitation. **(a)** The phase diagram in the entire ($\delta$, $E_{th}$) parameter space. We observe that the mixed unstable phase still exists, but for a very narrow range of $\delta$ values. The calculation of the effective number of classes $K$ (right-side y-axis) refers to $\delta = 0.90$. **(b-d)** Examples of dynamical traces under different parameter combinations. The curve color denotes the population state: self-incompatible (blue), mixed (green), or self-compatible (red). Note that the boundary between the phases is independent of $E_{th}$. The self-pollination rate is $\alpha = 0.60$. Other parameters remain intact (see Table 2, main text, $N = 2000$). The simulation time is $10^5$ generations for each combination of ($E_{th}$, $\delta$).

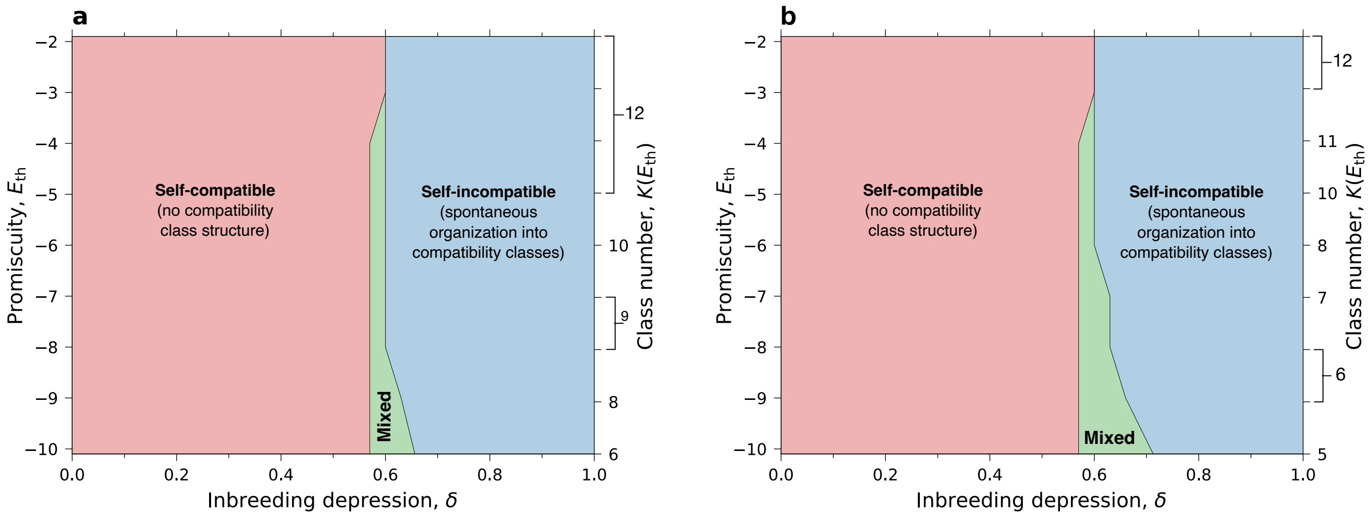

Figure S37: **The mixed phase broadens when the number of fertilization attempts per female $k$ increases.** We repeated the simulation shown in Fig. S36a, with $k = 3$ (**a**), and $k = 5$ (**b**) fertilization attempt per female. As $k$ increases from 1 to 2 and 3, the mixed phase is observed to be wider compared to $k = 1$ (Fig. S36a). The self-pollination rate is $\alpha = 0.60$. The calculation of the effective number of classes $K$ is carried out at $\delta = 0.90$. Other parameters remain intact (see Table 2, main text, $N = 2000$). The number of data points, the independent runs, and the total run-time for each combination of $E_{th}$, and $\delta$ are the same as in Fig. S36.
